# A novel approach to quantify out-of-distribution uncertainty in Neural and Universal Differential Equations

**DOI:** 10.1101/2025.10.01.679743

**Authors:** Stefano Giampiccolo, Giovanni Iacca, Luca Marchetti

## Abstract

Dynamical systems play a central role across the quantitative sciences, offering a powerful mathematical framework to describe, analyze, and predict the evolution of complex processes over time. In systems biology, dynamical systems provide a foundation for modeling and predicting the intricate behaviors of biological systems. Recent advances in data-driven approaches, such as Neural Ordinary Differential Equations (NODEs) and Universal Differential Equations (UDEs), have enabled the development of models that are either fully or partially data-driven. Integrating data-driven components into dynamical systems amplifies the challenge of generalization beyond training data, highlighting the need for robust methods to quantify uncertainty in out-of-distribution (OOD) scenarios—i.e., conditions not encountered during training. In this work, we investigate the reliability of uncertainty quantification (UQ) based on ensembles of models in the reconstruction of dynamical systems. We show that standard ensembles (i.e., models trained independently with different random initializations) risk producing overconfident predictions in previously unseen scenarios, as the models in the ensemble tend to exhibit similar behaviors. To address this issue, we propose a novel ensemble construction method for NODEs and UDEs that fosters diversity in the reconstructed vector field across models within specific regions of the state space, while maintaining explicit control over the fit on the training set. We first evaluate our method on numerical test cases derived from three models commonly used as benchmarks for data-driven reconstruction of dynamical systems: the Lotka–Volterra model, the damped oscillator, and the Lorenz system. We then apply the method to a biologically motivated model of cell apoptosis, considering more realistic conditions such as partial observability of the system outputs and noise in the training dataset. Overall, our results show that the proposed method improves the reliability of UQ in previously unseen scenarios compared with standard ensembles, especially where the latter exhibit overconfidence.

## Introduction

The mathematization of scientific disciplines has led to the widespread use of dynamical systems across a broad spectrum of fields, including systems biology. Systems biology focuses on the quantitative analysis of dynamic interactions among the components of a biological system, with the goal of understanding the behavior of the system as a whole [1]. Within this context, dynamical systems constitute a fundamental mathematical tool, providing a rigorous framework to describe, analyze, and predict the complex behaviors of biological systems over time [2]. By employing dynamical systems, researchers can investigate the interactions among biological entities, uncovering the mechanisms underlying various processes and predicting scenarios that have not been experimentally observed. Mathematically, dynamical systems are codified by systems of Ordinary Differential Equations (ODEs), which express the rate of change of a vector of variables **y** ∈ ℝ^*n*^ with respect to time as a function of **y** and *t*:

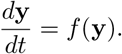

Traditionally, the governing function *f* is *mechanistic* [3], encoding mathematically the theoretical knowledge about the biological system to be modeled. Models synthesized in this way are physically plausible (i.e., they comply with conservation laws and other known governing principles) and are typically interpretable. However, the *mechanistic* approach requires a detailed understanding of the system to be modeled and often relies on strong assumptions that may not be satisfied in real-world scenarios [4].

In recent decades, the growing availability of measured data has fostered the development of *data-driven* approaches to model dynamical systems [5]. Unlike the traditional *mechanistic* approach, these methods do not rely on the physical expertise about the system, but try to infer models directly from observational data. Various methodologies have been proposed, including symbolic regression [6], autoregressive models [7], recurrent neural networks (NNs) [8–10], and Neural Ordinary Differential Equations (NODEs) [11]. Among these, NODEs have emerged as particularly promising, as they retain the continuous-time structure of classical differential equations while substituting the explicit form of the governing function *f* with a NN:

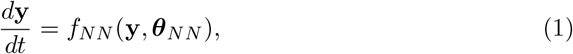

where *f*_*NN*_ represents an NN and ***θ***_*NN*_ the vector of NN parameters. Thanks to the universal approximation theorem for NNs [12], it is theoretically guaranteed that, given sufficient capacity, *f*_*NN*_ can approximate any continuous vector field *f* on a compact subset of ℝ^*n*^—corresponding to the state space of the system—to arbitrary accuracy. Moreover, NODEs can be naturally extended to incorporate partial mechanistic knowledge through the framework of *Universal Differential Equations* (UDEs) [13]. In this formulation, the vector field can be decomposed as

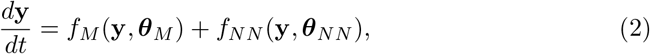

where *f*_*M*_ encodes partially known physical laws or domain-specific insights, ***θ***_*M*_ denotes the (possibly unknown) parameters associated with *f*_*M*_, *f*_*NN*_ is a neural network that models the unknown residual dynamics, and ***θ***_*NN*_ is the corresponding vector of neural network parameters. UDEs and NODEs are increasingly being adopted as modeling frameworks in systems biology, see e.g. [14–17].

Despite the promise shown by NODEs and UDEs, several challenges persist in the data-driven reconstruction of dynamical systems [4]. A central issue is ensuring that models provide reliable information when applied to novel, previously unseen regimes [4, 18]. This issue is not specific to dynamical systems; rather, it reflects a broader limitation in Machine Learning (ML), where model performance often degrades when evaluated on domains that differ from those encountered during training [19]. Formally, the statistical background of ML relies on the assumption that both training and test data are drawn from the same underlying distribution. When this assumption is violated, the test data are considered *out-of-distribution* (OOD) [20], and model predictions typically become significantly less reliable. This represents a major challenge for the applicability of NODEs and UDEs in systems biology, as computational models in this field are often employed to simulate new experiments under previously unseen conditions, predict the outcomes of unobserved biological processes, and generate new testable hypotheses [21]. In such cases, uncertainty quantification (UQ) plays a pivotal role as it allows us to systematically estimate the confidence in model outputs and identify when predictions should—or should not—be trusted [22]. UQ refers to the set of techniques used to measure and represent the uncertainty inherent in model predictions, often by modeling the distribution, variance, or confidence intervals of possible outcomes. Broadly, UQ distinguishes between two types of uncertainty: on one side, aleatoric uncertainty, which arises from the intrinsic randomness in the training data; on the other side, epistemic uncertainty, which stems from limited knowledge of the system to be modeled [23]. The latter cause of uncertainty becomes particularly important when the model is applied in OOD contexts [23]. As such, UQ methods that reliably quantify the epistemic uncertainty in OOD conditions are increasingly recognized as a key requirement in the data-driven reconstruction of dynamical systems [4].

Preliminary studies have begun to explore UQ methods for NODEs and UDEs. Dandekar *et al*. [24] and Ott *et al*. [25] investigate the extension of classical Bayesian techniques [26] to the NODE and UDE frameworks. Takeishi *et al*. [27] propose incorporating regularizers into UQ methods to ensure the identifiability of the mechanistic component in a UDE. Schmid *et al*. [28] compare various Bayesian and ensemble-based approaches for UDEs, offering a formalization of UQ based on maximum likelihood theory. However, none of these studies specifically focus on epistemic uncertainty, as the evaluation of the reliability of the quantified epistemic uncertainty is typically based only on isolated instances of new trajectories (with initial points perturbed relative to the training data) or, in the UDE context, by comparing the reconstructed distribution of the mechanistic component parameter values with the ground truth. To the best of our knowledge, no study has comprehensively investigated the reliability of epistemic uncertainty quantified on the reconstructed vector field of the dynamical system in regions different from the ones encountered during the training or from trajectories predicted by the model using initial states different from those encountered during training.

In this work, we focus on UQ based on standard ensembles (i.e., models trained independently with different random initializations), originally introduced for ML tasks unrelated to dynamical system modeling [29]. We have selected this approach because, despite its simplicity, it has been consistently shown to provide more reliable and practically useful epistemic uncertainty estimates compared to alternative techniques in standard regression and classification machine learning settings [22, 30–32]. Our aim is to investigate the reliability of the epistemic UQ provided by ensemble methods in OOD scenarios in the context of reconstructing dynamical systems using NODEs and UDEs. Consistent with observations in other ML applications [33–35], we find that in OOD dynamical system modeling contexts, often the models of standard ensembles exhibit a similar behavior: this leads to overconfident predictions, with the quantified uncertainty failing to account for the entire epistemic uncertainty in OOD scenarios. To address this challenge, we propose a novel method for constructing ensembles of NODEs and UDEs that fosters diversity in the reconstructed vector field across models in OOD regions of the state space, while maintaining explicit control over the loss function values of the models within the ensemble on the training trajectories. Our method builds on the assumption of *mode connectivity*, which posits that independently trained NODEs/UDEs can be connected via minimum-loss paths in parameter space. Inspired by gradient descent on Riemann manifolds [36], we route these paths to identify models that minimize training loss while maximizing OOD disagreement. Notably, mode connectivity has been validated for standard neural networks [37–39], but it has not yet been studied in the context of NODEs or UDEs. Here, we extend a formal proof of this property to the NODE setting under simplified assumptions, namely full observability of the system, an over-parameterized model, and the availability of densely sampled dynamical system observations, and we discuss its potential validity in the UDE framework.

First, we evaluate our method and compare it to standard ensemble-based UQ using three numerical test cases previously employed in UQ studies for particular types of NODEs [40] and for data-driven dynamical system recovery [11, 13, 41, 42]. These test cases are derived from three dynamical systems: (1) the Lotka–Volterra system, a nonlinear, non-chaotic system with periodic orbits; (2) a damped harmonic oscillator, a linear, non-chaotic system with a fixed-point global attractor; and (3) the Lorenz system, a nonlinear, chaotic system. These classical low-dimensional benchmarks exhibit qualitatively different vector field behaviors, posing varying challenges for data-driven reconstruction. Their low dimensionality also enables clear visualization of the state space, facilitating interpretation of results in terms of distance from the training set. In all test cases, we assume full observability of the system outputs and noiseless training data to isolate and better assess epistemic uncertainty quantification. We then consider a more realistic setting by applying the method to a biologically motivated model of cell apoptosis, where partial observability of the system outputs and noise in the training data are taken into account. In this case, we train the model under one biological condition, namely cell survival, and evaluate the ability of the method to provide meaningful UQ when predicting the system dynamics under a previously unseen condition, namely cell death.

## Results

In this section, we first introduce our proposed method for constructing ensembles of models, explicitly designed to maximize disagreement outside the training domain (Section Proposed method). We then evaluate the performance of our method on a series of numerical test cases, which allow us to rigorously assess the reliability of the uncertainty quantification under controlled conditions and across different levels of incorporated mechanistic prior knowledge (Sections Numerical test cases: benchmark dynamical systems and evaluation metrics, Numerical test cases: full reconstruction and Numerical test cases: partial reconstruction with unknown mechanistic parameters). Finally, we consider a more realistic biological application by analyzing a computational model of cell apoptosis, where we evaluate the uncertainty of a biologically meaningful prediction outside the training scenario (Section Computational biology test case).

### Proposed method

A well-documented limitation of standard ensembles for uncertainty quantification under OOD conditions is their tendency to be overconfident due to *simplicity bias*: models trained with different initializations often converge to similar solutions, exhibiting limited disagreement under OOD conditions [33–35]. This leads to low diversity among ensemble members and, consequently, to underestimated epistemic uncertainty. We provide evidence of this phenomenon in the context of dynamical system reconstruction with NODEs on several numerical test cases (the same used for the benchmark in Section Numerical test cases: benchmark dynamical systems and evaluation metrics) in Supplementary Section S1. To mitigate the risk of overconfidence, we propose a method for training ensembles of dynamical system reconstructions with *Maximized OOD Disagreement* (MOD). The goal of this approach is to induce greater variability in the OOD behavior of the models within the ensemble, resulting in more reliable uncertainty quantification when evaluated on OOD inputs. In the following, we describe the proposed procedure.

We assume here a generic UDE model defined as in Eq. (2); this formulation includes the pure NODE case as a special instance when *f*_*M*_ = **0**. For convenience, supposing that ***θ***_*M*_ ∈ ℝ^*k*^ and ***θ***_*NN*_ ∈ ℝ^*w*^, we denote by ***θ*** the concatenated parameter vector comprising both mechanistic and NN parameters, i.e., ℒ***θ*** = (***θ***_*M*_, ***θ***_*NN*_) ∈ ℝ^*k*+*w*^ and with *g*(***θ*, y**) the RHS of Eq. (2). We further indicate with (***θ***) the loss function that compares the predictions of the UDE with given experimental data. Training a UDE model involves minimizing the function ℒ (***θ***) over ℝ^*k*+*w*^. In general, due to the presence of the overparameterized data-driven component *f*_*NN*_ and the potential non-identifiability of some mechanistic parameters ***θ***^*M*^ [43], the minimum is not uniquely defined, but it is a generic subset

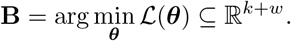

In these terms, assuming for simplicity only epistemic uncertainty, finding an ensemble of *p* models that fit the training set while exhibiting high disagreement under OOD conditions corresponds to finding **b**_**1**_, …, **b**_**p**_ ∈ ℝ^*k*+*w*^ such that

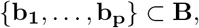

and such that the models corresponding to these parameterizations display high disagreement in OOD conditions (our specific measure of disagreement is detailed later in this section). Our approach is based on the following assumption:

#### Assumption (Mode Connectivity for UDEs and NODEs)

**B** is a connected subset of ℝ^*k*+*w*^, meaning that any two parameterizations achieving minimum loss can be connected by a curve whose points also correspond to minimum-loss parameterizations. More formally, given **b**_1_, **b**_2_ ∈ **B**, there exists a continuous path connecting them that remains entirely within **B**.

This hypothesis has been validated in various settings for standard neural networks [37–39]. To the best of our knowledge, however, it has not yet been formally established for either NODEs or UDEs. In Supplementary Section S2, we extend one of the formal proofs for NNs [44] to the cases of NODEs and UDEs with fixed mechanistic parameters, under hypotheses similar to those used for NNs. Given this property of the loss landscape—namely, that it is possible to move between any two minimum-loss parameterizations along a continuous path—the core idea of our method is to exploit this connectivity. Specifically, starting from a minimum-loss parameterization, we aim to explore the set of parameters achieving minimal loss by following such paths, in order to discover alternative parameterizations that retain low training error while exhibiting markedly different behavior under OOD conditions.

Let *S* be a set of points in the state space outside the trajectories used for training the UDE model, and let **b**_1_, …, **b**_*n*_ ∈ ℝ^*k*+*w*^ denote the parameterizations of an ensemble of models. We assume that their disagreement on *S* can be quantified by a function

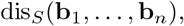

for which the mathematical definition is provided in Section Disagreement measure of reconstructed vector fields. Our approach aims to train an ensemble of models that both achieve the minimum value of the loss function and maximize the disagreement measure dis_*S*_. To this end, we proceed recursively as follows (a visual illustration is provided in Figure 1):

**Fig. 1:**
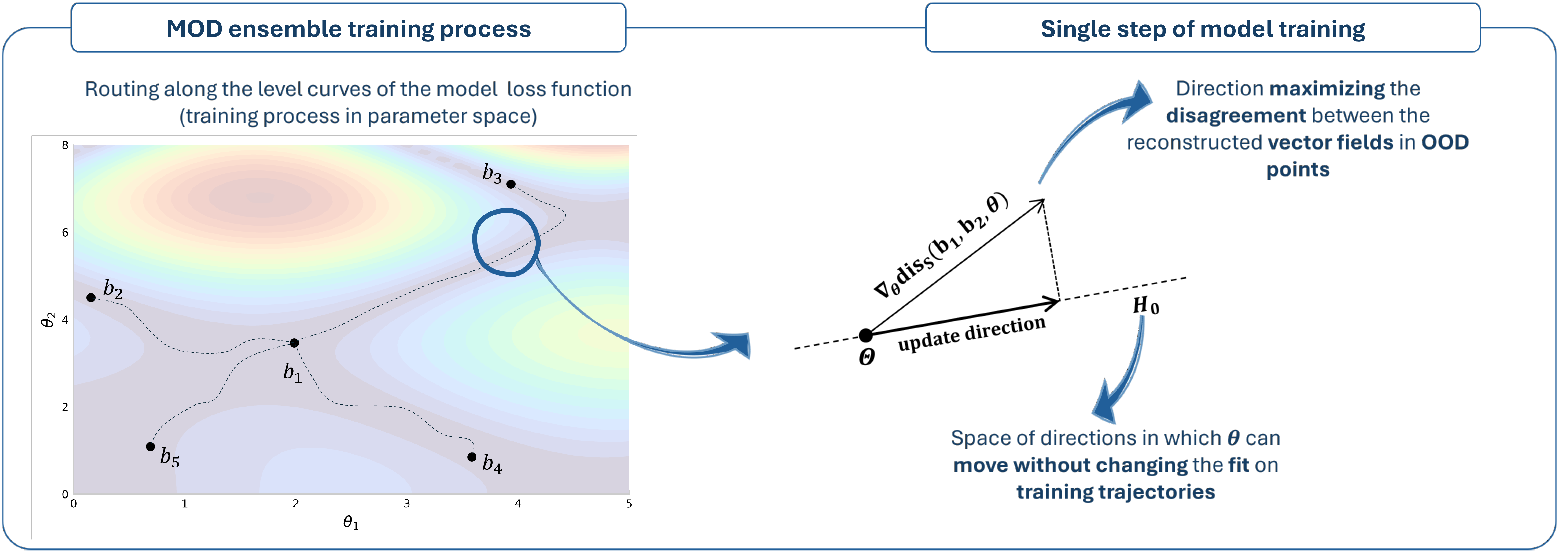
Visual description of the algorithm for training MOD ensembles of UDEs/NODEs. Starting from a base model **b**_**1**_, trained using standard procedures, the algorithm constructs the ensemble sequentially. Each ensemble member is initialized from **b**_**1**_ and trained by following level curves of the loss function, rather than minimizing it directly. The training objective is to maximize disagreement with previously trained models on the reconstructed vector fields, specifically at selected OOD points, thus promoting diversity within the ensemble. Each training process is carried out for a fixed, predefined number of training epochs.

- the first model, **b**_1_, is obtained by minimizing the loss function without considering disagreement;
- then, for each subsequent model **b**_*p*_, with 2 ≤ *p* ≤ *n*, we perform a constrained optimization that maximizes the disagreement dis_*S*_(**b**_1_, …, **b**_*p*_) while ensuring that **b**_*p*_ remains at a minimum of the loss function.

The constraint optimization leverages the mode connectivity hypothesis. Each new model is initialized at ***θ*** = **b**_1_, and the parameters are then updated locally by moving in the direction given by the projection of the gradient of the disagreement function ***θ*** → dis_*S*_(**b**_1_, …, ***θ***) onto the null space *H*_0_ of the Hessian of the loss function ℒ (***θ***), that defines the local directions in the parameter space along which the cost function is not increasing [43] (the constrained optimization step is described in detail in Section MOD ensemble training: constraint optimization step). This procedure is repeated for a fixed, predefined number of epochs. This approach is inspired by constrained optimization on Riemannian manifolds [36], and it is described in detail in Section MOD ensemble training: how to route loss contours. The complete MOD ensemble training procedure is described in Section MOD ensemble training: overall algorithm description and illustrated in Algorithm 1.

### Numerical test cases: benchmark dynamical systems and evaluation metrics

To evaluate the effectiveness of our algorithm, we consider three distinct dynamical systems: the Lotka–Volterra predator–prey model [45], the damped oscillator [42], and the Lorenz system [46]. For each system, we generated an *in silico* training dataset by discretely sampling multiple trajectories obtained through numerical simulation of the corresponding ODE system. Ground-truth dynamical systems are described in Section Test cases, while the corresponding literature parameters and training trajectories are reported in Supplementary Section S3. To isolate and better analyze epistemic uncertainty, no additional aleatoric noise was added to the training trajectories in these test cases.

We assess how an ensemble of models reconstructing the dynamical system quantifies uncertainty on OOD inputs, focusing on two outputs: the vector field and the trajectories. The vector field represents the predicted derivatives across the state space, while trajectories describe the system’s time evolution. Although trajectories are the primary quantities of biological interest, analyzing the vector field helps visualize how uncertainty evolves as one moves away from the training data. We consider OOD vector field predictions as those made outside the region of the state space covered by training trajectories, and OOD trajectory predictions as the predictions obtained with a model from new initial conditions not seen during training.

To rigorously assess the reliability of uncertainty quantification, we derive 0.95 prediction intervals from the ensembles and analyze their coverage properties with respect to the ground truth, which is known in these numerical test cases. A 0.95 prediction interval defines a range within which the true output is expected to lie with probability 0.95. Mathematical details on the construction of these intervals for vector-field and trajectory predictions are provided in Sections Prediction intervals on vector field and Prediction intervals on trajectories, respectively. Coverage indicates whether the ground-truth value lies within the prediction interval for a given evaluation input. Given a set of OOD state-space points and a set of OOD trajectories, the coverage proportion (CP) is defined as the fraction of evaluation inputs for which the prediction intervals cover the ground truth. Ideally, the CP should approach the nominal level of 0.95 [47]. Values below 0.95 indicate overconfidence, meaning that uncertainty is underestimated, whereas values above 0.95 indicate underconfidence, corresponding to conservative intervals. Since training NODE/UDE models involves stochastic optimization, multiple ensembles are generated. For each OOD input, we compute the mean coverage across ensembles, and then compute the mean CP over the set of OOD inputs. This value is compared to the nominal level of 0.95. Further details on the OOD input sets considered for vector fields and trajectories are provided in Section OOD input sets, while a graphical description is reported in Supplementary Section S4.

For each dynamical system, we consider two reconstruction scenarios corresponding to different levels of prior mechanistic knowledge:

- **full reconstruction**: no prior knowledge is assumed, and the system is reconstructed purely from data (Section Numerical test cases: full reconstruction);
- **partial reconstruction with unknown mechanistic parameters**: partial structural knowledge is assumed, and both mechanistic and NN parameters are estimated from data (Section Numerical test cases: partial reconstruction with unknown mechanistic parameters).

A third scenario, partial reconstruction with known mechanistic parameters, is reported in Supplementary Section S5.

For each scenario, we compare the results of standard and MOD ensembles. Details of this comparison are provided in Section Comparison between MOD and standard ensembles. In these numerical experiments, we assume prior knowledge of the region where disagreement should be maximized. Accordingly, we define the set *S*, on which the MOD algorithm maximizes disagreement, as a regular grid of state-space points where prediction interval coverage of the vector field is evaluated. Additional implementation details for the MOD algorithm are given in Section Algorithmic setup, while the procedure for training standard ensembles is described in Section Standard ensemble training; the corresponding hyperparameters are reported in Supplementary Section S6.

### Numerical test cases: full reconstruction

In the first scenario, we consider a purely data-driven setting, where all models are implemented as pure NODEs. The architecture of the NN consists of an input layer, two hidden layers with 32 nodes each using the *gelu* activation function, and an output layer, where the input and output dimensions depend on the dimensionality of the dynamical system. For each test case, we train 10 standard ensembles and 10 MOD ensembles. The training fit of all 50 NODEs comprising the standard ensembles, as well as the 50 NODEs in the MOD ensembles, is analyzed in Supplementary Section S7.

We begin by analyzing the coverage of 0.95-prediction intervals on the vector field obtained with the two approaches (standard and MOD ensembles). The CP results are reported in Table 1, whereas the mean coverage heatmaps in the considered regions of the state space are shown in Figure 2.

**Table 1:** Comparison of the CP of 0.95-prediction intervals on the vector field (full reconstruction scenario). Mean CP of 0.95-prediction intervals on the vector field within a selected region of the state space. The results are presented as the mean *±* SEM across 10 ensembles. Statistical significance was assessed using the Wilcoxon signed-rank test on absolute deviations from 0.95 between the two distributions of CP values. Asterisks indicate the following significance levels: * p ≤ 0.05, ** p ≤ 0.01, and *** p ≤ 0.001. The values shown in bold are those closest to the theoretical value.

|  | **Lotka–Volterra** | **Damped Oscillator** | **Lorenz** |
| --- | --- | --- | --- |
| Standard | 0.850 ± 0.024 | 0.669 ± 0.016 | 0.458 ± 0.010 |
| MOD | **0.986 ± 0.006 (\*)** | **0.993 ± 0.003 (\*\*)** | **0.760 ± 0.040 (\*\*)** |

**Fig. 2:**
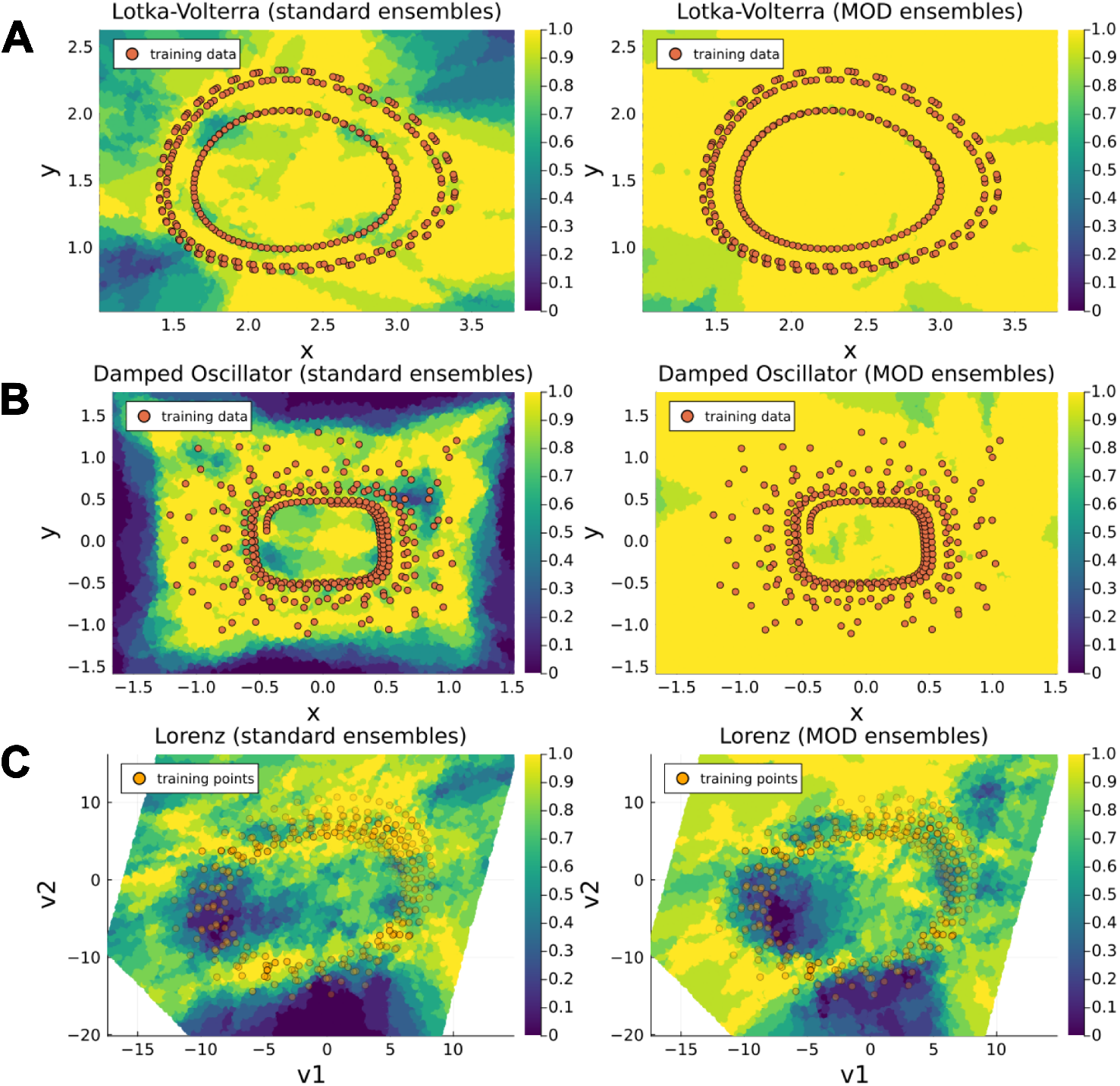
Comparison of 0.95-prediction interval coverage on state space: Standard vs. MOD Ensembles (full reconstruction scenario). Panels A, B, and C display heatmaps of the mean coverage (computed across 10 ensembles) of 0.95-prediction intervals on the state space for the Lotka–Volterra, Damped Oscillator, and Lorenz systems, respectively. For each system, the left subpanel shows results obtained with standard ensembles, while the right subpanel shows results obtained with MOD ensembles. Orange overlays represent the points from the training trajectories, providing spatial context. For the Lorenz system, the mean coverage is visualized on a two-dimensional plane spanned by the coordinates (*v*_1_, *v*_2_), obtained via linear regression on the Lorenz attractor (see Supplementary Section S8 for details). These coordinates provide a low-dimensional representation of the original three-dimensional state space (*x, y, z*); here, the transparency of training trajectory points indicates their orthogonal distance from the training trajectory points to the plane.

The results indicate that MOD ensembles yield mean CP values significantly closer to the nominal value of 0.95 than standard ensembles in all test cases (one-sided Wilcoxon signed-rank test, *p <* 0.01). In the Lotka–Volterra test case, standard ensembles produce prediction intervals with a mean CP close to the nominal value, which becomes even closer when MOD ensembles are applied. In the Damped Oscillator and Lorenz test cases, the prediction intervals produced by standard ensembles are distinctly overconfident, and this problem is mitigated by applying MOD ensembles. In the Damped Oscillator test case, the mean CP obtained with MOD ensembles reaches a value slightly above the nominal value, and the mean coverage heatmaps show that our approach successfully mitigates the overconfidence exhibited by standard ensembles across entire regions of the state space. In the chaotic Lorenz system, although the mean CP obtained with MOD ensembles remains below the nominal value, it is still closer to the nominal value than that achieved by standard ensembles.

The results for the coverage of the prediction intervals on trajectories are consistent with those observed on the vector field and are summarized in Table 2. In the Lotka–Volterra system, the improvement brought by MOD ensembles is statistically less pronounced, as standard ensembles already yield mean CP values relatively close to the nominal value of 0.95. In contrast, for the Damped Oscillator and Lorenz systems, the prediction intervals obtained with MOD ensembles produce mean CP values significantly closer to the nominal value than those from standard ensembles. An example illustrating how our algorithm improves the coverage for the trajectories is shown in Figure 3.

**Table 2:** Comparison of the CP of 0.95-prediction intervals on the trajectories (full reconstruction scenario). Mean CP of 0.95-prediction intervals on trajectories on the selected points of test trajectories. The results are presented as the mean *±* SEM across 10 ensembles. Statistical significance was assessed using the Wilcoxon signed-rank test on absolute deviations from 0.95 between the two distributions of CP values. Asterisks indicate the following significance levels: * p ≤ 0.05, ** p ≤ 0.01, and *** p ≤ 0.001. The values shown in bold are those closest to the theoretical value.

|  | **Lotka–Volterra** | **Damped Oscillator** | **Lorenz** |
| --- | --- | --- | --- |
| Standard | $0.800 \pm 0.042$ | $0.627 \pm 0.016$ | $0.546 \pm 0.030$ |
| MOD | **$0.926 \pm 0.019$ (\*)** | **$0.954 \pm 0.012$ (\*\*)** | **$0.738 \pm 0.030$ (\*\*)** |

**Fig. 3:**
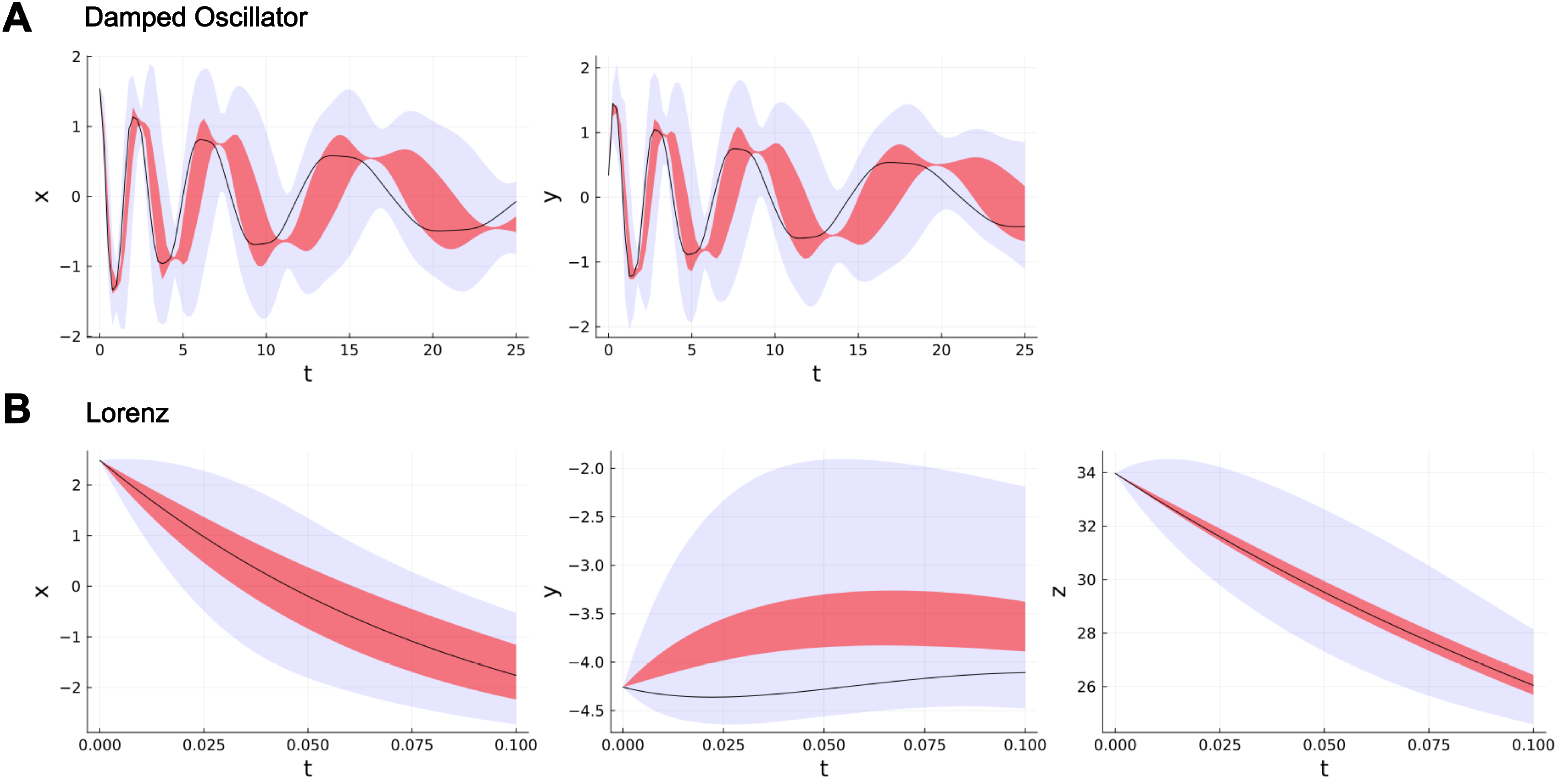
Comparison between prediction interval on trajectories obtained with a standard ensemble and a corresponding MOD ensemble (full reconstruction scenario). Panels A and B illustrate two trajectories for which the trained standard ensembles produce overconfident 0.95-prediction intervals in the Damped Oscillator and Lorenz test cases, respectively, whereas the corresponding MOD ensemble covers the ground truth. The shaded red regions denote the 0.95-prediction intervals obtained with the standard ensemble, the shaded blue regions denote the 0.95-prediction intervals obtained with the MOD ensemble, and the solid black line represents the corresponding ground-truth trajectory. Additional comparisons on further sampled trajectories are presented in Supplementary Section S10.

We note that, in some cases, the empirical CP distributions obtained with MOD ensembles—both for the vector field and the trajectory reconstructions—exceed the nominal coverage level of 0.95. This behavior may indicate a degree of underconfidence in the corresponding prediction intervals. The potential causes of this phenomenon are discussed in detail in Section Discussion.

We conclude this section by noting that the results do not depend on the specific training dataset. As shown in Supplementary Section S9, MOD ensembles trained on different datasets yield consistent outcomes, underscoring the generality of our approach.

### Numerical test cases: partial reconstruction with unknown mechanistic parameters

In this second scenario, we assume partial knowledge of the dynamical system, with only a portion of the system derivatives requiring reconstruction through data-driven methods. The specific portions assumed to be mechanistically known are reported in Section Test cases. We do not assume prior knowledge of the parameters of the known mechanistic portions of the model. Instead, we estimate these parameters alongside the NN parameters. The mechanistic parameters were initialized within a boundary range between 0.5 and 2 times the ground truth values (reported in Supplementary Section S3). The architecture used for the NN is identical to the one used in the first scenario. We train 10 standard ensembles and 10 MOD ensembles. The fit results on the training trajectories are reported in Supplementary Section S11.

The results for the coverage of the 0.95 prediction intervals on vector fields for standard ensembles and MOD ensembles, in terms of mean CP and mean coverage, are presented in Table 3 and Figure 4, respectively. In the Lotka–Volterra system, the prediction intervals generated by standard ensembles are not overconfident, and, in this case, MOD ensembles lead to a mean CP slightly above the nominal value, although in both cases the deviation from 0.95 is less than 0.05. In the Damped Oscillator system, standard ensembles exhibit distinct regions where the prediction intervals are overconfident, resulting in a mean CP of 0.597, while MOD ensembles increase the mean CP to 0.997, bringing it distinctly closer to the nominal value. In the Lorenz system, the prediction intervals from standard ensembles exhibit relatively high coverage compared to the results obtained for the same system in the first scenario, and the mean CP obtained with MOD ensembles is even closer to 0.95, although the improvement in CP between the model pairs is not statistically significant.

**Table 3:** Comparison of the CP of 0.95-prediction intervals on the vector field (partial reconstruction with unknown mechanistic parameters scenario). Mean CP of 0.95-prediction intervals on the vector field within a selected region of the state space. The results are presented as the mean *±* SEM across 10 ensembles. Statistical significance was assessed using the Wilcoxon signed-rank test on absolute deviations from 0.95 between the two distributions of CP values. Asterisks indicate the following significance levels: * p ≤ 0.05, ** p ≤ 0.01, and *** p ≤ 0.001. The values shown in bold are those closest to the theoretical value.

|  | **Lotka–Volterra** | **Damped Oscillator** | **Lorenz** |
| --- | --- | --- | --- |
| Standard | **0.955 ± 0.015** (**) | 0.597 ± 0.014 | 0.811 ± 0.054 |
| MOD | 0.999 ± 0.001 | **0.997 ± 0.002** (**) | **0.841 ± 0.055** |

**Fig. 4:**
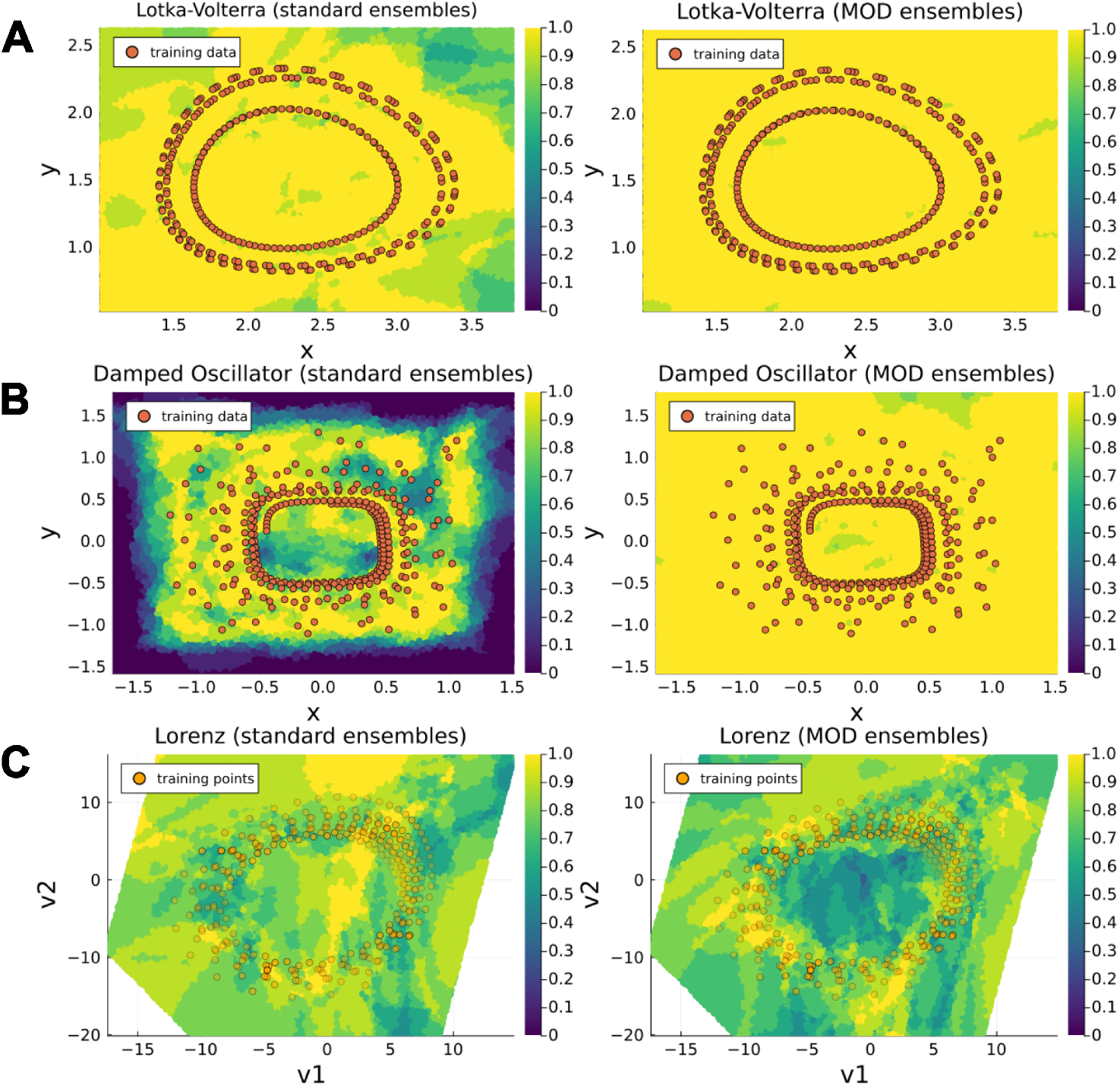
Comparison of 0.95-prediction interval coverage on state space: Standard vs. MOD Ensembles (partial reconstruction with unknown mechanistic parameters scenario). Panels A, B, and C display heatmaps of the mean coverage (computed across 10 ensembles) of 0.95-prediction intervals on the state space for the Lotka–Volterra, Damped Oscillator, and Lorenz systems, respectively. For each system, the left subpanel shows results obtained with standard ensembles, while the right subpanel shows results obtained with MOD ensembles. Orange overlays represent the points from the training trajectories, providing spatial context. For the Lorenz system, the mean coverage is visualized on a two-dimensional plane spanned by the coordinates (*v*_1_, *v*_2_), obtained via linear regression on the Lorenz attractor (see Supplementary Section S8 for details). These coordinates provide a low-dimensional representation of the original three-dimensional state space (*x, y, z*); here, the transparency of training trajectory points indicates their orthogonal distance from the training trajectory points to the plane.

The results for the CP of the 0.95 prediction intervals on trajectories (Table 4) are consistent with those observed on the vector field. In particular, for the Damped Oscillator and Lorenz systems, MOD ensembles mitigate the overconfidence present in the prediction intervals generated by standard ensembles. The improvement is especially pronounced in the Damped Oscillator test case, where the mean CP from standard ensembles is particularly low and is increased to above 0.90 by MOD ensembles.

**Table 4:** Comparison of the CP of 0.95-prediction intervals on the trajectories (partial reconstruction with unknown mechanistic parameters scenario). Mean CP of 0.95-prediction intervals on trajectories on the selected points of test trajectories. The results are presented as the mean *±* SEM across 10 ensembles. Statistical significance was assessed using the Wilcoxon signed-rank test on absolute deviations from 0.95 between the two distributions of CP values. Asterisks indicate the following significance levels: * p ≤ 0.05, ** p ≤ 0.01, and *** p ≤ 0.001. The values shown in bold are those closest to the theoretical value.

|  | **Lotka–Volterra** | **Damped Oscillator** | **Lorenz** |
| --- | --- | --- | --- |
| Standard | **0.947 ± 0.011** (**) | 0.440 ± 0.038 | 0.813 ± 0.047 |
| MOD | 0.987 ± 0.001 | **0.905 ± 0.026** (**) | **0.945 ± 0.019** |

The exploration of the parameter space performed by our algorithm is illustrated by the trajectories of the mechanistic parameters during MOD ensemble training and is analyzed in Supplementary Section S12, where the implications of mechanistic parameter identifiability are also discussed.

### Computational biology test case

In this section, we extend our analysis to a more biologically realistic scenario by considering a higher-dimensional computational biology model than those examined previously. We assume partial observability of the system dynamics and the presence of measurement noise, enabling us to evaluate the performance of the MOD training algorithm under conditions that more closely reflect real-world biological applications.

The model under consideration describes cell apoptosis [48], a core sub-network of the signal transduction cascade regulating the programmed cell death–versus–survival phenotypic decision. This model has previously been adopted as a benchmark to assess the reliability of scientific machine learning approaches [43, 49]. It is well known to exhibit stiffness (with literature parameter values) and to be characterized by both structural and practical parameter non-identifiability. It describes the temporal evolution of 8 different species: the model equations are reported in Section Test cases, while the original parameters’ values are reported in Supplementary Section S3. This system can reproduce two different scenarios, corresponding to cell survival and cell death [49], by modifying the initial conditions of the variable *y*_7_ (the different initial conditions are reported in Section Test cases).

Following what already done in [49], in our experimental setting we assume that only the variable *y*_4_ is observable. The training dataset was generated under the cell-survival scenario: Figure 5 shows the training set in the cell-survival scenario and the ground truth dynamics of the observable variable *y*_4_ in the cell-death scenario.

**Fig. 5:**
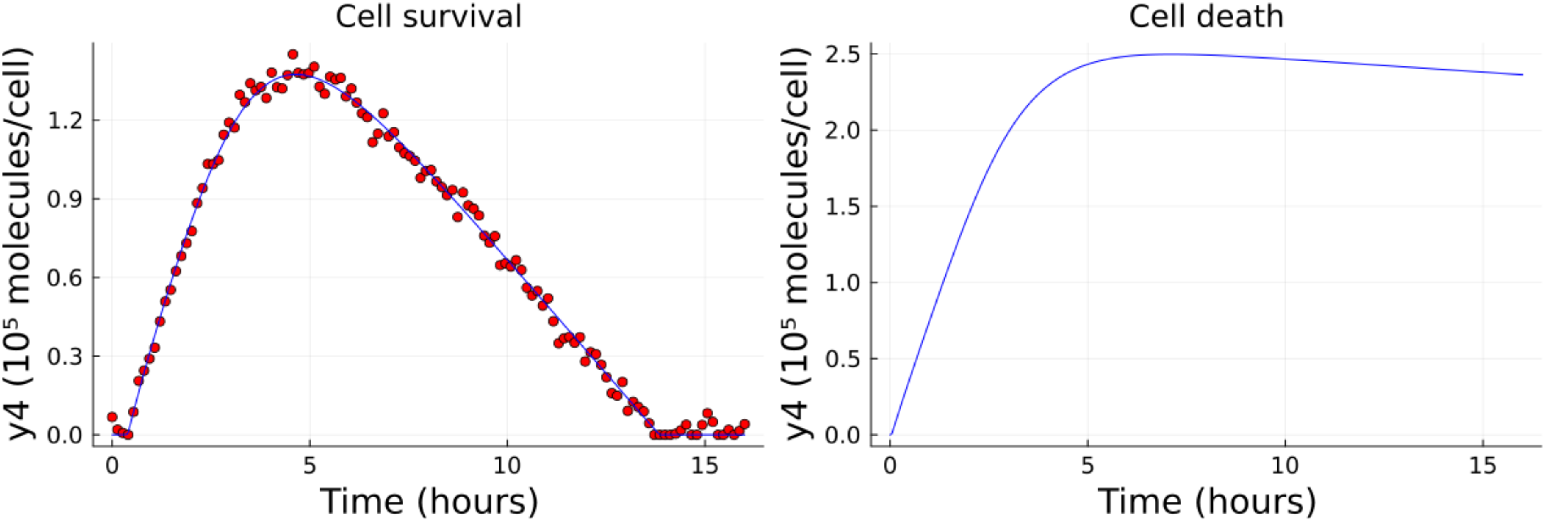
Training data and ground-truth dynamics of *y*_4_ in the cell-survival and cell-death scenarios. On the left, the ground-truth dynamics of *y*_4_ in the cell-survival scenario (blue line) together with the training data (red points). On the right, the ground-truth dynamics of the same variable in the cell-death scenario.

We assume no prior knowledge of the mechanistic equation governing *y*_4_. Consequently, this component is replaced by a neural network (the architecture of the NN consists of an input layer, two hidden layers with 32 nodes each using the *gelu* activation function, and an output layer). None of the mechanistic parameters is assumed to be known *a priori*, placing the problem in a partial reconstruction setting. The model is trained exclusively on data generated under the cell-survival scenario; the objective is to derive a reliable UQ that can be propagated when predicting system behavior in the cell-death scenario. To this end, as in the previous numerical test cases, we trained 10 standard ensembles, each composed of 5 different models. Subsequently, we trained a MOD ensemble by initializing it from one randomly selected model from each standard ensemble. We then compared the resulting predictions in terms of coverage of the ground-truth dynamics of the variable *y*_4_ in the cell-death scenario. Further details regarding the training of both the standard ensembles and the MOD ensemble—including the selection of the region of the state space in which the disagreement among models is maximized—are reported in Section Algorithmic setup. The fits of the models composing the standard and MOD ensembles are reported in Supplementary Section S13.

When evaluating the 0.95 prediction intervals on the ground-truth dynamics of *y*_4_ under cell-death conditions, we find that, on average, the intervals obtained from the standard ensembles contain only 62.5% of the ground-truth trajectory. Furthermore, half of the standard ensembles are highly overconfident, with prediction intervals covering less than 40% of the trajectory (see Figure 6). In these cases, the prediction intervals tend to pull the dynamics of *y*_4_ in this new scenario back toward the dynamics observed during training (see the yellow area in Figure 6 for an example involving the first two ensemble pairs; results for all ensemble pairs are reported in Supplementary Section S14). In contrast, the prediction intervals obtained from the MOD ensembles show a markedly different behavior. On average, they cover 96% of the ground-truth trajectory, which is close to the nominal level of 0.95 (see the green area in Figure 6 for an example involving the first two ensemble pairs; results for all ensemble pairs are reported in Supplementary Section S14). Moreover, all MOD ensembles except one contain more than 98% of the ground-truth trajectory, improving the overconfidence of the standard ensembles.

**Fig. 6:**
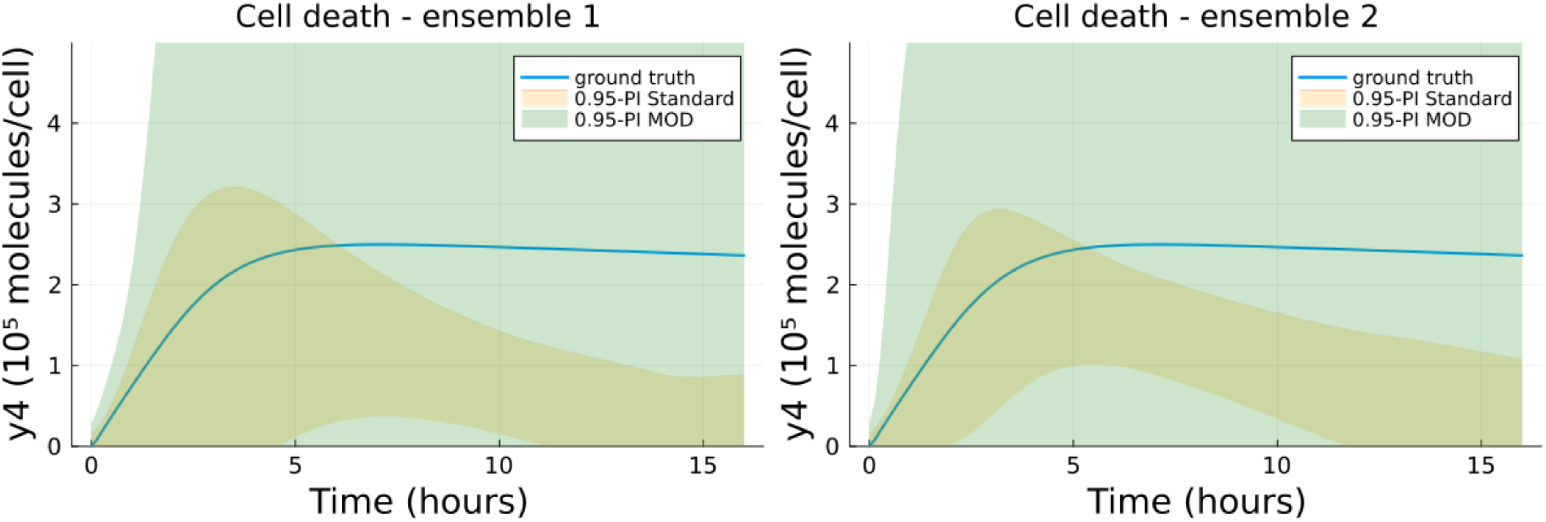
0.95-prediction intervals for *y*_4_ in the cell-death scenario. 0.95-prediction intervals obtained with standard ensembles (yellow) and MOD ensembles (green) for two representative ensemble pairs. The blue line represents the dynamics of the ground-truth model for the variable *y*_4_.

The prediction intervals derived from the MOD ensembles are very wide, indicating that, given the current dataset and the structure of the hybrid model, no reliable prediction can be made for the cell-death scenario. While this may appear as a limitation, it is in fact an informative outcome, as it explicitly reveals the limits of the model’s predictive capabilities. In the terminology of systems biology, this implies that the prediction of *y*_4_ under cell-death conditions does not constitute a *core prediction* of the model [50]. To show it, we selected, from the trajectories in parameter space explored by the MOD ensemble algorithm, a set of example parameterizations achieving a training cost lower than 0.0015 (approximately the midpoint of the costs obtained with the standard ensembles). Despite providing fits comparable to those of the standard ensembles, these parameterizations display markedly different predictions for the dynamics of *y*_4_ in the cell-death scenario. The corresponding fits on the training set and the resulting dynamics in the cell-survival scenario are reported in Figure 7.

**Fig. 7:**
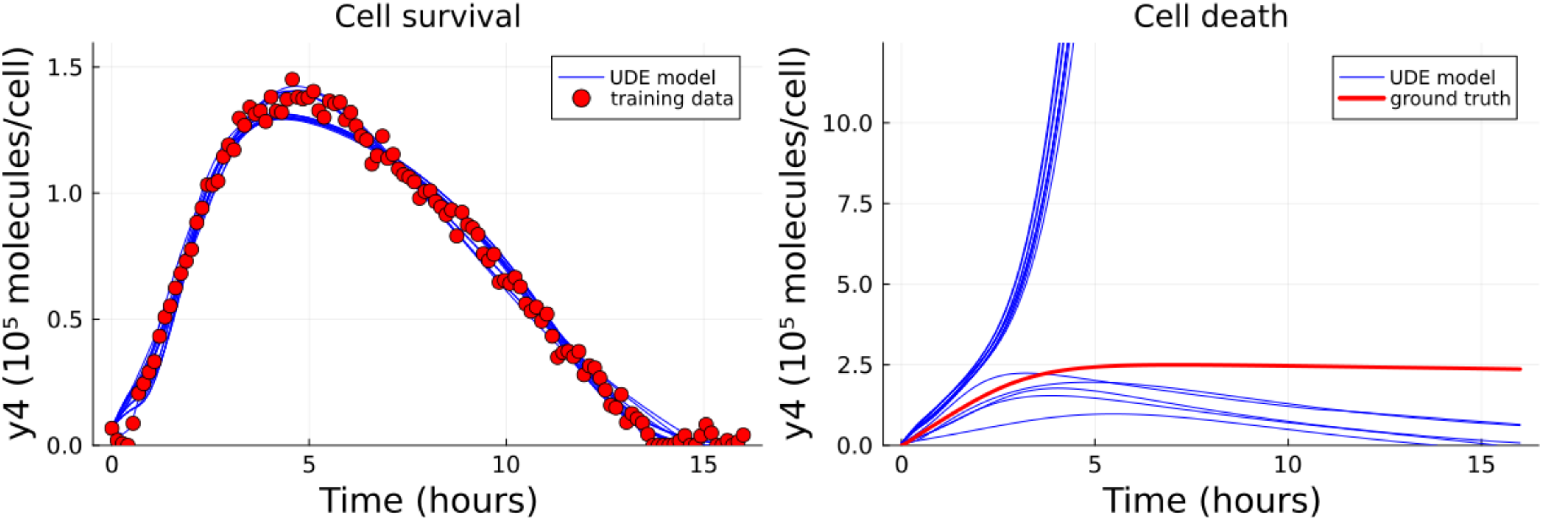
Fit of the models with the sampled parameterizations on the training set and corresponding predictions in the cell-death scenario. On the left, the simulations of *y*_4_ obtained with the sampled parameterizations under the training (cell-survival) scenario (blue lines), together with the training data (red points). On the right, the simulations of *y*_4_ obtained with the same parameterizations under the cell-death scenario (blue lines), together with the ground-truth dynamics of *y*_4_ (red line).

## Discussion

In this work, we address the challenge of uncertainty quantification (UQ) in the data-driven reconstruction of dynamical systems using neural ordinary differential equations (NODEs) and universal differential equations (UDEs). We consider two main outputs of the reconstruction process: the reconstructed vector fields and the prediction of temporal dynamics not observed during training. In particular, we investigate the problem of estimating the uncertainty both in out-of-distribution (OOD) regions of the state space and along previously unseen trajectories. To address the risk of overconfidence of standard ensembles, we introduce a novel training strategy that promotes diversity among ensemble members by maximizing their disagreement in selected OOD regions of the state space while maintaining a controlled training error. The proposed approach is first evaluated on three *in silico* numerical test cases, where we analyze both the reconstruction of the vector field and the prediction of unseen trajectories under different levels of prior physical knowledge about the system (a comparison of algorithm performance across different levels of prior physical knowledge is provided in Supplementary Section S15). We then consider a more challenging biological benchmark, the cell apoptosis model, where the focus is on predicting system dynamics under cell-death conditions, while the training dataset corresponds to the cell-survival condition and includes partial observability and noise. Across all these test cases, we show that the proposed MOD ensembles provide more reliable uncertainty estimates than standard ensembles, particularly in scenarios where the latter exhibit overconfidence. Within a single scenario of the numerical test cases, we compare our method against the Laplace approximation, highlighting its effectiveness relative to alternative uncertainty quantification approaches beyond standard ensembles (Supplementary Section S16).

The main limitations of this work are: (a) the occurrence of rare cases in which the mechanistic component of the model may violate the single-mode connectivity assumption; (b) the potential risk of underconfident prediction intervals; (c) the dependence on the chosen region of the state space, along with the lack of a clear stopping criterion for terminating the training of each model; and (d) the choice of the hyper-parameter used to distinguish directions in which the cost function does not increase. These aspects are examined in more detail in the following paragraphs.

The assumption of single-mode connectivity in UDEs fails when the mechanistic component of the model contains parameters that are locally identifiable but not globally identifiable. In such instances, the loss function may exhibit minima that are not connected. The rarity and the causes of this phenomenon have been explored in the context of biological mechanistic modeling by Barreiro *et al*. [51], who investigated its occurrence across a range of published models. They identified specific conditions for this phenomenon, such as parameters appearing with even exponents, which can admit both positive and negative values. In these cases, the single-mode connectivity assumption should be replaced with the hypothesis that the set of global minima in the UDE consists of several connected components. This would necessitate an adaptation of the MOD training algorithm, requiring initialization from multiple, traditionally trained parameterizations of the model, each located in a distinct connected component. Extending the methodology to address such (rare) cases is beyond the scope of the present study.

The prediction intervals obtained with both standard ensembles and the MOD approach may be affected by underconfidence (i.e., overly broad intervals). In our setting, synthetic data cannot be used to determine a “correct” width for epistemic uncertainty bands. Validating interval width would require knowledge of the full distribution of parameterizations consistent with the data, which is not available in practice. Nonetheless, the broader intervals observed with MOD do not arise from artificially inflating variance. Instead, the method constructs alternative parameterizations that fit the training data equally well while diverging in weakly constrained regions. By routing along low-loss contours and maximizing out-of-distribution disagreement, MOD reveals model variability already compatible with the data rather than introducing variability artificially. Potential sources of underconfidence may arise at two stages of the procedure. First, as the MOD training algorithm explores the parameter space, it may accumulate increases in the cost function, which are controlled by a predefined threshold. If this threshold is too permissive, suboptimal fits may inflate predictive variance; if too restrictive, plausible solutions may be excluded. We treat this threshold as a modeling choice: in the numerical test cases, we set it to 10^−3^, and we assess the robustness of the algorithm under a stricter threshold in the full reconstruction scenario of the numerical test cases (Supplementary Section S17). A detailed discussion on why the threshold is treated as a modeler choice is provided in Supplementary Section S18. Second, prediction intervals are derived under a Gaussian (or multivariate Gaussian) assumption over ensemble predictions. While this is a standard approximation in ensemble-based uncertainty estimation [29], it remains an assumption and may imperfectly reflect the true predictive distribution.

A further major limitation of the present work concerns the user-defined selection of the region of the state space in which model disagreement is maximized, as well as the absence of an adaptive stopping criterion during the routing of individual ensemble members. The MOD algorithm is designed to target a specific region of interest. In practical applications, when a scientist seeks to quantify uncertainty in a given model prediction, a region of the state space comprising that prediction must first be defined, and the algorithm is then applied within that region. Our results demonstrate that maximizing disagreement in such a region increases ensemble diversity and improves prediction interval coverage relative to standard ensembles, including over broader portions of the vector field (Supplementary Sections S19 and S20). Nevertheless, uncertainty quantification remains inherently sensitive to the chosen region: a MOD ensemble optimized for one portion of the state space may not be optimal for substantially different regions. In this sense, the method is goal-oriented rather than globally uniform, and careful definition of the region of interest constitutes an important modeling decision. As illustrated in the biological test case, one possible strategy is to first perform the prediction of interest using the starting model of the MOD ensemble and then derive the relevant region of the state space from that prediction. Furthermore, in our test cases, the ensemble models were trained for a fixed number of iterations without a convergence criterion or a threshold to stop the training. An analysis of the maximization curve for model disagreement during training is provided in Supplementary Section S21, which reveals key challenges in defining a threshold on the objective function or a convergence criterion. First, the values assumed by the objective function are dependent on the system considered and on the OOD region in which the disagreement is maximized, thus preventing the introduction of a fixed threshold to stop the training of models in MOD ensembles. Second, no clear maximum in model disagreement may exist, leading to a failure in optimization convergence. Nevertheless, as discussed in Supplementary Section S22 for the Lorenz test case in the full reconstruction scenario, a fixed and limited number of iterations may lead to underconfidence in the prediction intervals.

Finally, a limitation of the proposed method concerns the hyperparameter *ϵ*_*eig*_ used to identify the null space of the Hessian matrix. When the Hessian is computed numerically, a positive threshold is required to distinguish directions along which the cost function increases from those along which it remains approximately constant. The choice of this threshold is arbitrary; the value used for the test cases considered in this study is discussed in Supplementary Section S23.

Despite these limitations, the proposed work represents an initial step toward extending traditional UQ methods to address OOD UQ in dynamical system reconstruction. In future work, we aim to extend the method in two key directions. First, we will focus on incorporating the qualitative characteristics of the vector field into the ensemble training process. The analysis presented in Supplementary Section S24 shows that our training algorithm currently produces models that fail to respect key qualitative features of the vector field, such as periodicity in the Lotka–Volterra case or the existence of a global attractor in the Damped Oscillator case—features that a modeler could inductively deduce. To address this, we plan to introduce these characteristics as soft constraints in the loss function. Finally, we aim to address the sequential nature of our algorithm by introducing parallelization in the training process of the models composing the MOD ensemble. Specifically, we plan to train the ensemble models simultaneously, maximizing the disagreement between them during training.

## Methods

In this section, we present the methods used to obtain the results reported in this work. Specifically, Section Test cases describes the test cases. Sections Prediction intervals on vector field and Prediction intervals on trajectories detail the methods used to derive the 0.95 prediction intervals for the vector field and trajectories, respectively. Section OOD input sets describes the OOD input sets used to assess the reliability of the uncertainty quantification. Section Standard ensemble training explains the training procedure for standard ensembles. Sections Disagreement measure of reconstructed vector fields, MOD ensemble training: constraint optimization step, MOD ensemble training: how to route loss contours, and MOD ensemble training: overall algorithm description concern the MOD ensemble training and describe, respectively, the disagreement measure among reconstructed vector fields, the method for performing a local training step, the overall constraint optimization procedure, and the complete MOD ensemble training algorithm. Section Comparison between MOD and standard ensembles outlines the comparison between MOD and standard ensembles; and Section Algorithmic setup reports implementation details.

### Test cases

In the following, we report the equations of the models used as benchmarks in this work, including both the numerical and biological test cases. For the numerical test cases, three training trajectories per system were generated by numerically integrating the governing equations from different initial conditions.

The first system is the Lotka–Volterra predator–prey model, described by the following two-dimensional ODE system:

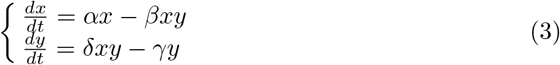

In the partial reconstruction scenario, we assume no knowledge of the interaction terms between the two species, and we replace them with an NN, leading to the following formulation:

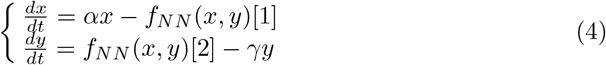

where *f*_*NN*_ : ℝ^2^ → ℝ^2^ is an NN that takes (*x, y*) as input and outputs the approximated interaction terms.

The second system is a nonlinear damped oscillator [40], defined by the two-dimensional ODE system:

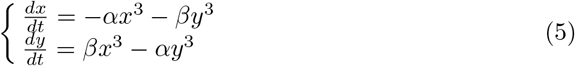

In the partial reconstruction scenario, we assume the nonlinear coupling terms between the variables to be unknown, and we replace them with an NN, yielding:

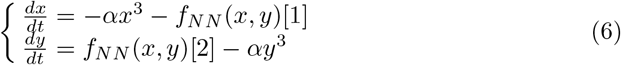

where, *f*_*NN*_ : ℝ^2^ → ℝ^2^ takes (*x, y*) as input and approximates the unknown terms.

The third system is the Lorenz system [46], originally introduced to model atmospheric convection and widely studied as a canonical example of chaotic dynamics [52]. It is defined by the three-dimensional ODE system:

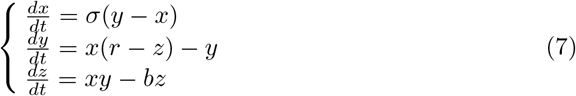

In the partial reconstruction scenario, we assume that the equation governing the dynamics of the third variable is entirely unknown and replace it with a neural network *f*_NN_ : ℝ^3^ → ℝ, taking (*x, y, z*) as input.

As a biological case study, we consider the cell apoptosis model introduced in [49], which describes the dynamics of eight interacting molecular species. The model is defined by the following system of ODEs:

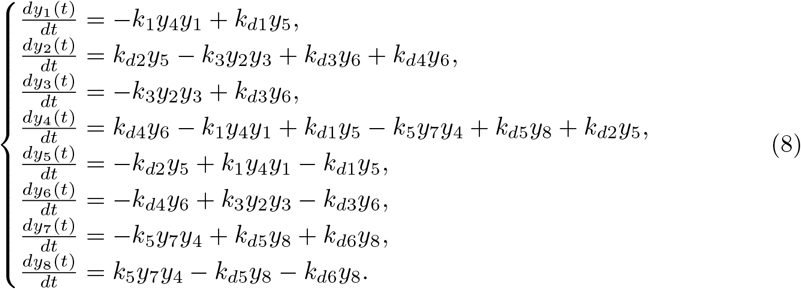

In the partial reconstruction scenario, we assume that the equation governing the dynamics of the fourth variable is entirely unknown and replace it with a neural network *f*_NN_ : ℝ^6^ → ℝ, taking (*y*_1_, *y*_4_, *y*_5_, *y*_6_, *y*_7_, *y*_8_) as input.

### Prediction intervals on vector field

In this and the following sections, we will extend the notation used in Section Proposed method. We denote by

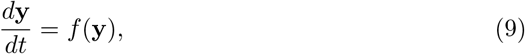

with **y** ∈ ℝ^*n*^, the ground-truth model to be reconstructed, and by **y**(*t*; *t*_0_, **y**_0_) the state of the ground-truth system at time *t*, obtained from the initial condition (*t*_0_, **y**_0_), with *t > t*_0_, given by

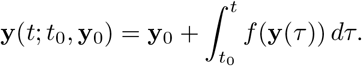

We denote by *y*^*j*^(*t*; *t*_0_, **y**_0_) the *j*-th component of **y**(*t*; *t*_0_, **y**_0_), i.e., the state of the *j*-th variable of the system at time *t*. We consider an ensemble of *p* UDE models trained to approximate the dynamics of the ground-truth system, that is, *p* parameterizations ***θ***_1_, …, ***θ***_*p*_ ∈ ℝ^*k*+*l*^, each trained to reproduce the training data describing the time evolution of the dynamical system in Eq. 9. For a fixed parameterization 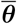, we denote by

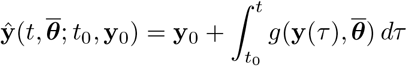

the prediction of the UDE model at time *t*, obtained by integrating the system from the initial condition (*t*_0_, **y**_0_), with *t > t*_0_. We further indicate with 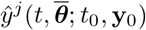 the *j*-th component of the predicted state.

Given a point in the state space **z** ∈ ℝ^*n*^, the prediction intervals for the value of the vector field in **z**, *f* (**z**), are computed by extending to the multivariate setting the procedure proposed for the univariate case in NN ensembles [29]. Fixed a level *α* ∈ (0, 1), the *α*-prediction interval on *f* (**z**) is computed, following [53], as the ellipsoidal region:

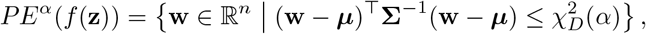

where ***µ*** and **Σ** are the sample mean and covariance matrix of the ensemble predictions *g*(**z, *θ***_1_), …, *g*(**z, *θ***_*p*_), respectively. The term 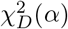 denotes the *α*-quantile of the chi-squared distribution with *D* degrees of freedom, corresponding to the number of vector field components being regressed. In our work, we use *α* = 0.95 and set *D* = *n* for every test case, except for the Lorenz system in the UDE with fixed physical parameters. In this case, we use *D* = 1 and *n* = 3, given that the estimation process involves parameters from only a single equation in the model.

### Prediction intervals on trajectories

The prediction intervals on the ground truth trajectories of the dynamical system in Eq. (9) are computed individually for each variable of the dynamical system. For a fixed *α* ∈ (0, 1), the *α*-prediction interval for the *j*-th variable at time *t* is computed as

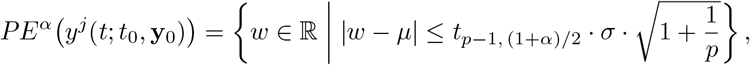

where *µ* and *σ* denote the sample mean and standard deviation of the ensemble predictions *ŷ*^*j*^(*t*, ***θ***_1_; *t*_0_, **y**_0_), …, *ŷ*^*j*^(*t*, ***θ***_*p*_; *t*_0_, **y**_0_), and *t*_*p*−1, (1+*α*)*/*2_ is the (1 + *α*)*/*2 quantile of the Student distribution with *p* − 1 degrees of freedom [54]. In the presence of noise in the training dataset, aleatoric uncertainty was incorporated as suggested in [29].

### OOD input sets

In this section, we describe how the OOD input sets–used to evaluate the coverage proportion of prediction intervals–were constructed. This is valid only for the numerical test cases, as in the biological test case, we evaluate the coverage on a single biologically meaningful prediction of the model unseen during the training process.

**OOD vector field predictions**: in each numerical test case, the OOD region of the state space where the 0.95-prediction intervals are evaluated is constructed by symmetrically expanding the bounding box of the training trajectories in the state space by 20% along each coordinate axis. The set of OOD inputs is derived by sampling 50000 points uniformly from this region.

**OOD trajectory predictions**: in each numerical test case, we generated 100 test trajectories by randomly selecting one of the training trajectories and perturbing its initial condition. The perturbation was sampled from a normal distribution with mean 0 and coefficient of variation 0.5.

### Standard ensemble training

In this section, we report the training of the standard ensembles. Each model was trained following the same process, outlined in the following, with the only variation being the random initialization of the model parameters. Given a time series of observed system states 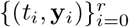, with **y**_*i*_ ∈ ℝ^*n*^, we randomly split the indices {1, …, r} into disjoint training and validation sets

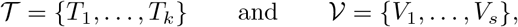

respectively, according to an 80:20 ratio. Let 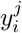 denote the *j*-th component of the observed state **y**_*i*_. To account for differences in variable magnitudes, we define Γ^*j*^ as the maximum variation of the *j*-th component across the trajectory:

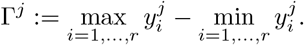

The normalized mean-squared error loss functions for training and validation used are:

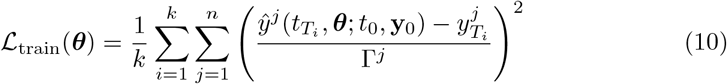

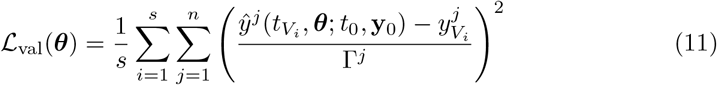

Since minimizing *L*_train_ directly is often suboptimal due to the potential risks of the training getting stuck in local minima [55], we articulate the training process in two different steps. In the first step, we use the multiple shooting technique [56] to initially optimize the fit of the model to the training set. The result from this initial step is refined in the second step, where the training loss function is optimized using the L-BFGS algorithm [57]. Since training NODEs and UDEs can encounter convergence issues and numerical instabilities, this may result in parameterizations with suboptimal fit to the training data. The optimization results are accepted, meaning the parameters are selected to be part of an ensemble, if

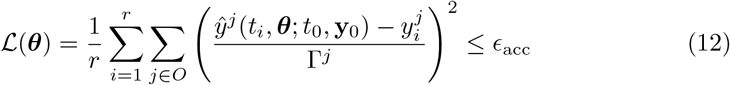

where *ϵ*_acc_ is a tolerance threshold and *O* denotes the set of observable variables of the system. In the numerical test cases presented in Section Numerical test cases: benchmark dynamical systems and evaluation metrics, we set *ϵ*_acc_ = 10^−3^. In the cell-apoptosis test case, due to the presence of noise in the training dataset, we instead set *ϵ*_acc_ = 5 *·* 10^−3^.

### Disagreement measure of reconstructed vector fields

Given a point in the state space 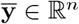, we measure the disagreement of an ensemble of models defined by the parameterizations **b**_**1**_, …, **b**_**p**_ as follows:

- If *p* ≥ *n*, we consider the generalized variance [58] of the vectors 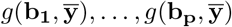, i.e., the scalar quantity

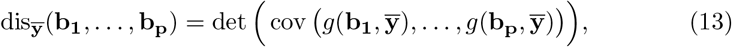

that, intuitively, measures the overall variability of the vectors 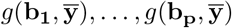 [58].
- If 1 *< p < n*, since the determinant in Eq. (13) is zero, we consider the sum of the variances along each axis of the space, that is:

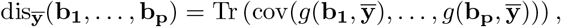

where Tr(·) denotes the trace operator, i.e., the sum of the diagonal entries of the covariance matrix.

We can summarize our disagreement measure as:

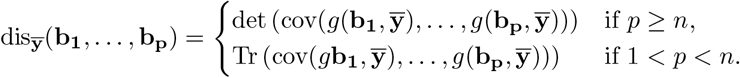

For a finite set of points 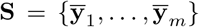, the disagreement of the models on **S**, indicated with dis_**S**_(**b**_**1**_, …, **b**_**p**_), is computed as the average disagreement across points in **S**.

### MOD ensemble training: constraint optimization step

Our approach is inspired by constrained optimization on Riemannian manifolds [36]. Since we do not know whether **B** is itself a Riemannian manifold (although mode connectivity results for standard NNs suggest that, in several cases, the set of minimum-loss solutions forms a manifold [59]), we motivate our method intuitively as follows. Suppose we have an ensemble of *p* parameterizations **b**_**1**_, …, **b**_**p**_ ∈ ℝ^*k*+*w*^ minimizing the loss function, i.e. **b**_**o**_ ∈ **B** for each *o* = 1, …, *p*. Let **S** ⊂ ℝ^*n*^ be a finite set of points in the state space of the model. Now assume that **b**_**1**_, …, **b**_**p**−**1**_ are fixed, and we seek to locally move **b**_**p**_ in order to increase the disagreement among the parameterizations **b**_**1**_, …, **b**_**p**_ on the set **S**, while maintaining **b**_**p**_ ∈ *B*. If our goal was solely to locally increase the disagreement among the parameterizations in **S**, the natural update direction for **b**_**p**_ would be given by the gradient:

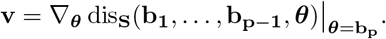

However, an update of **b**_**p**_ along the direction **v** does not necessarily result in a point that remains within **B**. On the other hand, denoting with *H* the Hessian matrix of ℒ evaluated at **b**_**p**_, the null space *H*^0^ of *H*—i.e., the eigenspace corresponding to zero eigenvalues, defines the subspace of directions along which the loss does not increase [43]. Therefore, the projection of **v** onto *H*^0^ locally identifies the direction along which disagreement can be maximally increased without leaving the minimum-loss set **B**. Our approach proceeds by locally updating **b**_**p**_ along this projected direction at each step. To compute the Hessian matrix, the loss function ℒ, computed over the entire training trajectories, is the same as the one used in the acceptance criterion for standard ensemble training, as defined in Eq. (12). We approximate the Hessian matrix *H* of ℒ at a point 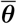 using the Gauss-Newton method. When computed numerically (under finite-precision arithmetic), the eigenvalues of *H*_ℒ_ (***θ***) are always strictly positive. To identify directions in the null space 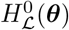, we approximate zero eigenvalues by applying a threshold *ϵ*_*eig*_, such that eigenvalues smaller than *ϵ* are considered null.

### MOD ensemble training: how to route loss contours

Given *p* − 1 models in an ensemble (with *p >* 1), i.e., *p* − 1 parameterizations **b**_1_, …, **b**_*p*−1_, the *p*-th model parameterization **b**_*p*_ is initialized at **b**_1_. We first perturb it randomly along the null space of the cost function 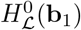, and then perform a fixed number of iterations *n*_iter_. In each iteration, as described in Section Proposed method, we update the parameters **b**_*p*_ according to the direction **w**, defined by the projection of the gradient of the disagreement

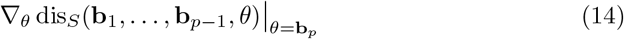

onto 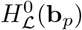.

Since the direction **w** guarantees no increase in the loss function ℒ only locally, we introduce a threshold *ϵ*_ℒ_ to define the maximum allowable value of ℒ for ensemble members. To ensure that optimization remains within this minimum-loss region, we adopt two complementary strategies. First, at each iteration of the MOD training algorithm, the parameters are updated along the direction **w** using an adaptive learning rate. Specifically, the learning rate for updating **b**_*p*_ along **w** is dynamically chosen so that it is as large as possible without causing the loss to increase by more than a prescribed threshold *ϵ*_step_. This adaptive rule allows us to control the learning rate at each iteration to remain on the level sets of ℒ. Second, if after an update, even with a minimum learning rate, the value ℒ (**b**_*p*_) exceeds the threshold *ϵ*_acc_, we perform a short optimization phase to explicitly minimize ℒ, effectively re-projecting **b**_*p*_ back onto the minimum-loss region. These two mechanisms work in tandem to guide the optimization along the level curves of the loss while maintaining the desired loss constraint. We adopt a stochastic approach, and at each optimization step, we randomly sample a subset of *S*^*′*^ ⊂ *S* and approximate the gradient of Eq. (14) with

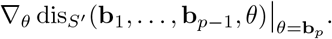

### MOD ensemble training: overall algorithm description

The overall algorithm is summarized in Algorithm 1. For each model in the ensemble, we optimize only a subset of the NN parameters in ***θ***, in order to further encourage diversity among ensemble members.

In the case of hybrid models where ***θ*** includes both mechanistic and NN parameters (as in the third scenario in Section Numerical test cases: benchmark dynamical systems and evaluation metrics), the mechanistic parameters are normalized with respect to their initial values—i.e., the values optimized in **b**_1_.

### Comparison between MOD and standard ensembles

In each scenario presented, we train 10 standard ensembles of five models using a fixed architecture, differing only in the random initialization of the NNs and mechanistic parameters when applicable (for a discussion about what the size of the ensemble implies in terms of uncertainty quantification, we refer to Supplementary Section S1). From each standard ensemble, one model is randomly selected as the initialization point **b**_1_ for constructing a corresponding five-member MOD ensemble. This design enables pairwise comparison between standard and MOD ensembles. Since NODE/UDE training may converge to local minima, an acceptance threshold *ϵ*_acc_ on the loss function is introduced. Models are included in standard ensembles only if their loss is below *ϵ*_acc_, and the same tolerance is used in the MOD algorithm to control how to route the contour curves of the loss function (see Section MOD ensemble training: overall algorithm description for details about the algorithm). Standard ensembles are trained until five models satisfy this criterion, ensuring that both standard and MOD ensembles contain exactly five members.

#### Algorithm 1

MOD Ensemble Training Procedure

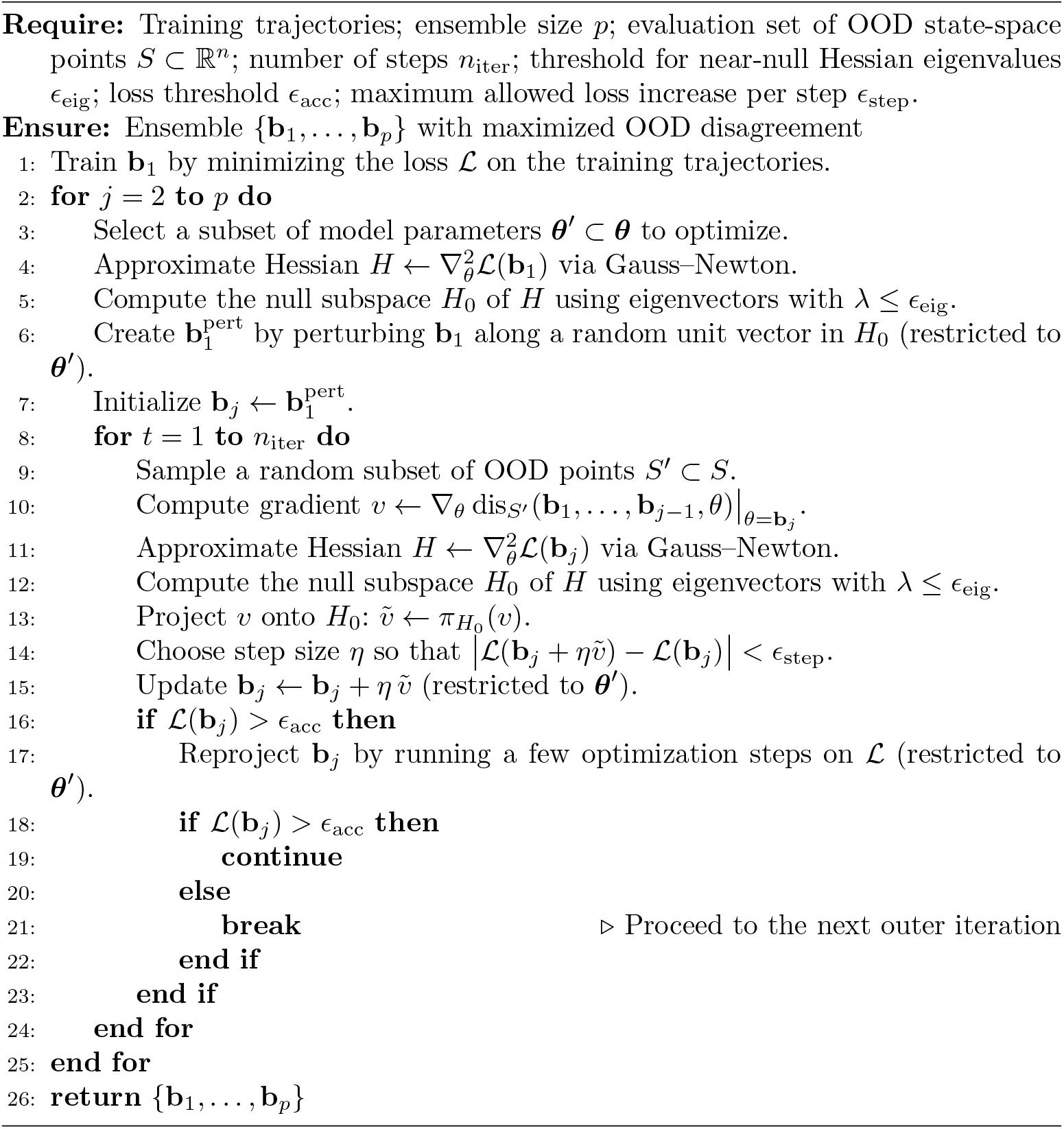

### Algorithmic setup

The results on the numerical test cases reported in the main text of the paper were obtained by running the MOD ensemble training algorithm with *ϵ*_acc_ and *n*_iter_ = 800. The step-size threshold *ϵ*_step_ was set as *ϵ*_acc_ · 10^−1^ = 10^−4^. At each iteration, the Hessian of the cost function was computed on a randomly sampled subset of 20 time points, selected from the total 100 time points available for each trajectory. Each ensemble member was trained by optimizing 200 NN parameters, plus mechanistic parameters if applicable. The finite set *S*, used to maximize model disagreement, was constructed as a regular grid over the OOD region of the state space. Specifically, we used a 10 *×* 10 grid for the Lotka–Volterra and the Damped Oscillator test cases, and a 5 × 5 × 5 grid for the Lorenz system. At each iteration, a random subset comprising 25% of the grid points was used to compute the gradient of the disagreement. The parameter projection, applied when the loss on the training trajectory exceeded *ϵ*_*L*_, was carried out by optimizing the loss function using 20 iterations of gradient descent, followed by 20 iterations of the L-BFGS algorithm. The threshold to identify near-null eigenvectors was fixed at *ϵ*_*eig*_ = 10^−1^.

The results on the numerical test cases presented in the Supplementary Material, which were obtained with a lower loss threshold, were generated using *ϵ*_acc_ = 10^−4^ and a correspondingly updated step-size threshold *ϵ*_step_ = 10^−5^. In this case, due to the smaller step sizes, the number of training steps was increased to *n*_iter_ = 1200 for each ensemble member. All the other settings are the same as in the ones reported in the main text.

The results for the cell apoptosis model reported in the main text, where noise is added to the training data, were generated using *ϵ*_acc_ = 5 × 10^−3^ and *n*_iter_ = 300, with a correspondingly adjusted step-size threshold *ϵ*_step_ = 5 × 10^−4^. To identify the region of the state space in which the disagreement among models should be maximized, we first simulated the cell-death scenario using the first model of the MOD ensemble. We then expanded the hypercube defined by the minimum and maximum values of the system variables by 50% along each axis. Given the higher dimensionality of the state space (comprising eight variables), we generated 1000 random points within this region and, at each iteration, selected 100 random points to compute the gradient of the disagreement. All other settings are identical to those used in the numerical test cases.

### Implementation

All the computations have been run on a Debian-based Linux cluster node, with two 24-core Xeon(R) CPU X5650 CPUs and 250 GB of RAM. The code is implemented in Julia v1.10.5 [60], using the SCiML environment [13], a software suite for modeling and simulation that also incorporates ML algorithms.

## Data availability

No experimental data were used as part of this study. The *in silico* generated datasets are available at https://github.com/cosbi-research/UQNU.

## Code availability

The scripts to reproduce the results are available at https://github.com/cosbi-research/UQNU.

## Acknowledgements

No funding was received for this research.

## Author contributions statement

S.G. ideated the workflow and selected the test cases under the supervision of L.M. and G.I.. S.G. developed the computational workflow. L.M. and G.I. provided overall guidance for the project. All authors contributed to manuscript preparation and approved the final version.

## Competing Interests

The authors declare no competing financial or non-financial interests.

## Supplementary information

The supplementary material is available in the file

***S1 File***

**Supplementary notes, tables and figures**

## Notes

### Competing Interest Statement

The authors have declared no competing interest.

### Summary of Updates

Updated the manuscript with a computational biology test case and improved analysis of numerical test cases.

