## Supplementary material for "A novel approach to quantify out-of-distribution uncertainty in Neural and Universal Differential Equations": S1 File

#### S1 File: Supplementary Sections, Tables and Figures

#### S1 Analysis of standard ensembles

In this section, we show that in the context of dynamical system reconstruction, standard ensembles risk being overconfident when quantifying uncertainty under Out Of Distribution (OOD) conditions. We consider the task of modeling, in a purely data-driven way, the three numerical test case dynamical systems described in Section **Methods: Test cases** of the main text. In this setting, we employ pure Neural Ordinary Differential Equations (NODEs)

$$\frac{d\mathbf{y}}{dt} = f_{NN}(\mathbf{y}, \boldsymbol{\theta}_{NN}), \tag{1}$$

with $\mathbf{y} \in \mathbb{R}^n$, with $n = 2$ for the Lotka–Volterra and Damped Oscillator systems, and $n = 3$ for the Lorenz system.

Our goal is to evaluate the coverage properties (see Section **Results: Numerical test cases: benchmark dynamical systems and evaluation metrics** of the main text) of the prediction intervals obtained with standard ensembles both on the vector fields and on the system trajectories. To this end, we train NODE ensembles on the three training trajectories detailed in the Section **Methods: Test cases** of the main text, by only varying the random initialization of $\boldsymbol{\theta}_{NN}$, and evaluate the coverage performances using the metrics introduced in the previous section. The training procedure of the NODEs is detailed in Section **Methods: Standard ensemble training** of the main text. The NN $f_{NN}$ used for all the test cases consists of an input layer, two hidden layers with 32 nodes each employing the *gelu* activation function, and an output layer, where the dimensionality of the respective dynamical system determines input and output dimensions. For each test case, to account for the variability arising from the stochasticity of NN training, we train 10 ensembles on the training set, with each ensemble comprising 5 NNs.

The quality of the fit on the training trajectories for the different test cases is illustrated in Figs. S1, S2, and S3. For each test case, a total of 50 models were trained (5 models per ensemble across 10 ensembles).

The influence of considering different training datasets and of the ensemble size—an inherently arbitrary modeling choice— will be discussed at the end of this section. The coverage properties of the resulting 0.95-prediction intervals are reported in Fig. S4.

We begin by analyzing the coverage of prediction intervals on the vector field. In the Damped Oscillator and Lorenz test cases, the mean CP indicates that the prediction intervals are overconfident in the region considered: the mean coverage proportion (CP) values are substantially below the nominal value of 0.95, and the CP distributions are significantly lower than 0.95 (Wilcoxon signed-rank test, $p < 0.001$ for both cases). In both systems, the mean coverage heatmaps reveal extensive regions in the state space where all trained ensembles produce prediction intervals that consistently fail to cover the ground truth. The situation differs for the Lotka–Volterra case, where the mean CP is closer to the nominal value of 0.95, although the CP distribution remains significantly lower than this value (Wilcoxon signed-rank test, $p = 0.004$). This result depends on the region of the state space analyzed: when the region is extended, the overconfidence of the prediction intervals becomes more evident here as well, consistent with the other test cases. To prove it, we report the analysis is performed over an extended region of the state space in the Lotka–Volterra test case, as shown in Fig. S5. The extended region is constructed by expanding each side of the original state space region used in the previous analysis by 100%. The mean CP and overall coverage are computed across 10 independently trained ensembles, each consisting of 5 models.

The coverage of the prediction intervals on the test trajectories is consistent with the results observed on the vector field. Specifically, it confirms the overconfidence of the prediction intervals for the Damped Oscillator and Lorenz test cases, with mean CP values of 0.63 and 0.59, respectively—well below the nominal

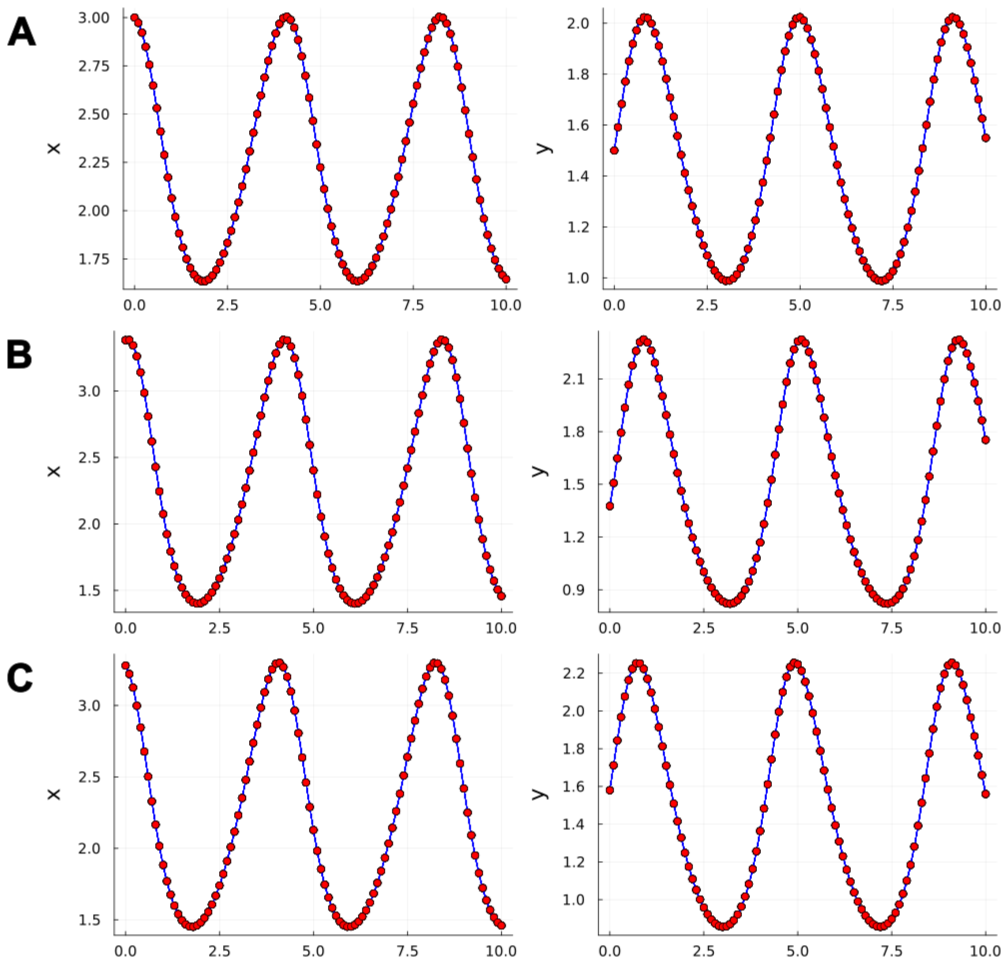


Figure S1: **Fit of trained models on training trajectories for the Lotka–Volterra test case.** Panels A, B, and C show the fits of the trained models (blue lines) to the data points of the first, second, and third training trajectories (red points), respectively.

value of 0.95—and CP distributions significantly lower than 0.95 (Wilcoxon signed-rank test, $p < 0.001$ for both cases). Fig. S6 shows two example test trajectories for which the coverage of the prediction intervals is low. In the Lotka–Volterra test case, the mean CP is 0.80, closer to the nominal value with respect to the other test-cases, with a CP distribution significantly lower than this value (Wilcoxon signed-rank test, $p < 0.001$).

We conclude the analysis by investigating the effect of varying ensemble sizes on the properties of the resulting prediction intervals in this scenario. One might expect that increasing the ensemble size introduces greater variability in the outputs, potentially leading to wider and more conservative prediction intervals. The impact of ensemble size on uncertainty quantification under OOD conditions has previously been explored by Ovadia et al.[GIP$^+$18], who, in standard ML settings (regression and classification), found that ensembles of 5 models often provide a sufficient trade-off between performance and computational cost, with diminishing returns beyond this point. Here, we test whether a similar hypothesis holds in the context of dynamical system reconstruction. To this end, we trained ensembles of sizes 5, 10, 20, and 30, with 5 ensembles for each size, and compared the resulting CP distributions for both vector fields and trajectories. The results, presented in Fig. S7, show that increasing the ensemble size does not produce a clear upward trend in CP for either trajectories or vector fields in any of the test cases. Figs. S8, S9, and S10 show how the mean coverage across the evaluated region of the state space varies with ensemble size for the Lotka–Volterra, Damped Oscillator, and Lorenz cases, respectively. These results suggests that enlarging the ensemble does

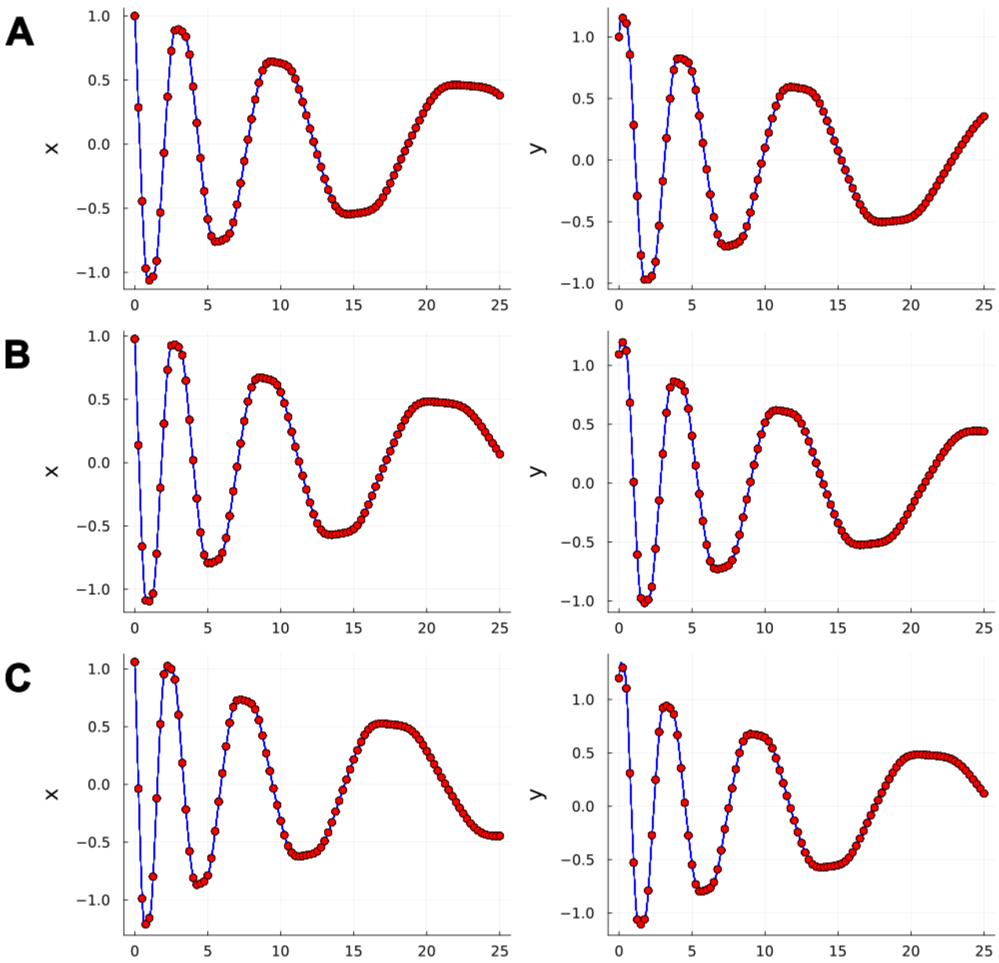


Figure S2: **Fit of trained models on training trajectories for the Damped Oscillator test case.** Panels A, B, and C show the fits of the trained models (blue lines) to the data points of the first, second, and third training trajectories (red points), respectively.

not substantially increase variability in OOD predictions.

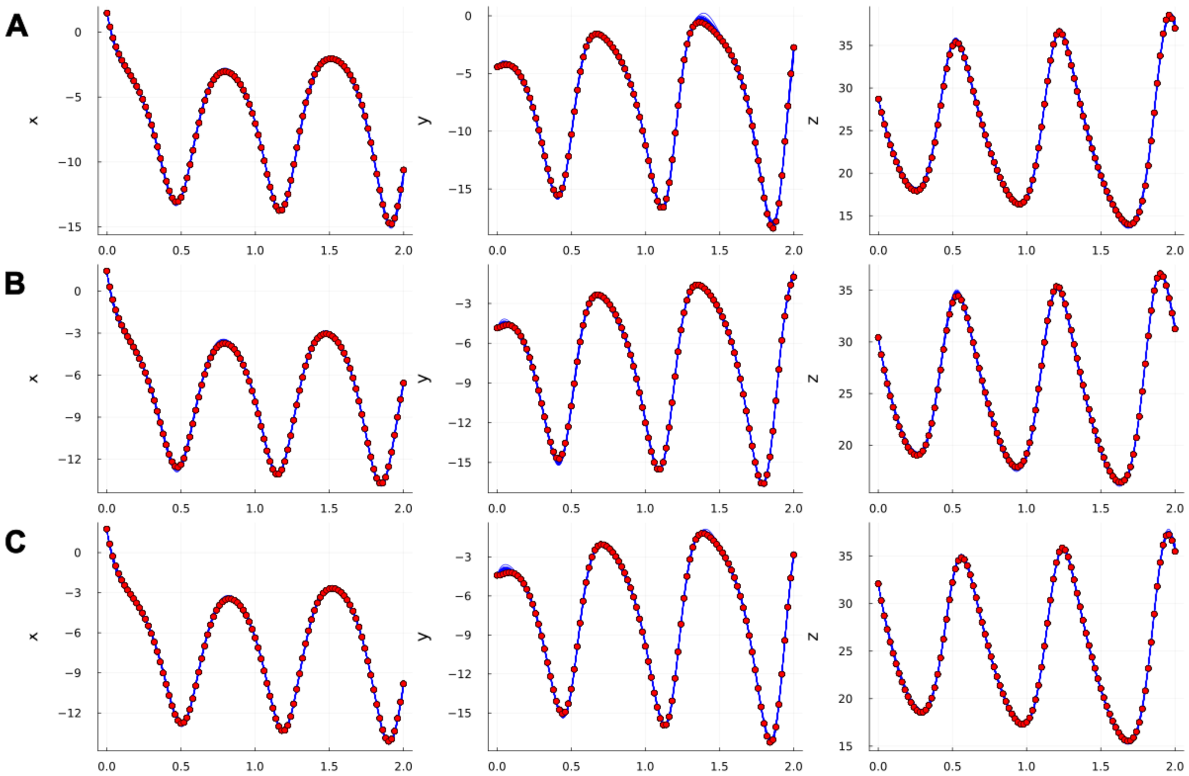


Figure S3: **Fit of trained models on training trajectories for the Lorenz test case.** Panels A, B, and C show the fits of the trained models (blue lines) to the data points of the first, second, and third training trajectories (red points), respectively.

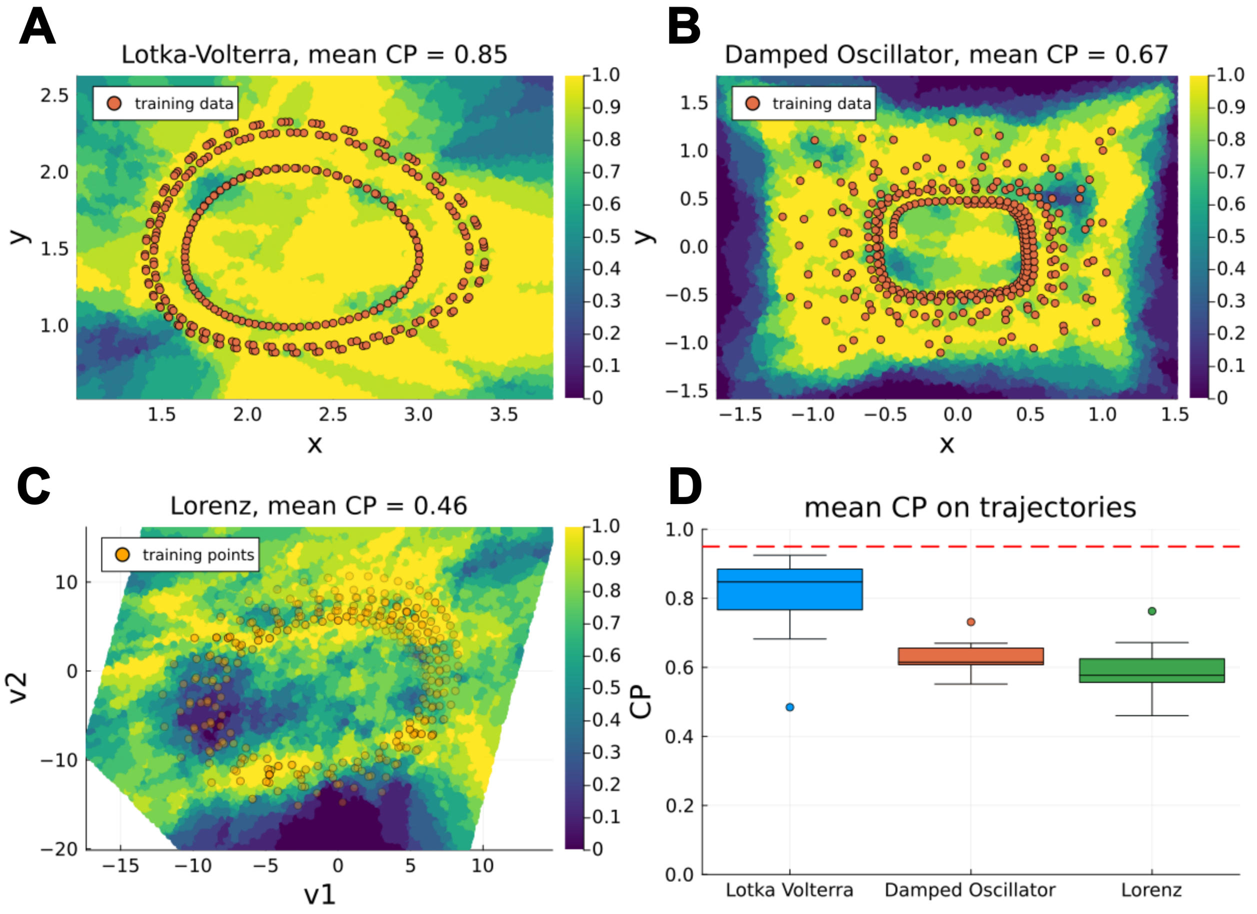


Figure S4: **Coverage of 0.95-prediction intervals with standard ensembles for data-driven reconstruction of dynamical systems.** Panels A, B, and C display heatmaps of the mean coverage on the vector field, computed from 10 independently trained ensembles, for the Lotka–Volterra, Damped Oscillator, and Lorenz systems, respectively. Each panel title indicates the resulting mean CP (theoretical nominal coverage level of 0.95). Orange overlays represent the points of the training trajectories, providing spatial context. For the Lorenz system, the mean coverage is visualized on a two-dimensional plane spanned by the coordinates $(v_1, v_2)$, obtained via linear regression on the Lorenz attractor (see Supplementary Section S8 for details). These coordinates provide a low-dimensional representation of the original three-dimensional state space $(x, y, z)$ ; here, training trajectory points are color-scaled based on their orthogonal distance to the plane. Panel D presents the distributions of CPs over reconstructed trajectories across the 10 ensembles trained for the different test cases. The dashed red line represents the theoretical nominal coverage level of 0.95, while the points outside the box plots represent outliers of the CP distributions.

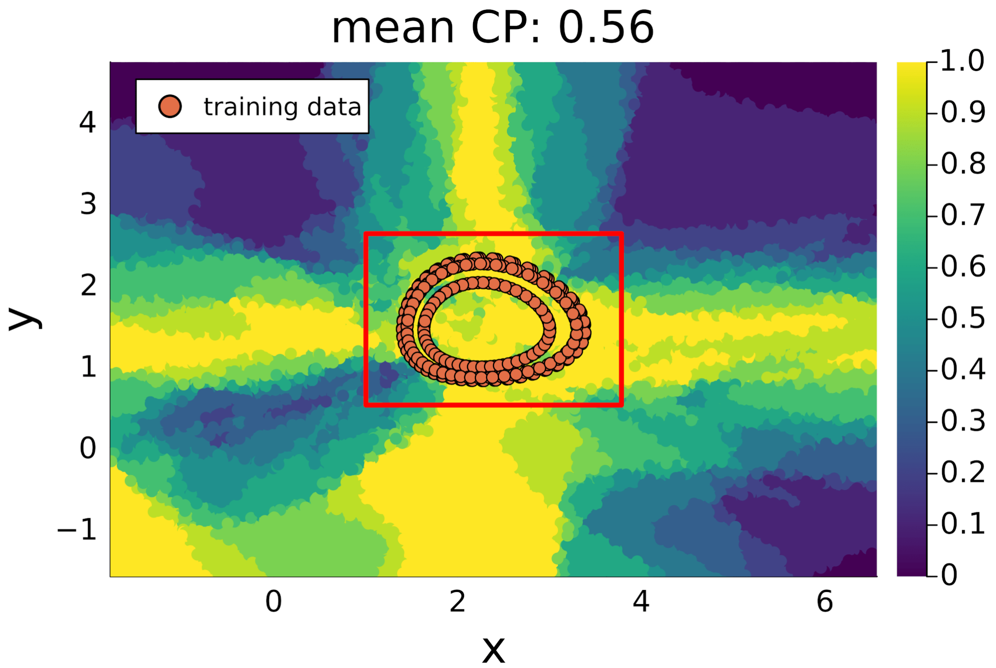


Figure S5: **Coverage of 0.95-prediction intervals using standard ensembles on an extended region of the state space for the Lotka–Volterra test case.** The heatmap shows the mean coverage of the vector field across an extended region of the state space, computed from 10 independently trained ensembles. Each panel corresponds to the Lotka–Volterra test case, with the title indicating the mean coverage proportion (CP) over the evaluated region. Orange overlays mark the locations of the training trajectories, providing spatial context. The red rectangle highlights the region of the state space considered in the main text.

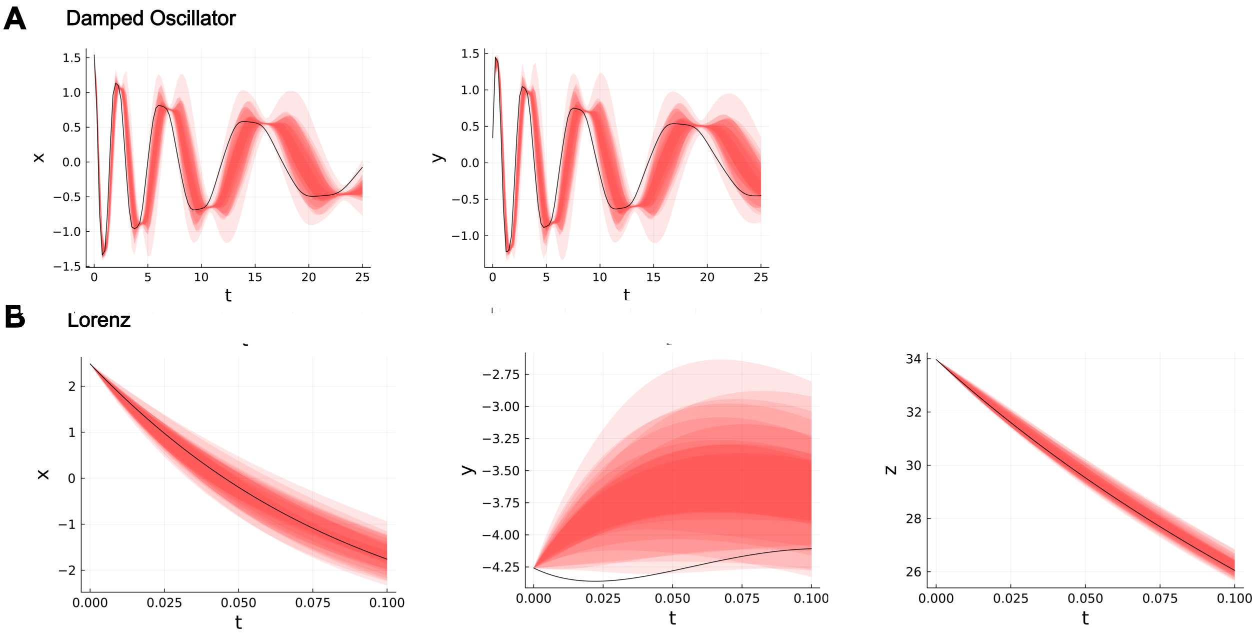


Figure S6: **Examples of prediction intervals failing to cover the ground-truth trajectories with standard ensembles.** Panel A shows a test trajectory (solid black line) of the Damped Oscillator system, where none of the variables is fully covered by the 0.95 prediction intervals (shaded red regions) produced by the trained standard ensembles. Panel B shows a test trajectory (solid black line) of the Lorenz system, where the values of the $y$ variable are covered by only one of the thirty 0.95 prediction intervals produced by the trained standard ensembles (shaded red regions). The different red shades result from the overlap of prediction intervals computed from the ten different ensembles.

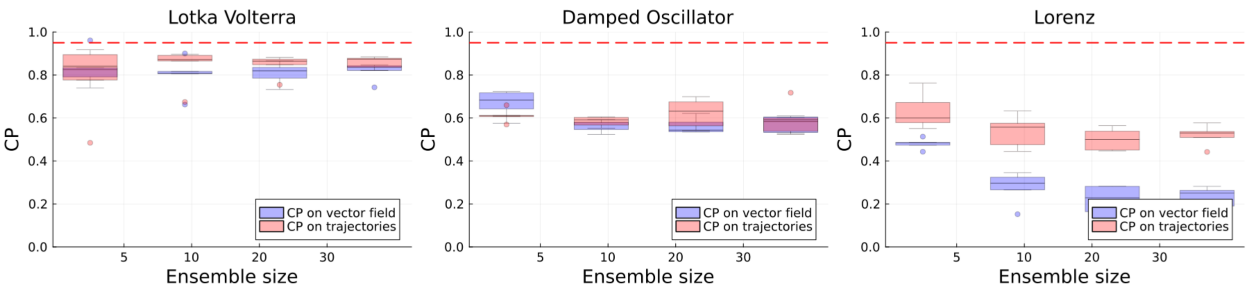


Figure S7: **CP on vector fields and trajectories with varying standard ensemble sizes for a purely data-driven reconstruction of dynamical systems.** CP distributions—shown in red for vector fields and in blue for trajectories—under OOD scenarios for the Lotka–Volterra, Damped Oscillator, and Lorenz test case considering different sizes of ensembles. The dashed red line represents the theoretical nominal coverage level of 0.95, while the points outside the box plots represent outliers of the CP distributions.

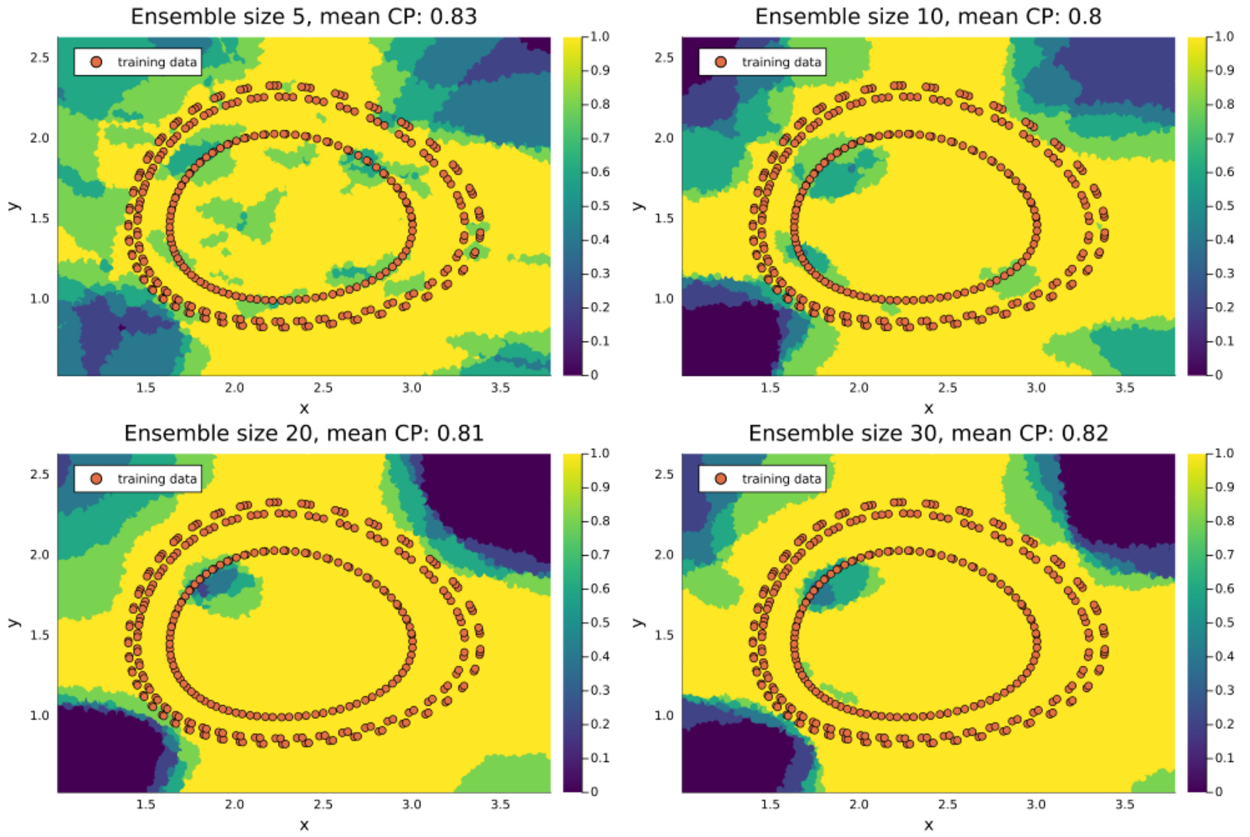


Figure S8: **Coverage of 0.95 prediction intervals on the vector field for standard ensembles of varying size (Lotka–Volterra test case).** The heatmaps display the mean coverage of the vector field over the considered region of the state space, computed from five independently trained ensembles of varying sizes. Each plot title indicates the ensemble size and the corresponding mean CP across the evaluated region. Orange overlays highlight the locations of the training trajectories, providing spatial context.

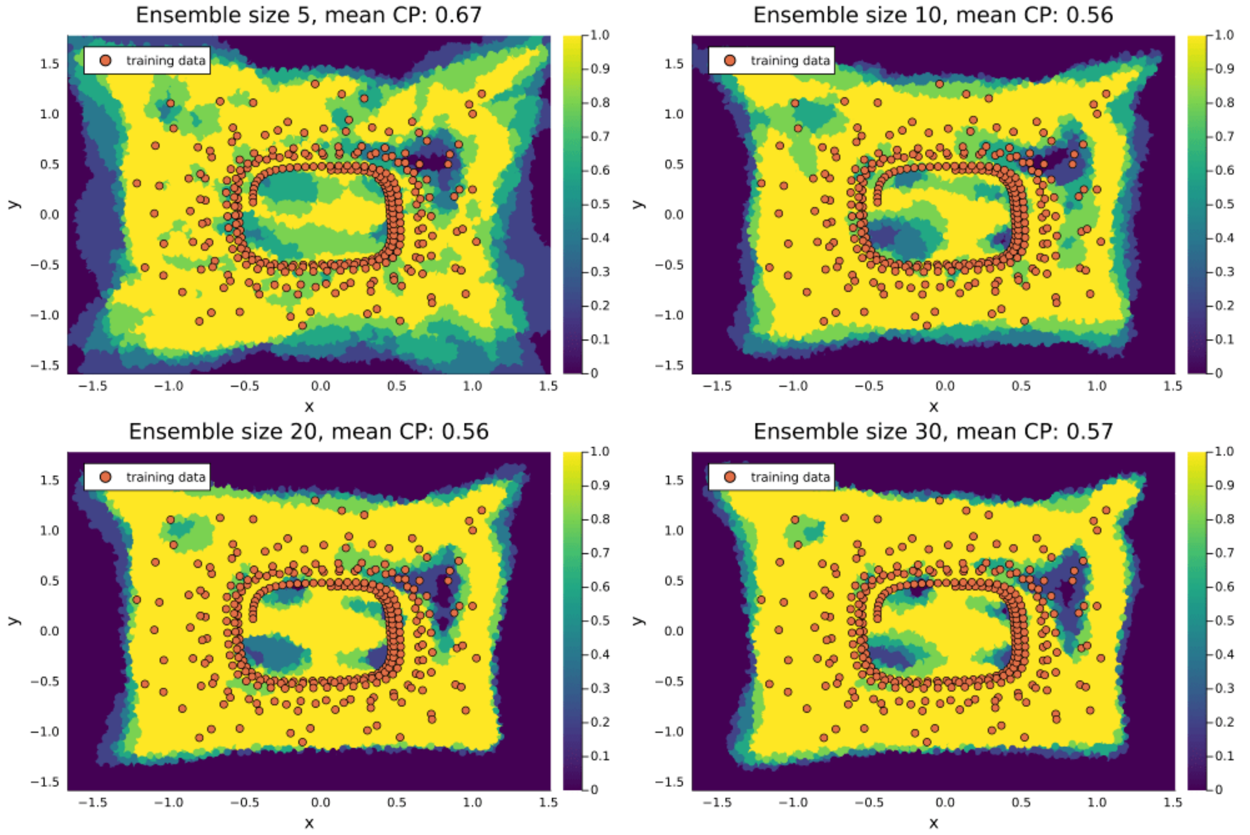


Figure S9: **Coverage of 0.95 prediction intervals on the vector field for standard ensembles of varying size (Damped Oscillator test case).** The heatmaps display the mean coverage of the vector field over the considered region of the state space, computed from five independently trained ensembles of varying sizes. Each plot title indicates the ensemble size and the corresponding mean CP across the evaluated region. Orange overlays highlight the locations of the training trajectories, providing spatial context.

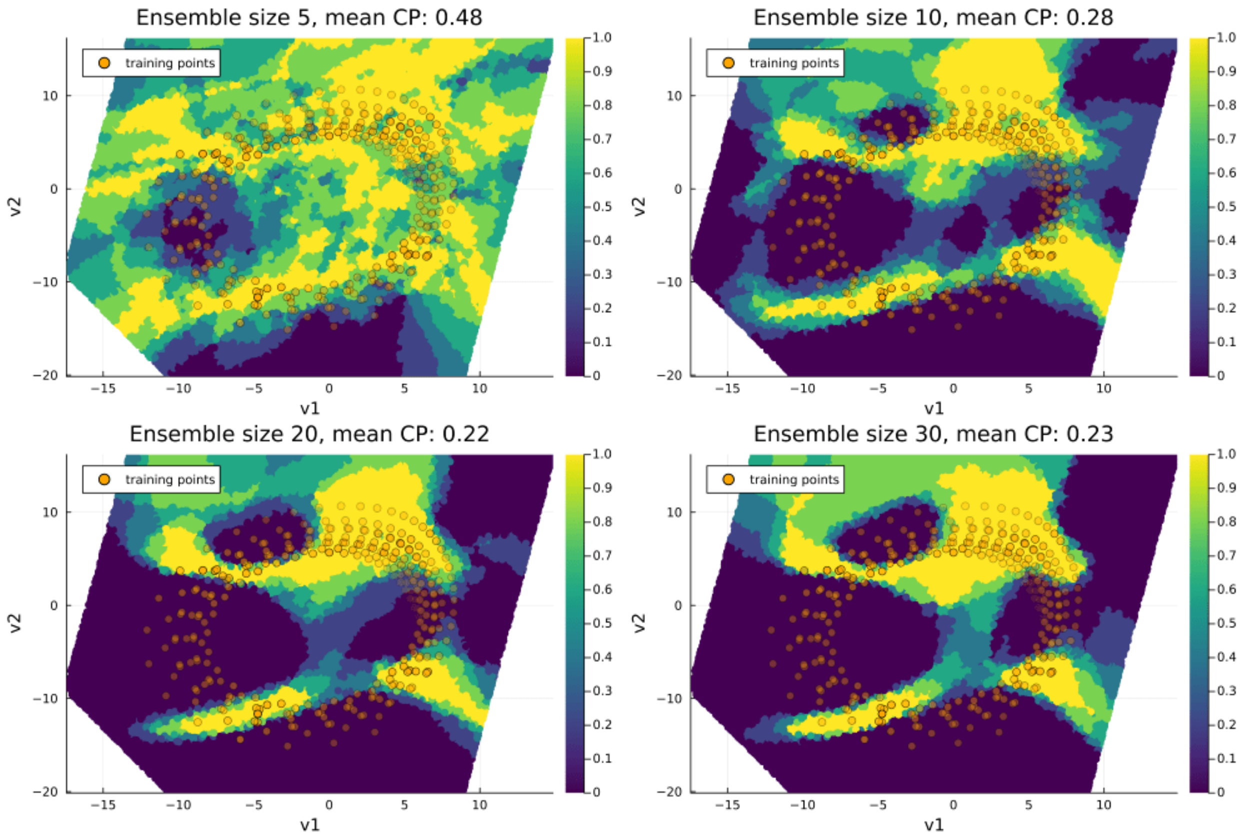


Figure S10: **Coverage of 0.95 prediction intervals on the vector field for standard ensembles of varying size (Lorenz test case).** The heatmaps display the mean coverage of the vector field, visualized on a two-dimensional plane spanned by the coordinates $(v_1, v_2)$, obtained via linear regression on the Lorenz attractor (see Supplementary Section S8 for details). These coordinates provide a low-dimensional representation of the original three-dimensional state space $(x, y, z)$. The mean coverage is computed from five independently trained ensembles of varying sizes. Orange overlays highlight the locations of the training trajectories, providing spatial context. The color is scaled based on the orthogonal distance between the training points and the visualization plane.

#### S2 Mode connectivity in NODEs and UDEs

In this section, we extend the proof of mode connectivity —previously established for standard NNs- to NODEs (a), and UDEs with known mechanistic parameters (b), under specific hypotheses on the training set; and discuss the validity of the mode connectivity assumption for UDEs with unknown mechanistic parameters (c). We conclude by analyzing the implications of relaxing or violating the key assumptions underlying the proof of mode connectivity.

**a. Extension of the proof of mode connectivity to NODEs.** Here, we build upon the proof provided for neural networks by Şimşek *et al.* [SGJ+21], who demonstrated that the zero-loss subspace of the parameter space of an over-parameterized $L$-layer neural network forms a connected manifold. Moreover, in the case $L = 2$, this connected manifold is characterized as a union of affine subspaces under the assumptions of a smooth activation function and infinitely many training data points with full support over the input space [SGJ+21]. Under similar conditions, we extend these results to over-parameterized NODEs.

Let us consider a training dataset describing the evolution of a dynamical system over a time interval $[0, T]$, with $T \in \mathbb{R}^+$:

$$\{(t_i, \mathbf{y}_i)\}_{i \in \mathrm{Tr}},$$

where $t_i \in [0, T]$ denotes time and $\mathbf{y}_i \in \mathbb{R}^n$ represents the state of the system at time $t_i$. Following the setup of Şimşek *et al.*, we assume that the set of time points $\{t_i\}_{i\in\mathrm{Tr}}$ is dense in the interval $[0, T]$, i.e.,

$$\overline{\{t_i\}_{i\in\mathrm{Tr}}} = [0, T],$$

where the overline denotes the closure of the set. We now consider an $L$-layer NODE defined by

$$\frac{d\mathbf{y}}{dt} = f_{NN}(\mathbf{y}, \boldsymbol{\theta}_{NN}),$$

where $f_{NN}$ is an $L$-layer neural network and $\boldsymbol{\theta}_{NN} \in \mathbb{R}^k$ are its parameters. Suppose two parameterizations, $\boldsymbol{\theta}^1_{NN}$ and $\boldsymbol{\theta}^2_{NN}$, achieve zero training loss, meaning that

$$\int_0^{t_i} f_{NN}(\mathbf{y}, \boldsymbol{\theta}^1_{NN})\, dt = \mathbf{y}_i \quad \text{and} \quad \int_0^{t_i} f_{NN}(\mathbf{y}, \boldsymbol{\theta}^2_{NN})\, dt = \mathbf{y}_i \tag{2}$$

for all $i \in \mathrm{Tr}$. Under the assumption that $\{t_i\}_{i\in\mathrm{Tr}}$ is dense in $[0, T]$ and using the uniqueness of the integral limit, this implies that

$$f_{NN}(\mathbf{y}, \boldsymbol{\theta}^1_{NN}) = f_{NN}(\mathbf{y}, \boldsymbol{\theta}^2_{NN}) \tag{3}$$

for all $\mathbf{y}_i$ in the training set. Consequently, the zero-loss set of NODE parameters corresponds to the zero-loss set of a standard regression problem with a dataset:

$$\left\{\left(\mathbf{y}_i,\ f_{NN}(\mathbf{y}_i, \boldsymbol{\theta}^1_{NN})\right)\right\}_{i\in\mathrm{Tr}},$$

where the inputs are system states and the targets are the derivatives required to yield zero integration error. Therefore, assuming that $f_{NN}$ is an over-parameterized neural network as defined in [SGJ+21], the results from that work apply directly: the set of parameters yielding zero loss forms a connected manifold in parameter space.

**b. Extension of the proof of mode connectivity to UDEs with known mechanistic parameters.** The general UDE case can be represented by the equation

$$\frac{d\mathbf{y}}{dt} = f_M(\mathbf{y}, \boldsymbol{\theta}_M) + f_{NN}(\mathbf{y}, \boldsymbol{\theta}_{NN}), \tag{4}$$

where $f_M$ is the mechanistic component of the model, $\boldsymbol{\theta}_M \in \mathbb{R}^l$ denotes its parameters, $f_{NN}$ is an $L$-layer neural network, and $\boldsymbol{\theta}_{NN} \in \mathbb{R}^k$ are the corresponding neural network parameters. In this paragraph, we assume that $\boldsymbol{\theta}_M$ are known and fixed. In this case, under the same hypothesis on the training set $\{(t_i, \mathbf{y}_i)\}_{i\in\mathrm{Tr}}$ done in the case (a), the proof can be easily reconducted to the previous one. Let us suppose that two parameterizations of the model, $(\boldsymbol{\theta}_M, \boldsymbol{\theta}^1_{NN})$ and $(\boldsymbol{\theta}_M, \boldsymbol{\theta}^2_{NN})$, achieve zero training loss. That is,

$$\int_0^{t_i} \left(f_M(\mathbf{y}, \boldsymbol{\theta}_M) + f_{NN}(\mathbf{y}, \boldsymbol{\theta}^1_{NN})\right)\, dt = \mathbf{y}_i \quad \text{and} \quad \int_0^{t_i} \left(f_M(\mathbf{y}, \boldsymbol{\theta}_M) + f_{NN}(\mathbf{y}, \boldsymbol{\theta}^2_{NN})\right)\, dt = \mathbf{y}_i$$

for all $i \in \mathrm{Tr}$. By the uniqueness of the integral limit, and under the assumption of dense sampling, this implies

$$f_M(\mathbf{y}, \boldsymbol{\theta}_M) + f_{NN}(\mathbf{y}, \boldsymbol{\theta}_{NN}^1) = f_M(\mathbf{y}, \boldsymbol{\theta}_M) + f_{NN}(\mathbf{y}, \boldsymbol{\theta}_{NN}^2)$$

and therefore that

$$f_{NN}(\mathbf{y}, \boldsymbol{\theta}_{NN}^1) = f_{NN}(\mathbf{y}, \boldsymbol{\theta}_{NN}^2).$$

Therefore, assuming that $f_{NN}$ is an over-parameterized neural network as defined in [SGJ$^+$21], the results from that work apply directly as in the case (a).

**c. Discussion of mode connectivity in the general case of UDEs.** If the parameters $\boldsymbol{\theta}_M$ in Eq. (4) are unknown, proving mode connectivity requires additional assumptions on $f_M$ that fall outside the scope of this work. Nonetheless, preliminary evidence that mode connectivity may generally hold—except in rare cases— for systems biology models is provided by the work of Barreiro *et al.* [BV23]. Here, we restrict ourselves to analyzing a specific condition under which the proof can be adapted, and to highlighting some cases in which the structure of $f_M$ may prevent mode connectivity from holding.

In particular, if the parameters $\boldsymbol{\theta}_M$ are structurally globally identifiable [HOPY20]—meaning, intuitively, that their values can be uniquely determined from observations of the system dynamics—then the proof developed for NODEs can be readily adapted to this setting. Assuming the same hypothesis on the training dataset as in point (a), let us suppose that two parameterizations of the model, $(\boldsymbol{\theta}_M^1, \boldsymbol{\theta}_{NN}^1)$ and $(\boldsymbol{\theta}_M^2, \boldsymbol{\theta}_{NN}^2)$, achieve zero training loss. That is,

$$\int_0^{t_i} \left( f_M(\mathbf{y}, \boldsymbol{\theta}_M^1) + f_{NN}(\mathbf{y}, \boldsymbol{\theta}_{NN}^1) \right) dt = \mathbf{y}_i \quad \text{and} \quad \int_0^{t_i} \left( f_M(\mathbf{y}, \boldsymbol{\theta}_M^2) + f_{NN}(\mathbf{y}, \boldsymbol{\theta}_{NN}^2) \right) dt = \mathbf{y}_i$$

for all $i \in \mathrm{Tr}$. By the uniqueness of the integral limit, and under the assumption of dense sampling, this implies

$$f_M(\mathbf{y}, \boldsymbol{\theta}_M^1) + f_{NN}(\mathbf{y}, \boldsymbol{\theta}_{NN}^1) = f_M(\mathbf{y}, \boldsymbol{\theta}_M^2) + f_{NN}(\mathbf{y}, \boldsymbol{\theta}_{NN}^2)$$

for all $\mathbf{y}_i$ in the training set.

Now, by structural global identifiability of $\boldsymbol{\theta}_M$, it follows that

$$\boldsymbol{\theta}_M^1 = \boldsymbol{\theta}_M^2,$$

which implies

$$f_{NN}(\mathbf{y}, \boldsymbol{\theta}_{NN}^1) = f_{NN}(\mathbf{y}, \boldsymbol{\theta}_{NN}^2)$$

for all $\mathbf{y}_i \in \mathrm{Tr}$, and the proof follows as in the case (b).

There are cases in which the formulation of $f_M$ causes the mode connectivity property to fail; however, such cases are identified as rare in biological modeling by Barreiro *et al.* [BV23]. Generalizing the observations made in their analysis of various mechanistic models, we consider the following setting: given a parameterization $(\boldsymbol{\theta}_M^1, \boldsymbol{\theta}_{NN}^1)$ that achieves zero training loss, define the set

$$H = \left\{ \boldsymbol{\theta}_M \in \mathbb{R}^l \;\middle|\; f_M(\mathbf{y}_i, \boldsymbol{\theta}_M) = f_M(\mathbf{y}_i, \boldsymbol{\theta}_M^1) \; \forall i \in \mathrm{Tr} \right\}.$$

If $H$ is not a connected subset of $\mathbb{R}^l$, then mode connectivity does not hold. A concrete example arises when $f_M$ includes a parameter raised to an even exponent. In such cases, both positive and negative values of the parameter yield the same model output (e.g., $\theta$ and $-\theta$ both result in $\theta^2$). If these values are distinct from zero, the set $H$ consists of disjoint components—thus breaking the connectedness required for mode connectivity.

The final part of this section is dedicated to analyzing the implications of relaxing or violating one of the key assumptions underlying the proof of mode connectivity. Specifically, we have assumed that the set of time points

$$\{t_i\}_{i \in \mathrm{Tr}}$$

at which the training states of the system are observed forms a dense subset of the interval $[0, T]$. However, this assumption is generally unrealistic in real-world scenarios, where only a finite number of training samples are available. If the set $\{t_i\}_{i \in \mathrm{Tr}}$ is not dense in $[0, T]$, then we can no longer conclude that Eq. (2) implies

Eq. (3). In this setting, multiple distinct dynamical systems—i.e., vector fields with different parameterizations and derivatives—may still yield zero training loss by matching the training data at the observed time points only. To formalize this, let us define the zero-loss set:

$$L_0 = \left\{ \boldsymbol{\theta}_{NN} \in \mathbb{R}^k \;\middle|\; \int_0^{t_i} f_{NN}(\mathbf{y}, \boldsymbol{\theta}_{NN})\, dt = \mathbf{y}_i \quad \forall i \in \mathrm{Tr} \right\}.$$

The set $L_0$ can be viewed as the disjoint union of equivalence classes, where each class consists of parameter values that yield the same integrated dynamics over the training time points. Although we cannot currently prove that $L_0$ is connected, each equivalence class corresponds to a connected submanifold of the parameter space. Therefore, $L_0$ is the (potentially disconnected) union of non-degenerate connected manifolds.

### S3 Test cases: parameters and training trajectories

In this section, we report the ground-truth parameter values used in the test cases, as well as the training trajectories used to train the ensembles.

For each numerical test case, three training trajectories are generated by numerically integrating the governing equations from different initial conditions. Each trajectory is discretized into 100 equally spaced time points (excluding the initial condition), yielding 100 samples per trajectory and a total of 300 data points per system. The equations, parameter values, integration intervals, and initial conditions used to generate the trajectories are reported below.

In addition to these numerical test cases, we also describe the parameter values and simulation settings used to generate the training data (in the cell-survival regime) and the evaluation scenario (in the cell-death regime).

#### S3.1 Lotka–Volterra

The three ground-truth trajectories were generated by numerically integrating the system equations using the parameter values $\alpha = 1.3$, $\beta = 0.9$, $\gamma = 0.8$, and $\delta = 1.8$, following [RMM$^+$20]. For this choice of parameters, the system exhibits non-chaotic periodic dynamics. The integration was performed over the time interval $[0.0, 10.0]$ using the following initial conditions:

$$\begin{aligned}
&\text{Trajectory 1:} \quad x(0) = 3.00, \quad y(0) = 1.50,\\
&\text{Trajectory 2:} \quad x(0) = 3.38, \quad y(0) = 1.38,\\
&\text{Trajectory 3:} \quad x(0) = 3.28, \quad y(0) = 1.58.
\end{aligned}$$

#### S3.2 Damped oscillator

The three ground-truth trajectories were obtained by numerically integrating the system equations using the parameter values $\alpha = 0.1$ and $\beta = 2.0$, as in [FYSP24]. For this choice of parameters, this system is non-chaotic and exhibits spiral trajectories converging toward a unique global attractor at the origin. The integration interval was $[0.0, 25.0]$, with the following initial conditions:

$$\begin{aligned}
&\text{Trajectory 1:} \quad x(0) = 1.00, \quad y(0) = 1.00\\
&\text{Trajectory 2:} \quad x(0) = 1.06, \quad y(0) = 1.20\\
&\text{Trajectory 3:} \quad x(0) = 0.98, \quad y(0) = 1.09.
\end{aligned}$$

#### S3.3 Lorenz

The trajectories were generated using the classical chaotic parameter values $\sigma = 10.0$, $r = 28.0$, and $b = \frac{8}{3}$ [GG03]. For this choice of parameters, the system exhibits sensitive dependence on initial conditions and the emergence of the well-known Lorenz attractor [Wil79]. The system was integrated over the interval $[0.0, 2.0]$ using the following initial conditions:

$$\begin{aligned}
&\text{Trajectory 1:} \quad x(0) = 1.47, \quad y(0) = -4.44, \quad z(0) = 28.71\\
&\text{Trajectory 2:} \quad x(0) = 1.43, \quad y(0) = -4.85, \quad z(0) = 30.42\\
&\text{Trajectory 3:} \quad x(0) = 1.76, \quad y(0) = -4.41, \quad z(0) = 32.08.
\end{aligned}$$

###### S3.3.1 Cell apoptosis

The model depends on nine kinetic parameters whose ground-truth values (with units) are taken from [YLRK20]:

$$\begin{aligned}
k_1 &= 2.67 \times 10^{-9} \quad \text{cell} \cdot (\text{s} \cdot \text{molecules})^{-1}, \\
k_{d1} &= 1 \times 10^{-2} \quad \text{s}^{-1}, \\
k_{d2} &= 8 \times 10^{-3} \quad \text{s}^{-1}, \\
k_3 &= 6.8 \times 10^{-8} \quad \text{cell} \cdot (\text{s} \cdot \text{molecules})^{-1}, \\
k_{d3} &= 5 \times 10^{-2} \quad \text{s}^{-1}, \\
k_{d4} &= 1 \times 10^{-3} \quad \text{s}^{-1}, \\
k_5 &= 7 \times 10^{-5} \quad \text{cell} \cdot (\text{s} \cdot \text{molecules})^{-1}, \\
k_{d5} &= 1.67 \times 10^{-5} \quad \text{s}^{-1}, \\
k_{d6} &= 1.67 \times 10^{-4} \quad \text{s}^{-1}.
\end{aligned}$$

The training trajectory was generated under cell-survival conditions using the following initial state (expressed in $10^5\times$ molecules per cell):

$$y_1(0) = 1.34, \quad y_2(0) = 1, \quad y_3(0) = 2.67, \quad y_4(0) = 0,$$

$$y_5(0) = 0, \quad y_6(0) = 0, \quad y_7(0) = 2.9 \times 10^{-2}, \quad y_8(0) = 0.$$

To evaluate the model under a cell-death scenario, simulations were performed using the same parameter values while modifying the initial condition of $y_7$ to $y_7(0) = 2.9 \times 10^{-3}$. The training dataset is generated by integrating the system over the time interval (0, 16) hours in the cell-survival condition and uniformly sampling 120 time points. Zero-mean Gaussian noise with a standard deviation equal to 5% of min-max variation of the dynamics was added the simulated trajectory to 24 mimic measurement noise.

#### S4 OOD input sets

In this section, we describe the OOD regions over which the coverage proportion of the ensembles is evaluated in the numerical test cases, for both vector field and trajectory predictions. Fig. S11 illustrates the state-space regions used to evaluate coverage on the vector field.

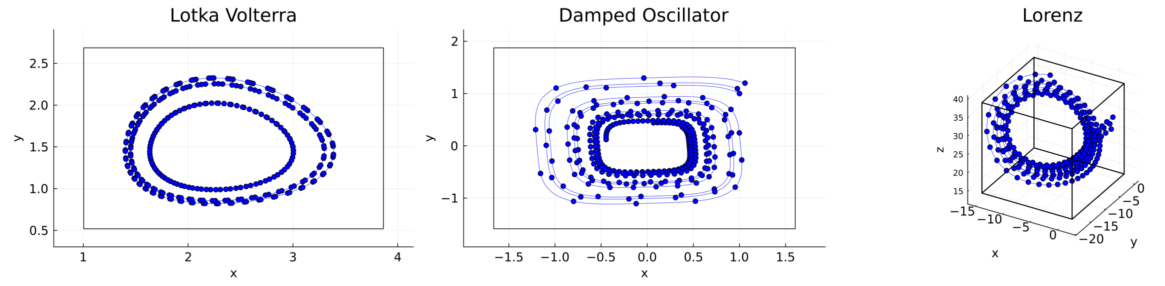


Figure S11: **Training trajectories and OOD regions for vector fields.** Phase portraits of the sampled training trajectories for the Lotka–Volterra, damped oscillator, and Lorenz systems. Black contours indicate the OOD regions used to compute uncertainty quantification metrics on the vector field.

Fig. S12 illustrates the OOD trajectories used to evaluate coverage.

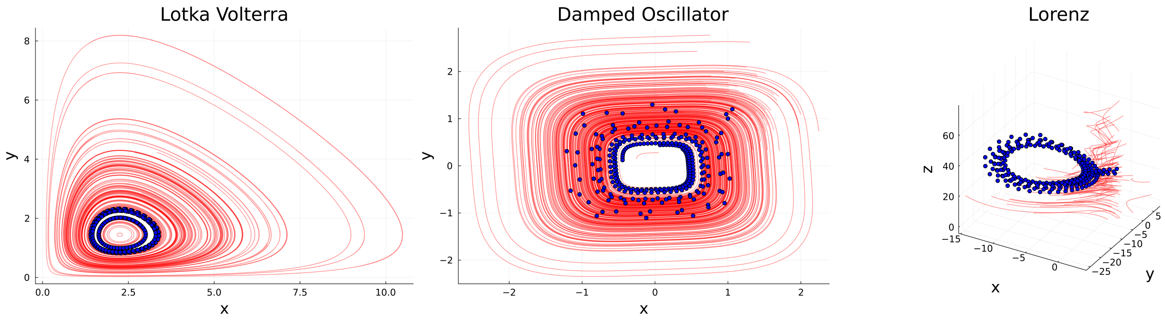


Figure S12: **Training and OOD trajectories.** The OOD trajectories used to assess model performance are shown in red for the Lotka–Volterra, damped oscillator, and Lorenz systems, while the training trajectories are shown in blue.

### S5 Partial reconstruction with known mechanistic parameters

In this scenario, we assume partial knowledge of the dynamical system, with only a portion of the system derivatives requiring reconstruction through data-driven methods.

Specifically, for the Lotka–Volterra system, we assume no knowledge of the interaction terms between the two species, and we replace them with an NN, leading to the following formulation:

$$\begin{cases} \frac{dx}{dt} = \alpha x - f_{NN}(x, y)[1] \\ \frac{dy}{dt} = f_{NN}(x, y)[2] - \gamma y \end{cases} \tag{5}$$

where $f_{NN} : \mathbb{R}^2 \to \mathbb{R}^2$ is an NN that takes $(x, y)$ as input and outputs the approximated interaction terms.

In the Damped Oscillator test case, we assume the nonlinear coupling terms between the variables to be unknown, and we replace them with an NN, yielding:

$$\begin{cases} \frac{dx}{dt} = -\alpha x^3 - f_{NN}(x, y)[1] \\ \frac{dy}{dt} = f_{NN}(x, y)[2] - \alpha y^3 \end{cases} \tag{6}$$

where, $f_{NN} : \mathbb{R}^2 \to \mathbb{R}^2$ takes $(x, y)$ as input and approximates the unknown dynamics.

In the Lorenz system, we assume that the equation defining the dynamics of the third variable of the system is entirely unknown and we substitute it with an NN, while keeping the first two equations intact:

$$\begin{cases} \frac{dx}{dt} = \sigma(y - x) \\ \frac{dy}{dt} = x(r - z) - y \\ \frac{dz}{dt} = f_{NN}(x, y, z) \end{cases} \tag{7}$$

where $f_{NN} : \mathbb{R}^3 \to \mathbb{R}$ is an NN that takes $(x, y, z)$ as input and outputs the reconstructed component of the dynamics.

In this scenario, for every test case, we assume to know the values of the parameters of the known portion of the system. The architecture of the NN used in this scenario is consistent with the one used in the first scenario, comprising two hidden layers, each containing 32 neurons, and employing the *gelu* activation function. In the case of the Lorenz system, however, the output layer consists of a single neuron, reflecting the dimensionality of the portion of the derivatives to be approximated. As in the other scenarios, for each test case, we train 10 standard ensembles and 10 corresponding MOD ensembles. The fit on the training trajectories for the models within the standard ensembles are shown in Figs. S13, S14, and S15, whereas the fits for the models within the MOD ensembles are reported in Figs. S16, S17, and S18.

The results in terms of CP for the prediction intervals on the vector fields are presented in Table S1 and are consistent with those observed in the other scenarios considered in the main text. For the Lotka–Volterra system, the standard ensembles already produce prediction intervals with a mean CP close to the nominal value of 0.95. When MOD ensembles are applied, the mean CP slightly exceeds the nominal value; in this case, the standard ensembles yield a mean CP closer to 0.95, although the absolute deviations are less than 0.05 in both cases. In contrast, MOD ensembles lead to prediction intervals with mean CP values distinctly closer to the nominal value in the Damped Oscillator and Lorenz test cases, where standard ensembles produce overconfident prediction intervals, with entire regions of the state space in which the prediction intervals fail to cover the ground truth for all trained standard ensembles (Fig. S19).

Interestingly, these results persist when the evaluation region is extended beyond the portion of the state space considered in the main analysis. To verify this, we analyze the coverage proportion of 0.95 prediction intervals for the vector field obtained using both standard and MOD ensembles, evaluated over an extended region of the state space. The extended region is defined by expanding the original bounding box—used in the main text—by 100% along each dimension. The results for the three test cases are presented in Fig. S20.

The coverage of the prediction intervals on trajectories is closely aligned with the behavior observed on the vector field (Table S2). For the Lotka–Volterra system, standard ensembles already produce prediction intervals with a mean CP close to the nominal value, and MOD ensembles result in a change that is not statistically significant. In contrast, in the Damped Oscillator and Lorenz test cases, the mean CP of the prediction intervals generated by MOD ensembles is distinctly closer to the nominal value, exceeding 0.94 in both cases.

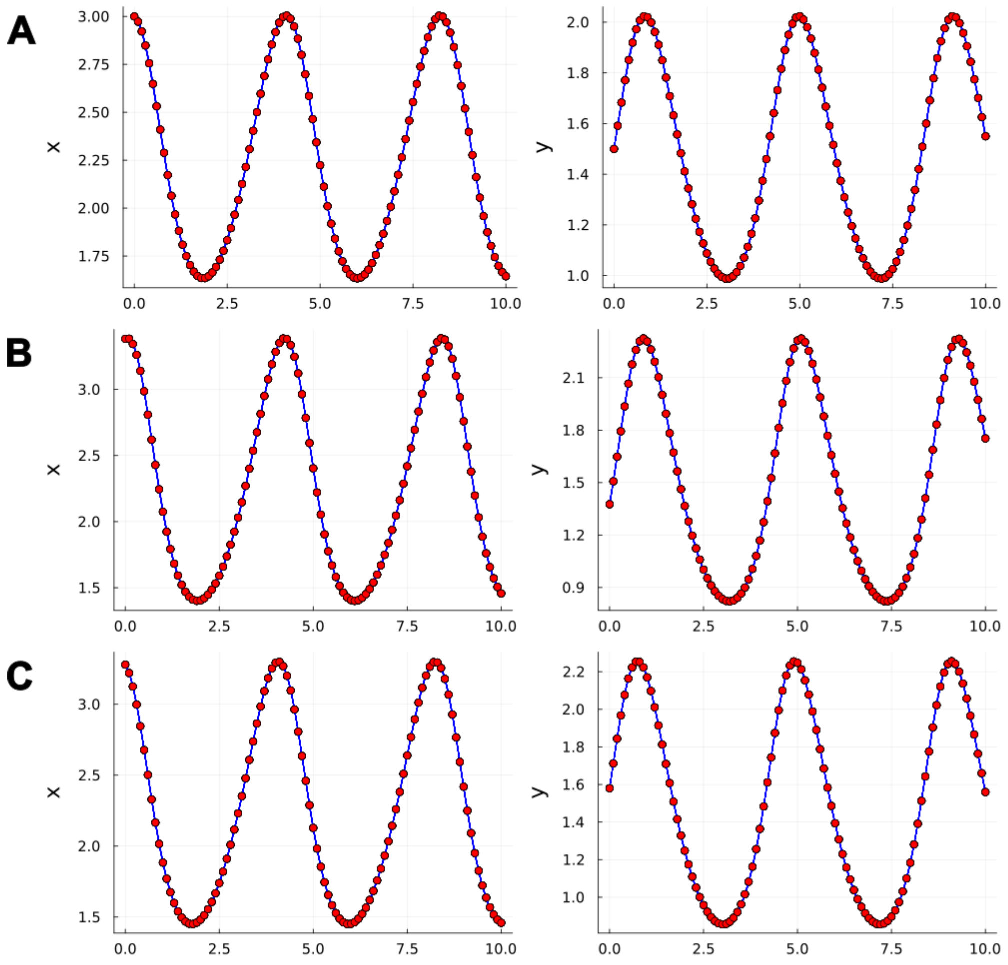


Figure S13: **Fit of trained data-driven models composing the standard ensembles on training trajectories for the Lotka–Voterra test case.** Panels A, B, and C show the fits of the trained models (blue lines) to the data points of the first, second, and third training trajectories (red points), respectively.

| | Lotka–Volterra | Damped Oscillator | Lorenz |
|---|---|---|---|
| Standard | **0.941 ± 0.018** (**) | 0.620 ± 0.014 | 0.566 ± 0.036 |
| MOD | 0.998 ± 0.001 | **0.997 ± 0.001** (**) | **0.909 ± 0.016** (**) |

Table S1: **Comparison of the CP of 0.95-prediction intervals on the vector field (partial reconstruction with known mechanistic parameters scenario).** Mean CP of 0.95-prediction intervals on the vector field within a selected region of the state space. The results are presented as the mean ± SEM across 10 ensembles. Statistical significance was assessed using the Wilcoxon signed-rank test on absolute deviations from 0.95 between the two distributions of CP values. Asterisks indicate the following significance levels: * $p \leq 0.05$, ** $p \leq 0.01$, and *** $p \leq 0.001$. The values shown in bold are those closest to the theoretical value.

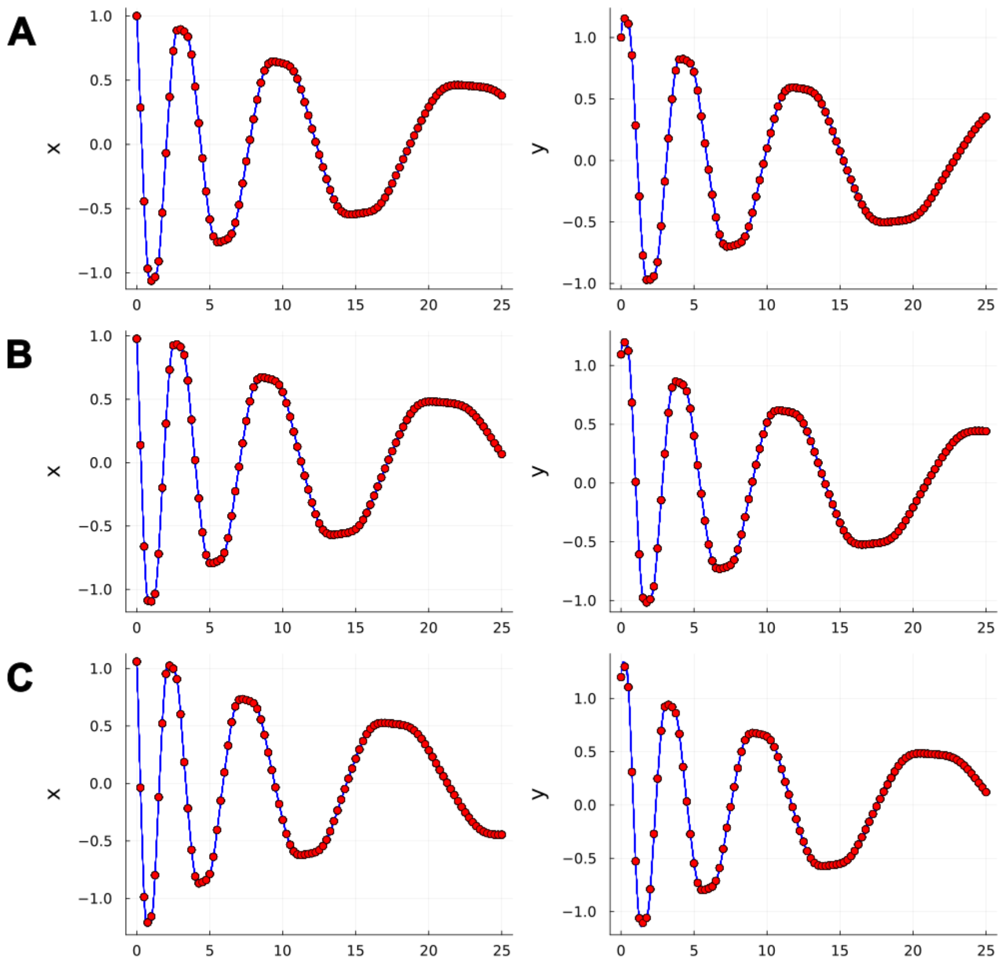


Figure S14: **Fit of trained data-driven models composing the standard ensembles on training trajectories for the Damped Oscillator test case.** Panels A, B, and C show the fits of the trained models (blue lines) to the data points of the first, second, and third training trajectories (red points), respectively.

| | **Lotka–Volterra** | **Damped Oscillator** | **Lorenz** |
|---|---|---|---|
| Standard | **0.922 ± 0.010** | 0.454 ± 0.027 | 0.811 ± 0.035 |
| MOD | 0.982 ± 0.002 | **0.946 ± 0.011** (**) | **0.950 ± 0.014** (**) |

Table S2: **Comparison of the CP of 0.95-prediction intervals on the trajectories (partial reconstruction with known mechanistic parameters scenario).** Mean CP of 0.95-prediction intervals on trajectories on the selected points of test trajectories. The results are presented as the mean ± SEM across 10 ensembles. Statistical significance was assessed using the Wilcoxon signed-rank test on absolute deviations from 0.95 between the two distributions of CP values. Asterisks indicate the following significance levels: * $p \leq 0.05$, ** $p \leq 0.01$, and *** $p \leq 0.001$. The values shown in bold are those closest to the theoretical value.

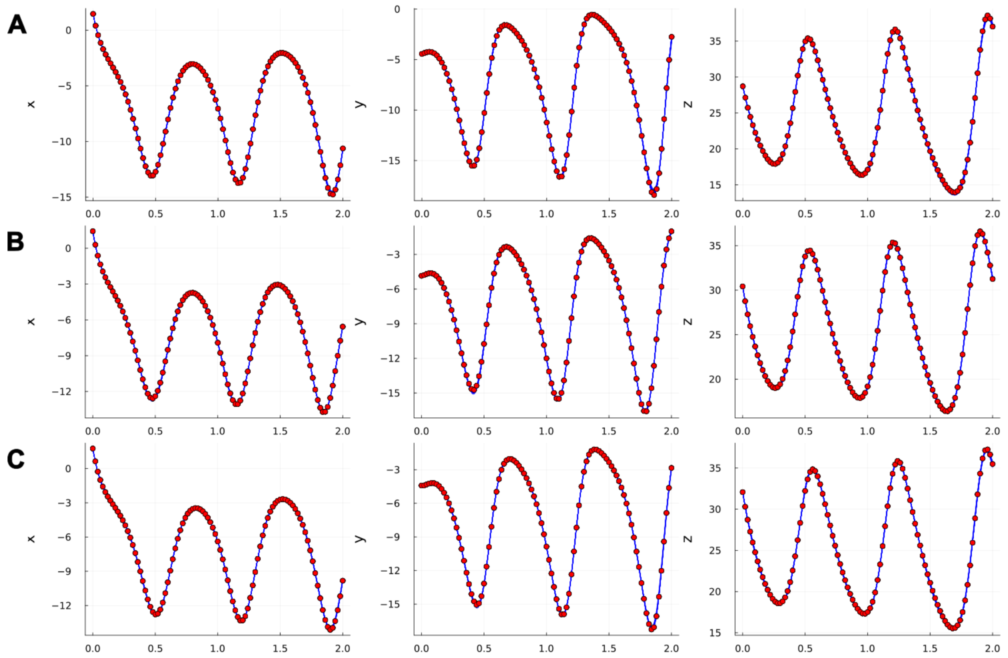


Figure S15: **Fit of trained data-driven models composing the standard ensembles on training trajectories for the Lorenz test case.** Panels A, B, and C show the fits of the trained models (blue lines) to the data points of the first, second, and third training trajectories (red points), respectively.

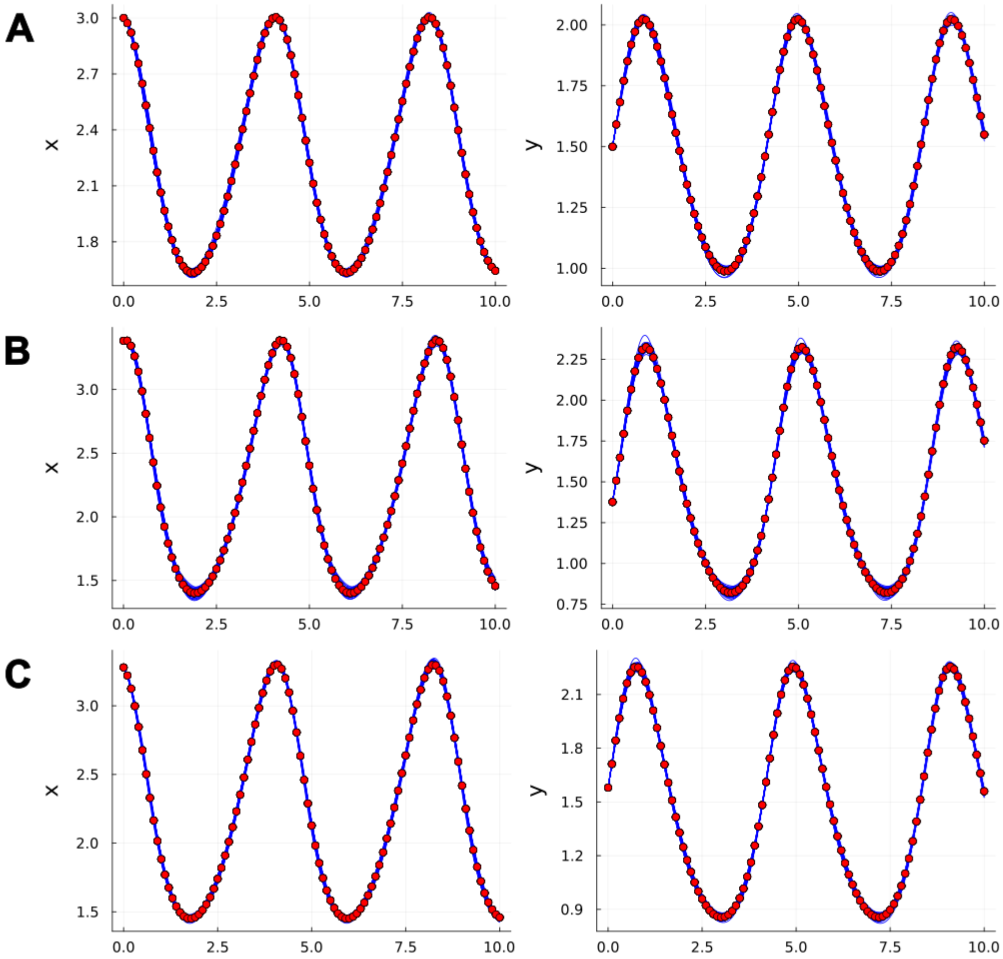


Figure S16: **Fit of trained data-driven models composing the MOD ensembles on training trajectories for the Lotka–Voterra test case.** Panels A, B, and C show the fits of the trained models (blue lines) to the data points of the first, second, and third training trajectories (red points), respectively.

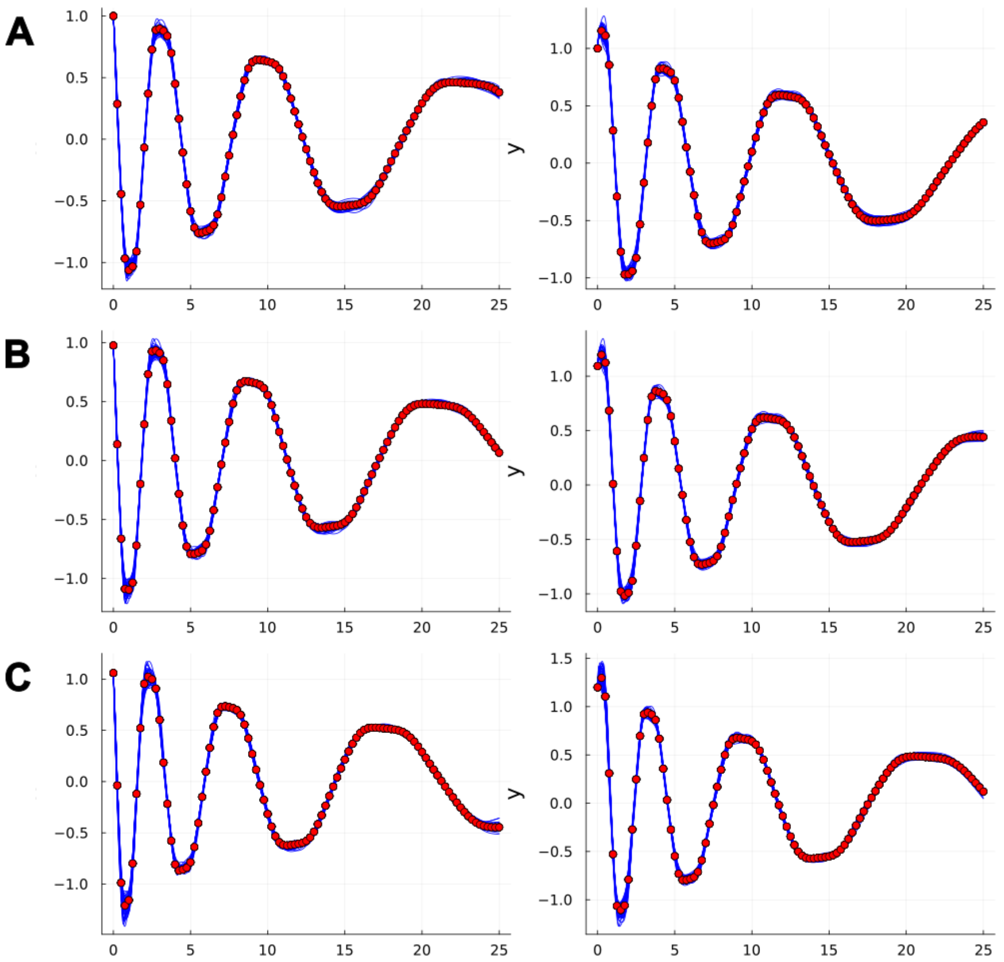


Figure S17: **Fit of trained data-driven models composing the MOD ensembles on training trajectories for the Damped Oscillator test case.** Panels A, B, and C show the fits of the trained models (blue lines) to the data points of the first, second, and third training trajectories (red points), respectively.

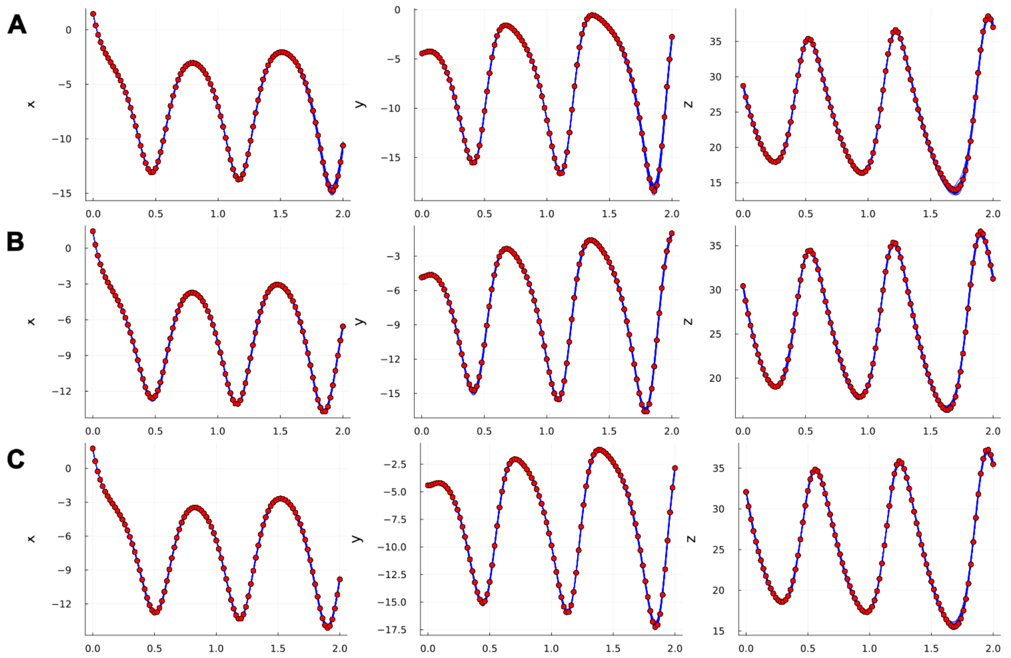


Figure S18: **Fit of trained data-driven models composing the MOD ensembles on training trajectories for the Lorenz test case.** Panels A, B, and C show the fits of the trained models (blue lines) to the data points of the first, second, and third training trajectories (red points), respectively.

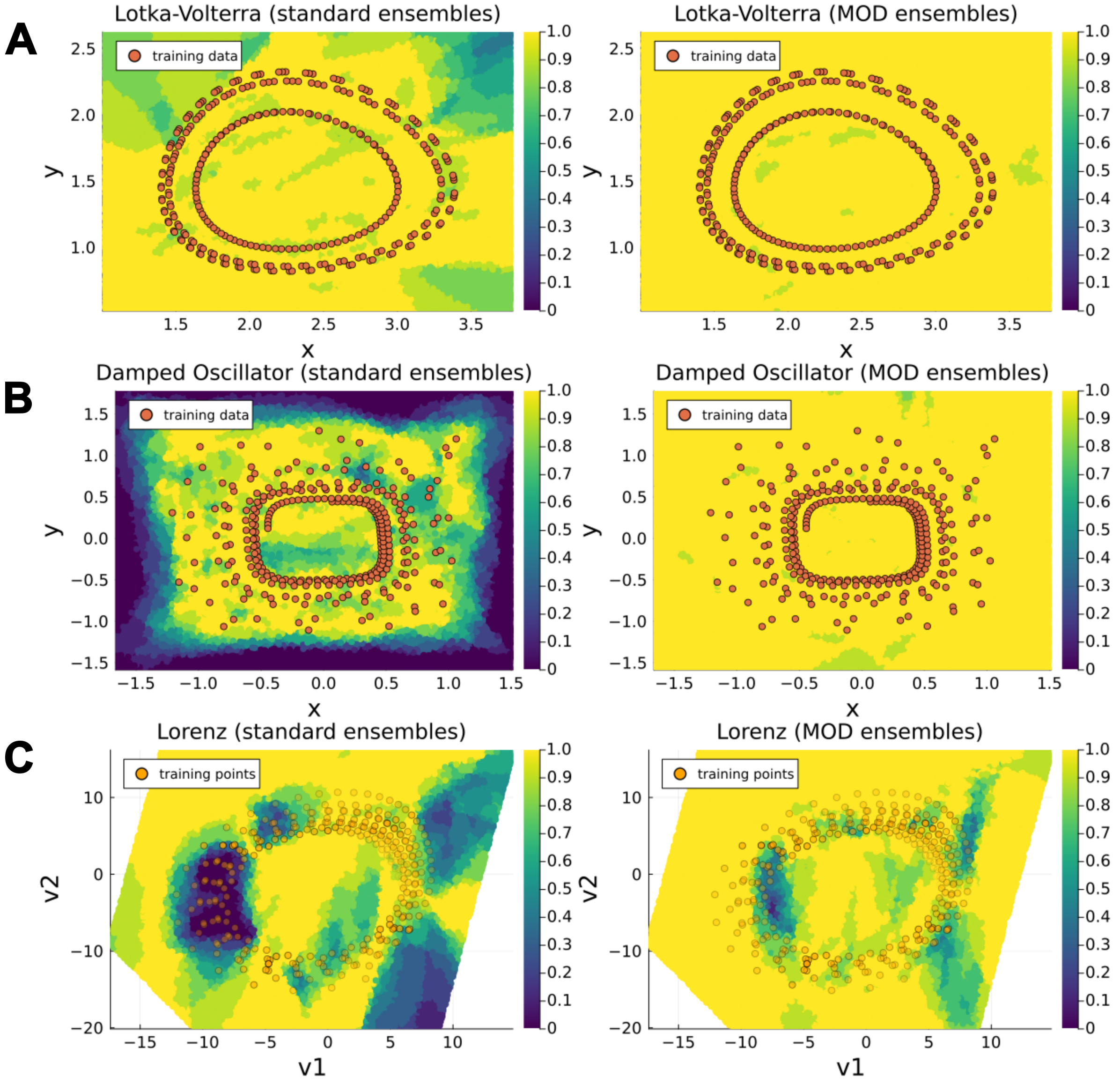


Figure S19: **Comparison of** $0.95$**-prediction interval coverage on state space: Standard vs. MOD Ensembles (partial reconstruction with known mechanistic parameters scenario).** Panels A, B, and C display heatmaps of the mean coverage (computed across 10 ensembles) of 0.95-prediction intervals on the state space for the Lotka–Volterra, Damped Oscillator, and Lorenz systems, respectively. For each system, the left subpanel shows results obtained with standard ensembles, while the right subpanel shows results obtained with MOD ensembles. Orange overlays represent the points from the training trajectories, providing spatial context. For the Lorenz system, the mean coverage is visualized on a two-dimensional plane spanned by the coordinates $(v_1, v_2)$, obtained via linear regression on the Lorenz attractor (see Supplementary Section S8 for details). These coordinates provide a low-dimensional representation of the original three-dimensional state space $(x, y, z)$ ; here, the transparency of training trajectory points indicates their orthogonal distance from the training trajectory points to the plane.

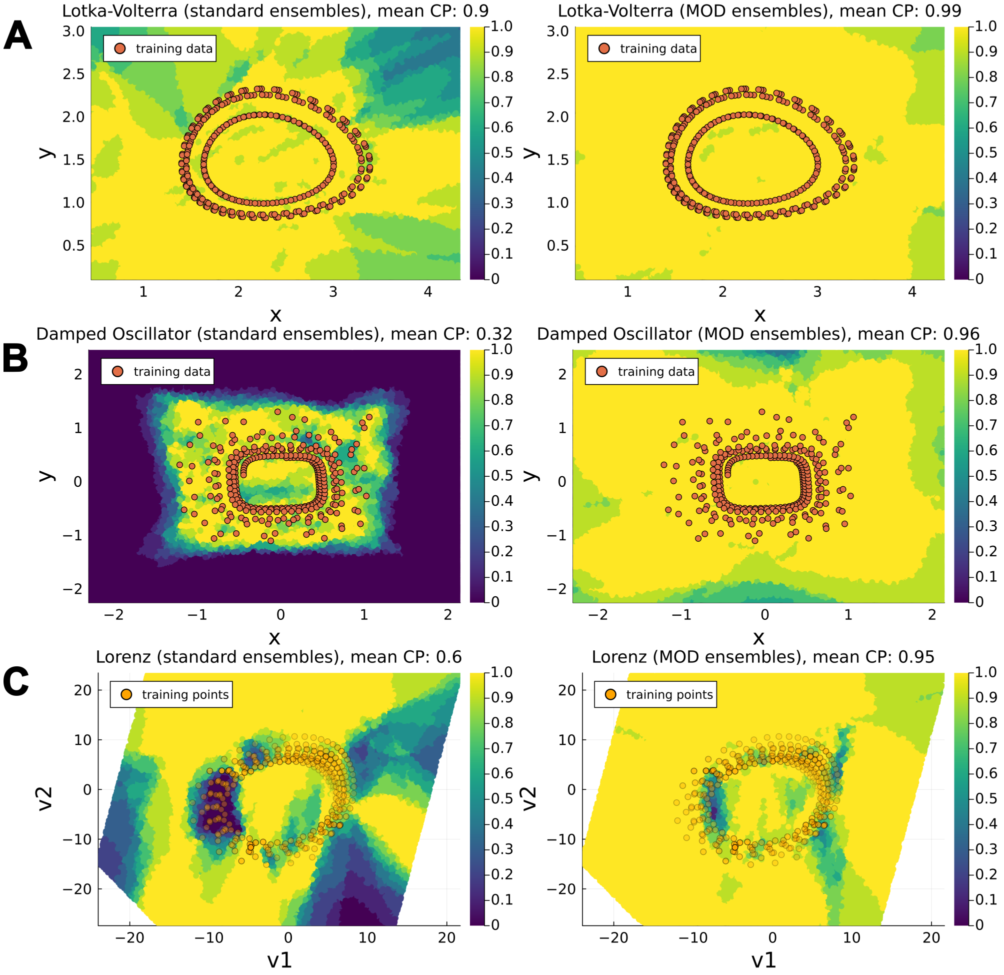


Figure S20: **Comparison of 0.95-prediction interval coverage on vector field in extended region of state space: Standard vs. MOD Ensembles.** Panels A, B, and C display heatmaps of the mean coverage (computed across 10 ensembles) of 0.95-prediction intervals on the state space for the Lotka–Volterra, Damped Oscillator, and Lorenz systems, respectively. For each system, the left subpanel shows results obtained with standard ensembles, while the right subpanel shows results obtained with MOD ensembles. Orange overlays represent the points from the training trajectories, providing spatial context. For the Lorenz system, the mean coverage is visualized on a two-dimensional plane spanned by the coordinates $(v_1, v_2)$, obtained via linear regression on the Lorenz attractor (see Supplementary Section S8 for details). These coordinates provide a low-dimensional representation of the original three-dimensional state space $(x, y, z)$ ; here, the transparency of training trajectory points indicates their orthogonal distance from the training trajectory points to the plane. The mean coverage proportion in the region is reported in the title for each plot.

### S6 Training hyperparameters for standard ensembles

In this section, we report the hyperparameters used to train the models in the standard ensembles. The overall training procedure, detailed in the **Methods** section of the main text, consists of two stages. The first stage employs a multiple shooting approach [TJ21] with the Adam optimizer (learning rate $\lambda_{\text{ADAM}}$), while the second stage fine-tunes the parameters using an L-BFGS optimizer.

In the numerical test cases, the first stage is performed for 1000 epochs (500 epochs for the cell-apoptosis test case). When both neural network and mechanistic parameters are optimized, the latter are normalized within their prescribed bounds. The second stage is performed for up to 500 epochs in the numerical test cases and 100 epochs in the cell-apoptosis test case, with early stopping based on the validation loss to prevent overfitting.

In the multiple shooting stage, each trajectory is divided into segments containing an equal number of training points ($n_{\text{points}}$). For each segment, the loss is computed as the weighted sum of two components:

1. **Fitting term** ($\mathcal{L}_{\text{fit}}$), which measures the model's accuracy in reproducing the observed data within each segment.

2. **Continuity term** ($\mathcal{L}_{\text{continuity}}$), which penalizes discontinuities between adjacent reconstructed segments.

Using the notation from the main text, let us consider the $l$-th segment, which contains the states of the system at training times $t_{l_0}, \ldots, t_{l_{n_{\text{points}}}}$. The two components of the loss for this segment are:

$$\mathcal{L}^l_{\text{fit}}(\boldsymbol{\theta}) = \frac{1}{n_{\text{points}}} \sum_{i=1}^{n_{\text{points}}} \sum_{j=1}^{n} \left( \frac{\hat{y}^j(t_{l_i}, \boldsymbol{\theta}; t_{l_0}, \mathbf{y}_{l_0}) - y^j_{l_i}}{\Gamma^j} \right)^2,$$

$$\mathcal{L}^l_{\text{continuity}}(\boldsymbol{\theta}) = \sum_{j=1}^{n} \left( \frac{\hat{y}^j(t_{n_{\text{points}}}, \boldsymbol{\theta}; t_{l_0}, \mathbf{y}_{l_0}) - y^j_{l_{n_{\text{points}}}}}{\Gamma^j} \right)^2.$$

The loss components $\mathcal{L}_{\text{fit}}(\boldsymbol{\theta})$ and $\mathcal{L}_{\text{continuity}}(\boldsymbol{\theta})$ over the entire trajectory are obtained by summing the components over all segments. The total multiple shooting loss is then given by:

$$\mathcal{L}_{\text{MS}}(\boldsymbol{\theta}) = \mathcal{L}_{\text{fit}}(\boldsymbol{\theta}) + \mu_{\text{MS}} \cdot \mathcal{L}_{\text{continuity}}(\boldsymbol{\theta}),$$

where $\mu_{\text{MS}}$ is a hyperparameter controlling the trade-off between fitting accuracy and trajectory continuity.

The hyperparameters for the different test cases and scenarios have been manually tuned and are reported in Tables S3–S5.

| | **NODE** | **UDE (fixed mechanistic parameters)** | **UDE** |
|---|---|---|---|
| $\lambda_{\text{ADAM}}$ | 0.05 | 0.005 | 0.005 |
| $n_{\text{points}}$ | 15 | 15 | 15 |
| $\mu_{\text{MS}}$ | 0.1 | 0.1 | 0.1 |

Table S3: **Hyperparameters used for training NODE and UDE models for the Lotka–Volterra system**. The table reports the hyperparameters employed for training the models in the Lotka–Volterra test case under three configurations: the purely data-driven scenario (**NODE**), the partially data-driven scenario with fixed mechanistic parameters (**UDE** with fixed mechanistic parameters), and the partially data-driven scenario with trainable mechanistic parameters (**UDE**).

| | NODE | UDE (fixed mechanistic parameters) | UDE |
|---|---|---|---|
| $\lambda_{\text{ADAM}}$ | 0.01 | 0.01 | 0.01 |
| $n_{\text{points}}$ | 5 | 5 | 5 |
| $\mu_{\text{MS}}$ | 0.1 | 0.1 | 0.1 |

Table S4: **Hyperparameters used for training NODE and UDE models for the Damped Oscillator system**. The table reports the hyperparameters employed for training the models in the Damped Oscillator test case under three configurations: the purely data-driven scenario (**NODE**), the partially data-driven scenario with fixed mechanistic parameters (**UDE** with fixed mechanistic parameters), and the partially data-driven scenario with trainable mechanistic parameters (**UDE**).

| | NODE | UDE (fixed mechanistic parameters) | UDE |
|---|---|---|---|
| $\lambda_{\text{ADAM}}$ | 0.01 | 0.01 | 0.01 |
| $n_{\text{points}}$ | 5 | 5 | 5 |
| $\mu_{\text{MS}}$ | 10.0 | 10.0 | 10.0 |

Table S5: **Hyperparameters used for training NODE and UDE models for the Lorenz system**. The table reports the hyperparameters employed for training the models in the Lorenz test case under three configurations: the purely data-driven scenario (**NODE**), the partially data-driven scenario with fixed mechanistic parameters (**UDE** with fixed mechanistic parameters), and the partially data-driven scenario with trainable mechanistic parameters (**UDE**).

| | UDE |
|---|---|
| $\lambda_{\text{ADAM}}$ | 0.005 |
| $n_{\text{points}}$ | 20 |
| $\mu_{\text{MS}}$ | 0.01 |

Table S6: **Hyperparameters used for training UDE models for the cell apoptosis test case**. The table reports the hyperparameters employed for training the models in the cell apoptosis test case under the configuration used in the main text (UDE).

#### S7 Fit on training set (full reconstruction)

In this section, we present the fits for the training trajectories of the models composing the MOD ensembles analyzed in the full reconstruction of dynamical systems. A total of 50 models were trained for each test case, consisting of five models per ensemble across 10 different ensembles. The corresponding fits for the models within the standard ensembles are discussed in Section S1. Figures S21, S22, and S23 display the fit on the training trajectories for the three test cases. Additionally, Figures S24, S25, and S26 provide a comparison of the loss values between the models in the standard ensembles and those in the MOD ensembles.

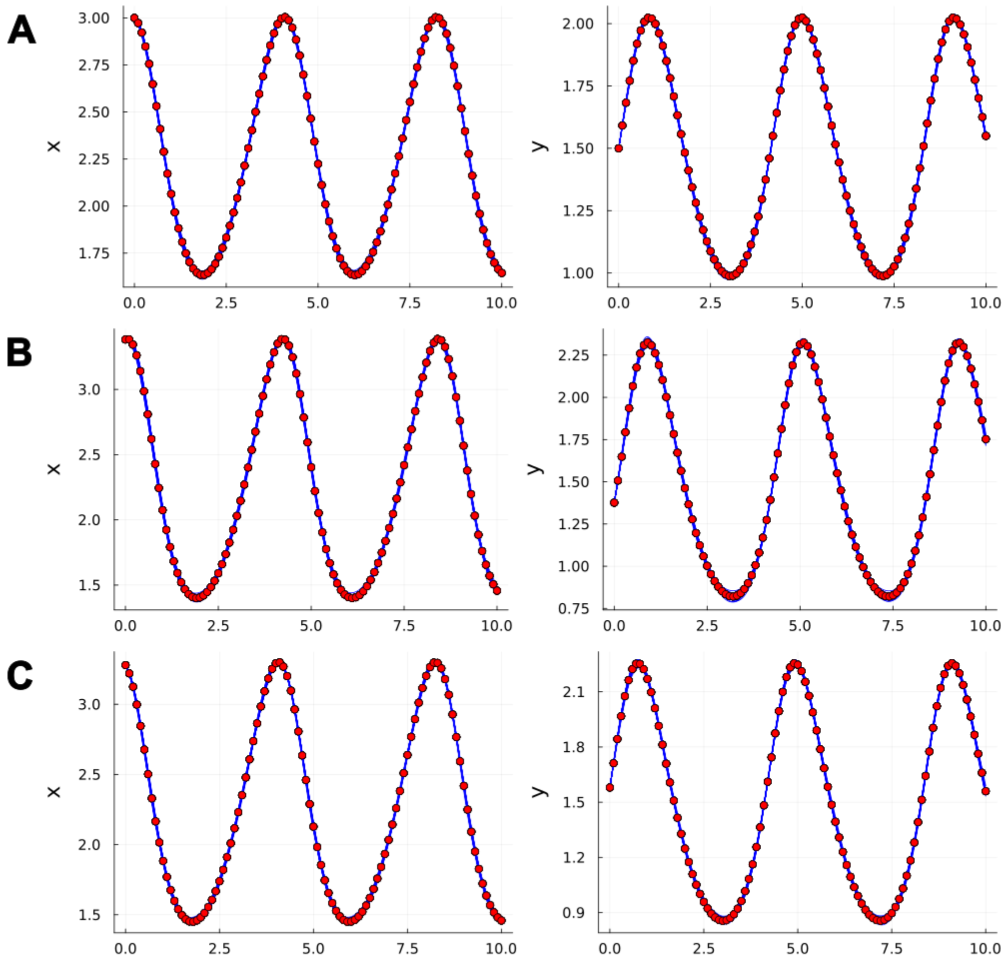


Figure S21: **Fit of trained data-driven models composing the MOD ensembles on training trajectories for the Lotka–Voterra test case.** Panels A, B, and C show the fits of the trained models (blue lines) to the data points of the first, second, and third training trajectories (red points), respectively. A total of 50 models were trained (5 per ensemble, across 10 ensembles).

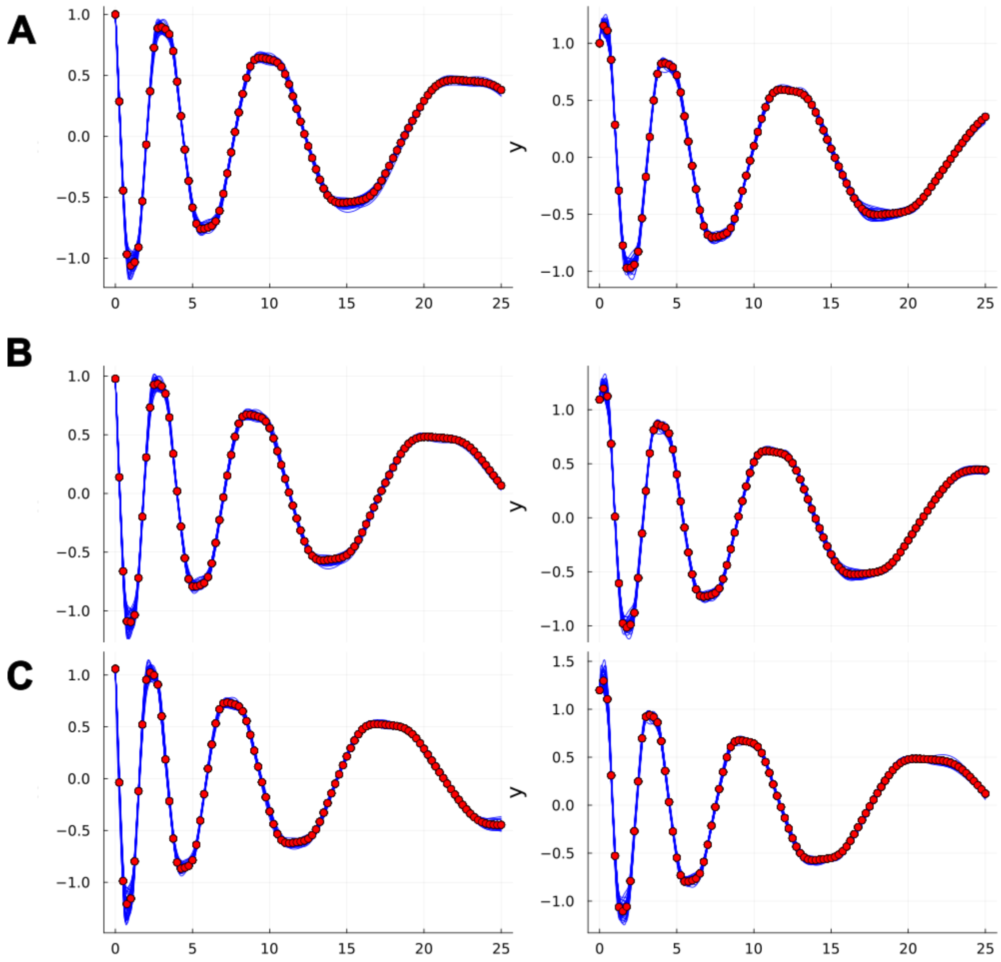


Figure S22: **Fit of trained data-driven models composing the MOD ensembles on training trajectories for the Damped Oscillator test case.** Panels A, B, and C show the fits of the trained models (blue lines) to the data points of the first, second, and third training trajectories (red points), respectively. A total of 50 models were trained (5 per ensemble, across 10 ensembles).

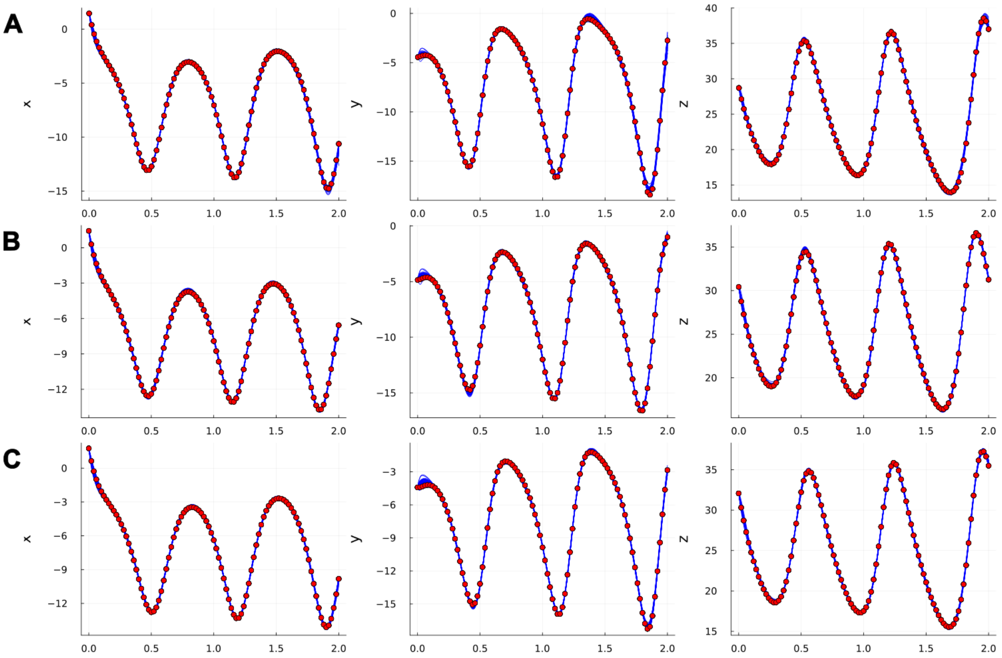


Figure S23: **Fit of trained data-driven models composing the MOD ensembles on training trajectories for the Lorenz test case.** Panels A, B, and C show the fits of the trained models (blue lines) to the data points of the first, second, and third training trajectories (red points), respectively. A total of 50 models were trained (5 per ensemble, across 10 ensembles).

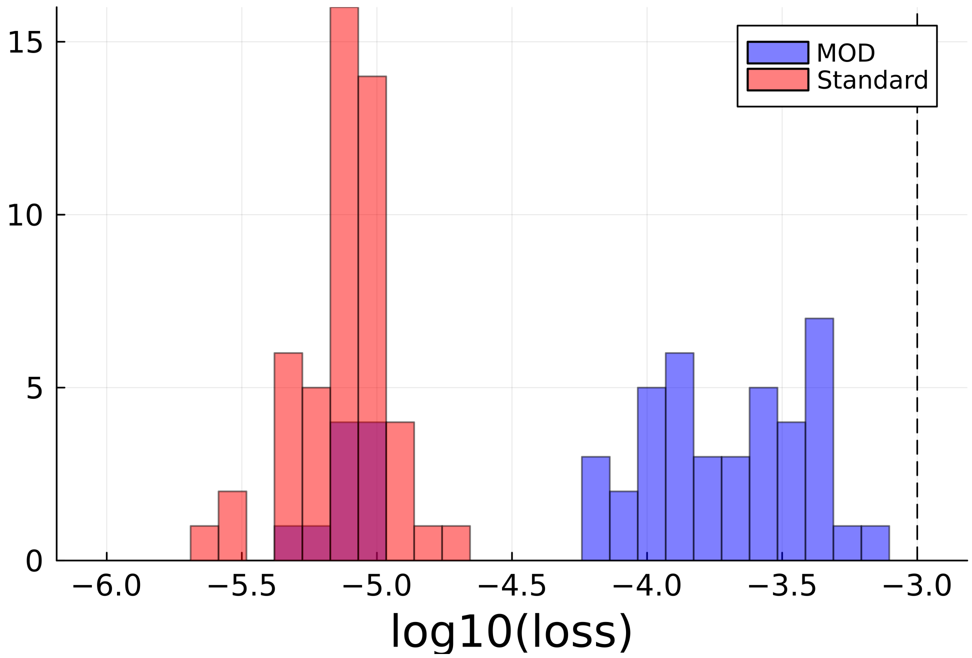


Figure S24: **Comparison among loss function values on training trajectories between models composing the standard and the MOD ensembles in the Lotka–Volterra test case.** The histogram represents the distribution of the loss function values on training trajectories achieved by the models composing the standard ensembles (red) and the MOD ensembles (blue). The dashed line represents the threshold on the loss function $\epsilon_{\text{acc}}$.

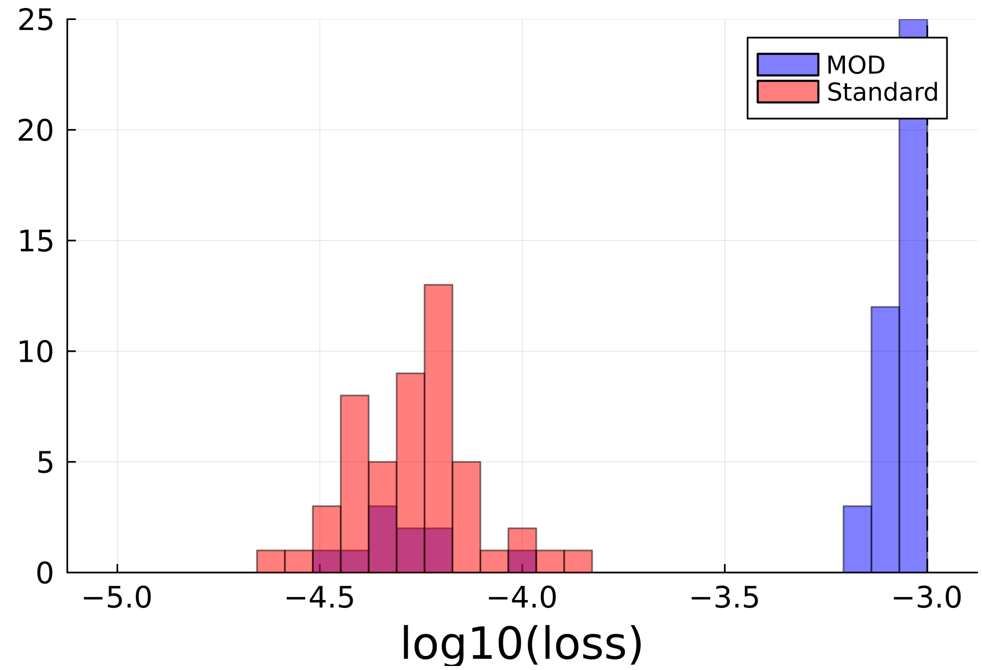


Figure S25: **Comparison among loss function values on training trajectories between models composing the standard and the MOD ensembles in the Damped Oscillator test case.** The histogram represents the distribution of the loss function values on training trajectories achieved by the models composing the standard ensembles (red) and the MOD ensembles (blue). The dashed line represents the threshold on the loss function $\epsilon_{acc}$.

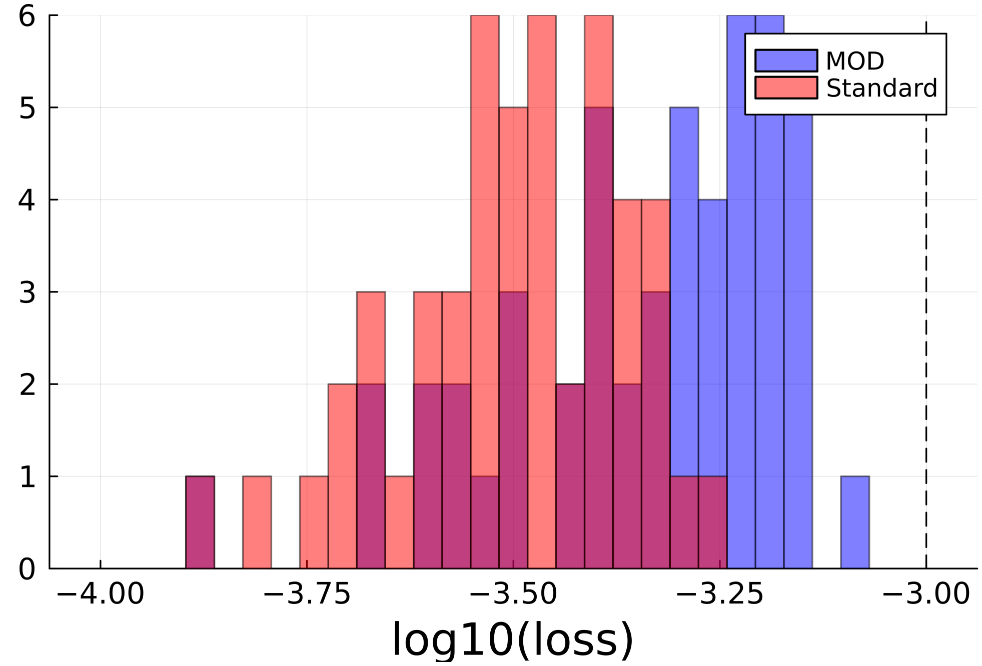


Figure S26: **Comparison among loss function values on training trajectories between models composing the standard and the MOD ensembles in the Lorenz test case.** The histogram represents the distribution of the loss function values on training trajectories achieved by the models composing the standard ensembles (red) and the MOD ensembles (blue). The dashed line represents the threshold on the loss function $\epsilon_{acc}$.

### S8 Visualization of the Lorenz system in 2D

This section describes the method used to project the training trajectories of the Lorenz system onto a two-dimensional space, enabling the visualization of a representative section of the selected region of the state space. Although the Lorenz attractor—under the considered parameter regime—evolves in a three-dimensional phase space, its structure is confined to a thin, folded set with fractal geometry. Both the Lyapunov and Hausdorff dimensions are slightly greater than 2 [DG95, Vis04], indicating that the attractor is effectively two-dimensional in spatial extent, albeit with fine-scale complexity. This near-planar geometry justifies the projection of the trajectories onto a two-dimensional subspace.

Specifically, we fit a linear regression plane to the training trajectories, considering only the portion with time greater than $t = 0.75$ to exclude the initial transient phase. The resulting plane is given by:

$$\delta : 2.64 \cdot x - 1.37 \cdot y - z = 15.48,$$

with a coefficient of determination $R^2 = 0.96$, suggesting that the fitted plane captures the dominant structure of the attractor with high fidelity. Fig. S27, Panel A displays the training trajectories and the regression plane $\delta$.

To visualize the training points in this plane, we project them using a reference frame embedded in $\delta$, with origin at

$$O = (-7.45, -7.81, 24.38)$$

and orthonormal basis vectors

$$v_1 = (0.46, 0.89, 0.0), \quad v_2 = (0.28, -0.15, -0.95).$$

Fig. S27, Panel B illustrates this reference system. The region shown in the heatmaps of mean coverage corresponds to the intersection between the regression plane and the three-dimensional region of the state space used in the subsequent analyses.

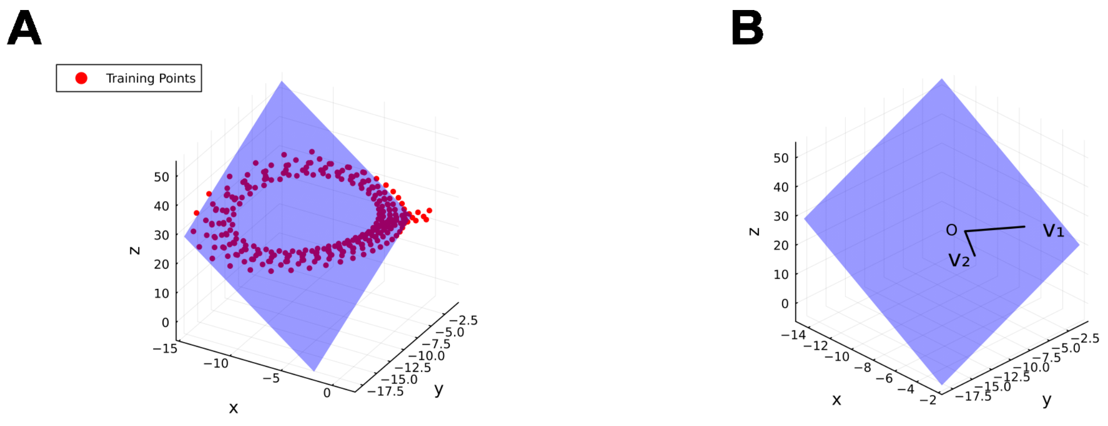


Figure S27: **Regression plane and coordinate system used to visualize the Lorenz system in 2D.** Panel A displays the training trajectories (red points) along with the regression plane $\delta$, shown in light blue. Panel B illustrates the reference frame defined by the origin $O$ and the orthonormal basis vectors $v_1$ and $v_2$, lying on the regression plane $\delta$, also rendered in light blue. The apparent difference in vector lengths is due to the non-uniform scaling of the plot axes.

### S9 Effects of considering different training datasets (full reconstruction)

In the main text, both the MOD and standard ensembles are trained on a single dataset (comprising three trajectories) for each test case. Here, we evaluate the effect of using different training datasets for ensemble training. To this end, we generate two additional sets of three trajectories each by perturbing the initial conditions of the original training trajectories with Gaussian noise (coefficient of variation 0.5). The initial points used are reported at the end of this section. We then integrate the ground-truth systems over the same time interval as in the main text and sample the trajectories at the same resolution. For each test case, we train 10 standard ensembles on each training set (each consisting of three trajectories), resulting in a total of 30 ensembles. From each ensemble, we then randomly select one model to train a corresponding MOD ensemble.

The comparison of the mean CP of the 0.95-prediction intervals on the vector field generated with the standard and MOD ensembles is reported in Tab. S7. The evaluation is carried out on the same region of the training set as described in the main text. The mean CP does not increase toward the theoretical value of 0.95 in the Lotka–Volterra test case, where the standard ensemble already achieves a mean CP close to the theoretical value. In the remaining test cases, however, the standard ensembles exhibit overconfidence, which is mitigated (with statistical significance) by the MOD ensembles. The mean coverage heatmaps are reported in Fig. S28. As in the main text, the results depend on the chosen region of the state space: when the region is extended, the overconfidence of the prediction intervals becomes evident as well (Fig. S29).

| | **Lotka–Volterra** | **Damped Oscillator** | **Lorenz** |
|---|---|---|---|
| Standard | **0.921 ± 0.013** | 0.557 ± 0.016 | 0.484 ± 0.010 |
| MOD | 0.992 ± 0.003 | **0.988 ± 0.002** (**) | **0.716 ± 0.020** (**) |

Table S7: **Comparison of the CP of 0.95-prediction intervals on the vector field (full reconstruction scenario).** Mean CP of 0.95-prediction intervals on the vector field within a selected region of the state space. The results are presented as the mean ± SEM across 10 ensembles. Statistical significance was assessed using the Wilcoxon signed-rank test on absolute deviations from 0.95 between the two distributions of CP values. Asterisks indicate the following significance levels: * $p \leq 0.05$, ** $p \leq 0.01$, and *** $p \leq 0.001$. The values shown in bold are those closest to the theoretical value.

The results in terms of the coverage of the 0.95-prediction intervals on the test trajectory are consistent with those observed on the vector field. The mean CP analysis is reported in Table S8: in this case, the mean CP of the MOD ensembles is closer to the nominal value of 0.95, with statistical significance in the Damped Oscillator and Lorenz test cases, where the standard ensembles exhibited the highest overconfidence.

| | **Lotka–Volterra** | **Damped Oscillator** | **Lorenz** |
|---|---|---|---|
| Standard | 0.905 ± 0.020 | 0.618 ± 0.019 | 0.611 ± 0.021 |
| MOD | **0.957 ± 0.008** | **0.936 ± 0.012** (***) | **0.683 ± 0.019** (**) |

Table S8: **Comparison of the CP of 0.95-prediction intervals on the trajectories (full reconstruction scenario).** Mean CP of 0.95-prediction intervals on trajectories on the selected points of test trajectories. The results are presented as the mean ± SEM across 10 ensembles. Statistical significance was assessed using the Wilcoxon signed-rank test on absolute deviations from 0.95 between the two distributions of CP values. Asterisks indicate the following significance levels: * $p \leq 0.05$, ** $p \leq 0.01$, and *** $p \leq 0.001$. The values shown in bold are those closest to the theoretical value.

**Training trajectories: Lotka–Volterra**. Integration was carried out over the time interval $[0.0, 10.0]$ with the following initial conditions:

Traj 1: $\mathbf{y}_0 = (3.00, 1.50)$ Traj 2: $\mathbf{y}_0 = (3.38, 1.38)$ Traj 3: $\mathbf{y}_0 = (3.28, 1.58)$
Traj 4: $\mathbf{y}_0 = (3.02, 1.54)$ Traj 5: $\mathbf{y}_0 = (2.82, 1.51)$ Traj 6: $\mathbf{y}_0 = (3.33, 1.26)$
Traj 7: $\mathbf{y}_0 = (2.58, 1.32)$ Traj 8: $\mathbf{y}_0 = (3.18, 1.48)$ Traj 9: $\mathbf{y}_0 = (2.81, 1.69)$

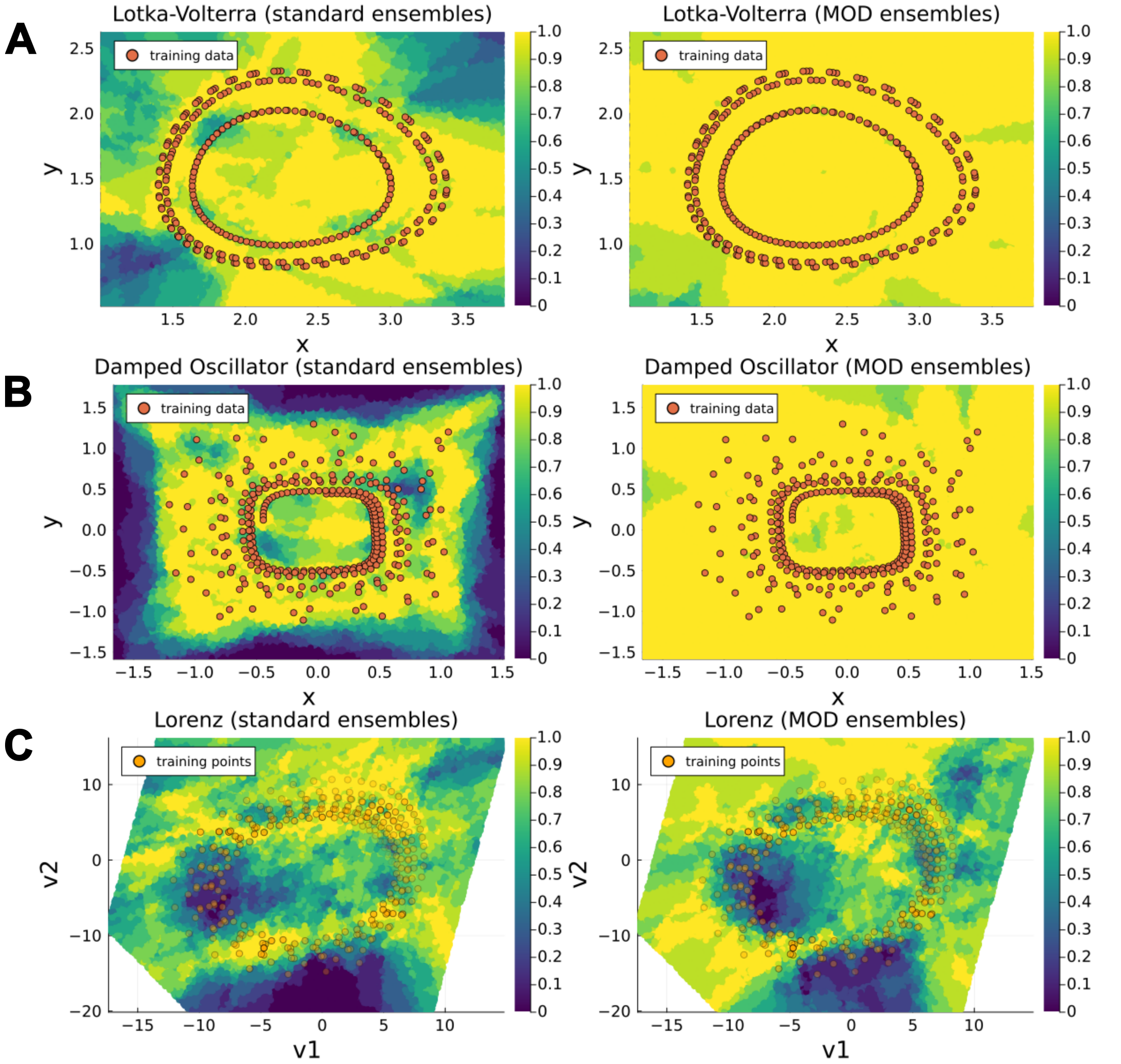


Figure S28: **Comparison of 0.95-prediction interval coverage on state space: Standard vs. MOD Ensembles (full reconstruction scenario).** Panels A, B, and C display heatmaps of the mean coverage (computed across 10 ensembles) of 0.95-prediction intervals on the state space for the Lotka–Volterra, Damped Oscillator, and Lorenz systems, respectively. For each system, the left subpanel shows results obtained with standard ensembles, while the right subpanel shows results obtained with MOD ensembles. Orange overlays represent the points from the training trajectories, providing spatial context. For the Lorenz system, the mean coverage is visualized on a two-dimensional plane spanned by the coordinates $(v_1, v_2)$, obtained via linear regression on the Lorenz attractor (see Supplementary Section S8 for details). These coordinates provide a low-dimensional representation of the original three-dimensional state space $(x, y, z)$ ; here, the transparency of training trajectory points indicates their orthogonal distance from the training trajectory points to the plane.

Ten independent ensembles were trained on Traj 1–3, ten on Traj 4–6, and ten on Traj 7–9.

**Training trajectories: Damped Oscillator**. Integration was carried out over the time interval

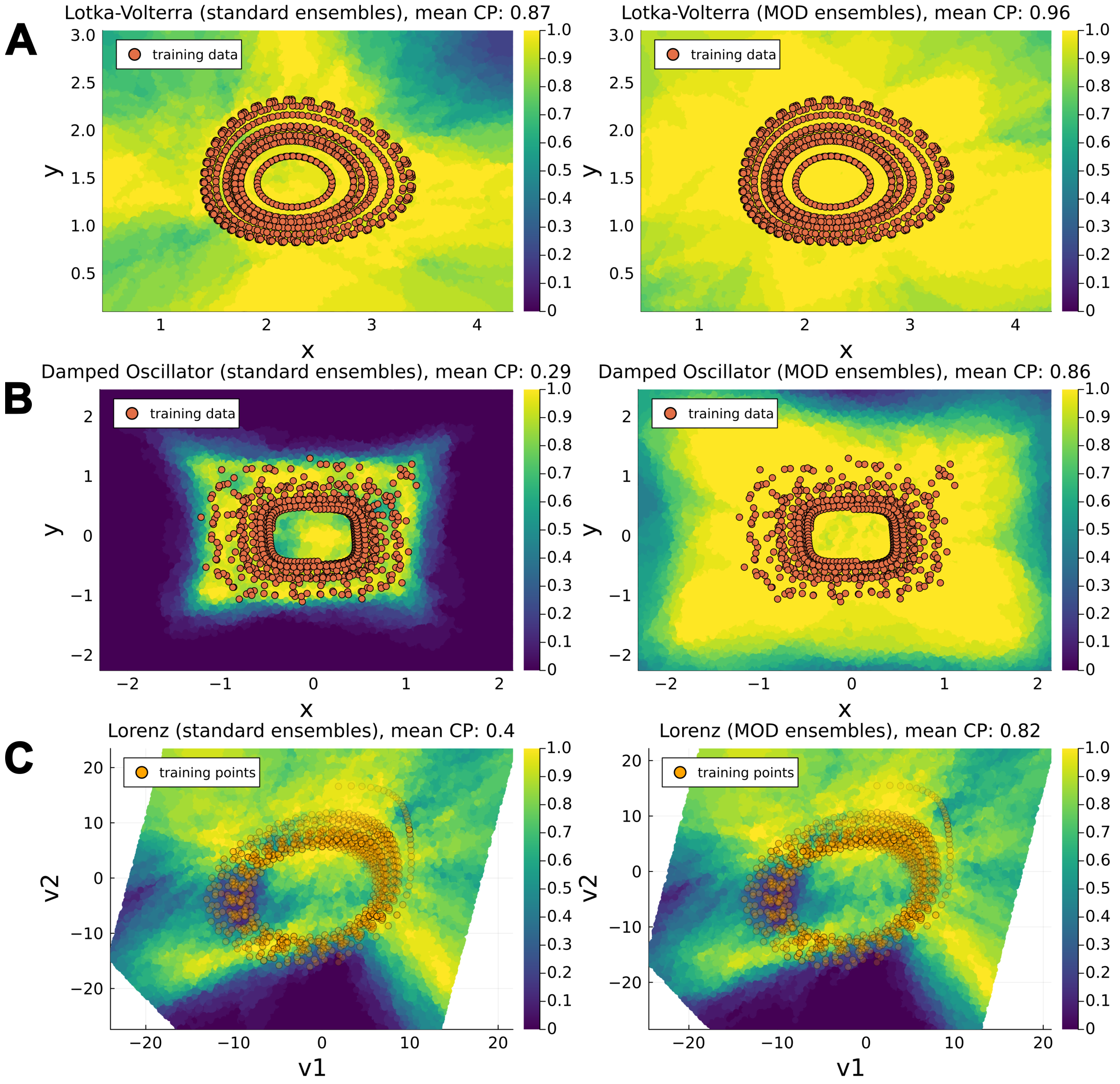


Figure S29: **Comparison of 0.95-prediction interval coverage on state space: Standard vs. MOD Ensembles (full reconstruction scenario).** Panels A, B, and C display heatmaps of the mean coverage (computed across 10 ensembles) of 0.95-prediction intervals on the state space for the Lotka–Volterra, Damped Oscillator, and Lorenz systems, respectively. For each system, the left subpanel shows results obtained with standard ensembles, while the right subpanel shows results obtained with MOD ensembles. Orange overlays represent the points from the training trajectories, providing spatial context. For the Lorenz system, the mean coverage is visualized on a two-dimensional plane spanned by the coordinates $(v_1, v_2)$, obtained via linear regression on the Lorenz attractor (see Supplementary Section S8 for details). These coordinates provide a low-dimensional representation of the original three-dimensional state space $(x, y, z)$ ; here, the transparency of training trajectory points indicates their orthogonal distance from the training trajectory points to the plane.

$[0.0, 25.0]$ with the following initial conditions:

$$\begin{array}{lll}
\text{Traj 1:} \quad \mathbf{y}_0 = (1.00, 1.00) & \text{Traj 2:} \quad \mathbf{y}_0 = (0.98, 1.09) & \text{Traj 3:} \quad \mathbf{y}_0 = (1.06, 1.20) \\
\text{Traj 4:} \quad \mathbf{y}_0 = (1.01, 1.03) & \text{Traj 5:} \quad \mathbf{y}_0 = (0.94, 1.00) & \text{Traj 6:} \quad \mathbf{y}_0 = (1.11, 0.84) \\
\text{Traj 7:} \quad \mathbf{y}_0 = (0.86, 0.88) & \text{Traj 8:} \quad \mathbf{y}_0 = (1.06, 0.99) & \text{Traj 9:} \quad \mathbf{y}_0 = (0.94, 1.13)
\end{array}$$

Ten independent ensembles were trained on Traj 1–3, ten on Traj 4–6, and ten on Traj 7–9.

**Training trajectories: Lorenz system**. Integration was carried out over the time interval $[0.0, 2.0]$, with the following initial conditions:

| | | | |
|---|---|---|---|
| Traj 1: | $\mathbf{y}_0 = (1.47, -4.44, 28.71)$ | Traj 2: | $\mathbf{y}_0 = (1.43, -4.85, 30.42)$ |
| Traj 3: | $\mathbf{y}_0 = (1.76, -4.41, 32.08)$ | Traj 4: | $\mathbf{y}_0 = (1.65, -4.07, 31.38)$ |
| Traj 5: | $\mathbf{y}_0 = (1.55, -4.48, 28.10)$ | Traj 6: | $\mathbf{y}_0 = (1.27, -4.57, 26.57)$ |
| Traj 7: | $\mathbf{y}_0 = (1.26, -3.90, 30.43)$ | Traj 8: | $\mathbf{y}_0 = (1.45, -4.15, 32.31)$ |
| Traj 9: | $\mathbf{y}_0 = (1.43, -3.35, 30.52)$ | | |

Ten independent ensembles were trained on Traj 1–3, ten on Traj 4–6, and ten on Traj 7–9.

#### S10 Comparison of prediction intervals on further sample trajectories (full reconstruction)

In this section, we present a comparison of the 0.95 prediction intervals obtained with a standard ensemble and the corresponding MOD ensemble for additional sample trajectories, complementing the results shown in Fig. 3 of the main text. The comparison is provided for the Damped Oscillator (Fig. S30) and for the Lorenz test case (Fig. S31).

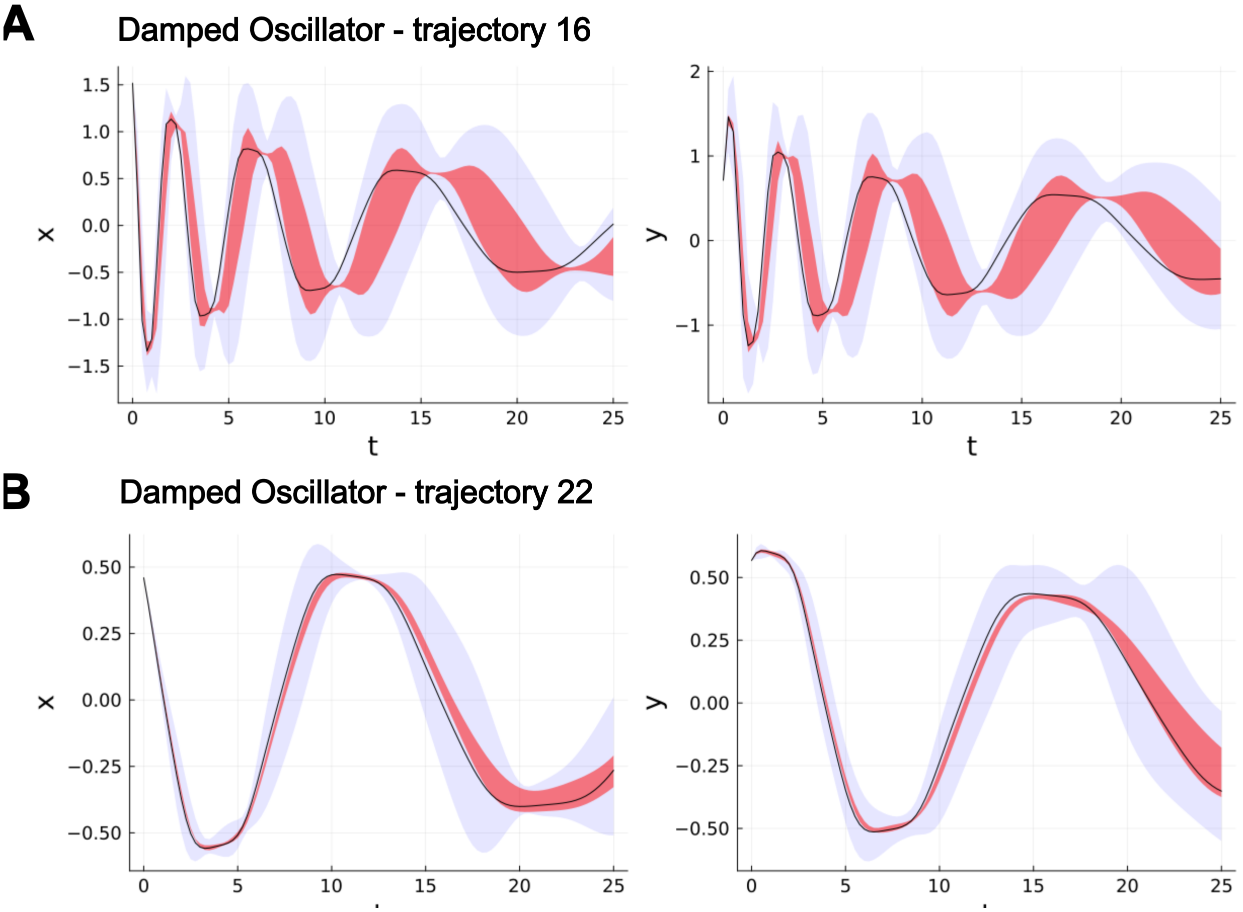


Figure S30: **Comparison between the prediction intervals on trajectories obtained with a standard ensemble and the corresponding MOD ensemble (Damped Oscillator test case).** Panels A and B show test trajectories of the Damped Oscillator (solid black line), together with the 0.95 prediction intervals of the dynamics produced by a standard ensemble (shaded red regions) and by the corresponding MOD ensemble (shaded blue regions).

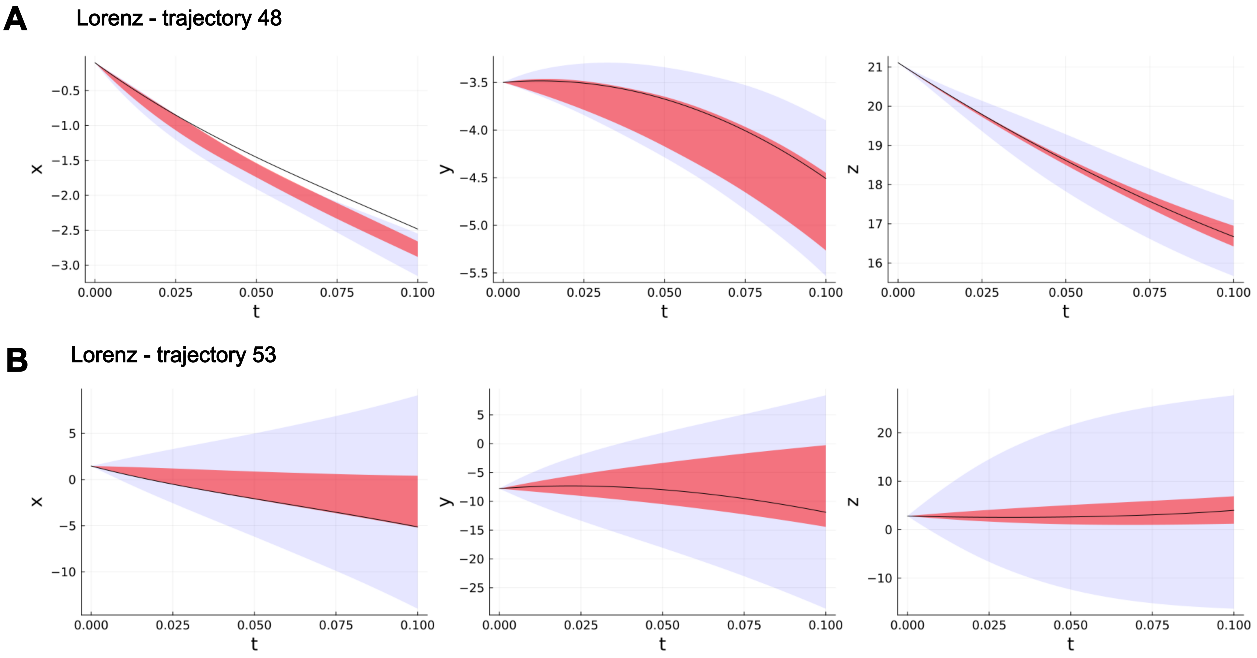


Figure S31: **Comparison between the prediction intervals on trajectories obtained with a standard ensemble and the corresponding MOD ensemble (Lorenz test case).** Panels A and B show test trajectories of the Lorenz system (solid black line), together with the 0.95 prediction intervals of the dynamics produced by a standard ensemble (shaded red regions) and by the corresponding MOD ensemble (shaded blue regions).

### S11 Fit on training set (partial reconstruction with unknown mechanistic parameters)

In this section, we present the fits for the training trajectories of the partial data-driven models (UDEs) within both the MOD ensembles and the standard ensembles. A total of 50 models were trained for each test case, consisting of five models per ensemble across 10 different ensembles. The corresponding fits for the models within the standard ensembles are shown in Figs. S32, S33, and S34, whereas the fits for the models within the MOD ensembles are reported in Figs. S35, S36, and S37.

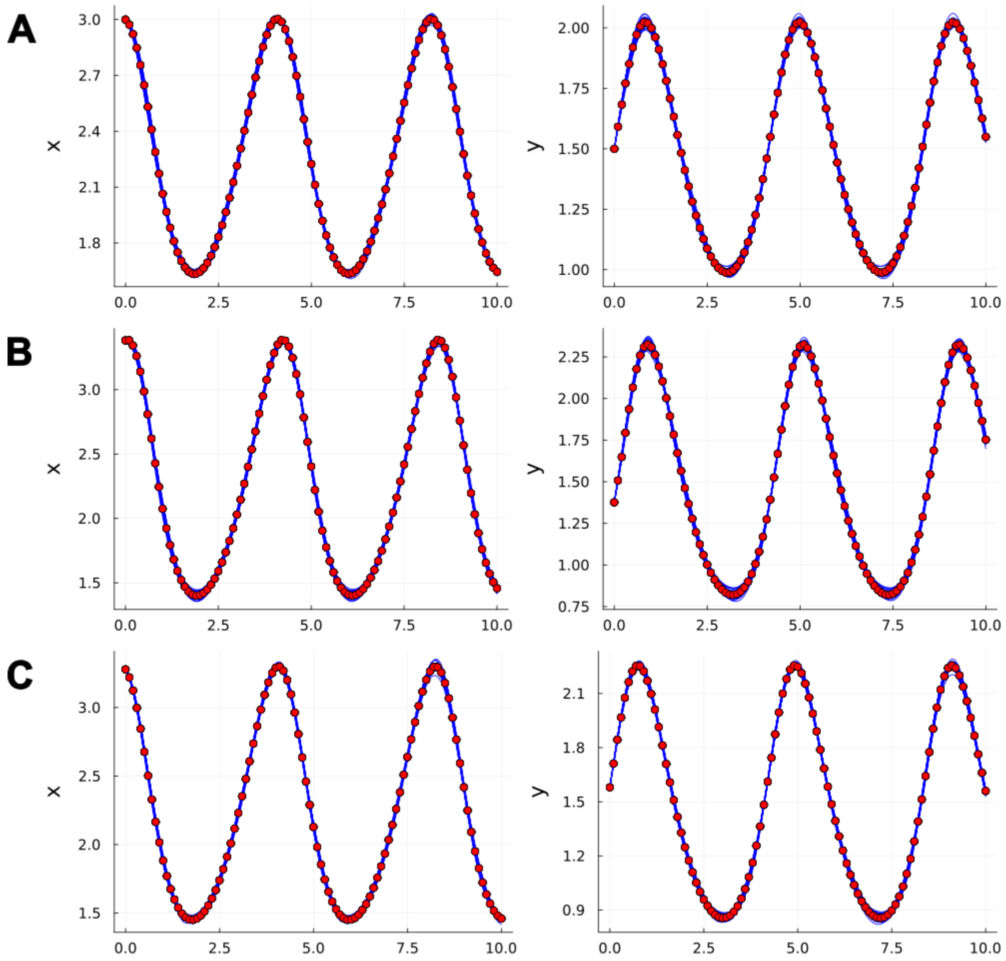


Figure S32: **Fit of trained data-driven models composing the standard ensembles on training trajectories for the Lotka–Voterra test case.** Panels A, B, and C show the fits of the trained models (blue lines) to the data points of the first, second, and third training trajectories (red points), respectively.

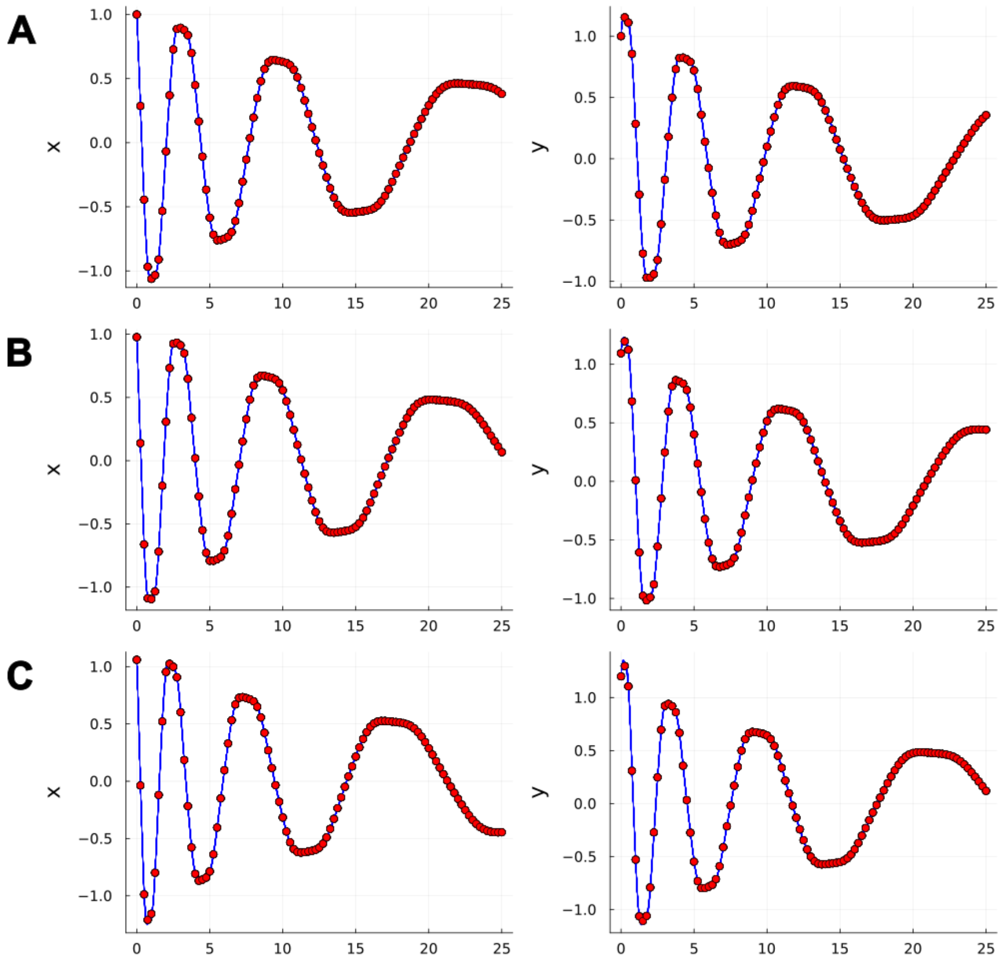


Figure S33: **Fit of trained data-driven models composing the standard ensembles on training trajectories for the Damped Oscillator test case.** Panels A, B, and C show the fits of the trained models (blue lines) to the data points of the first, second, and third training trajectories (red points), respectively.

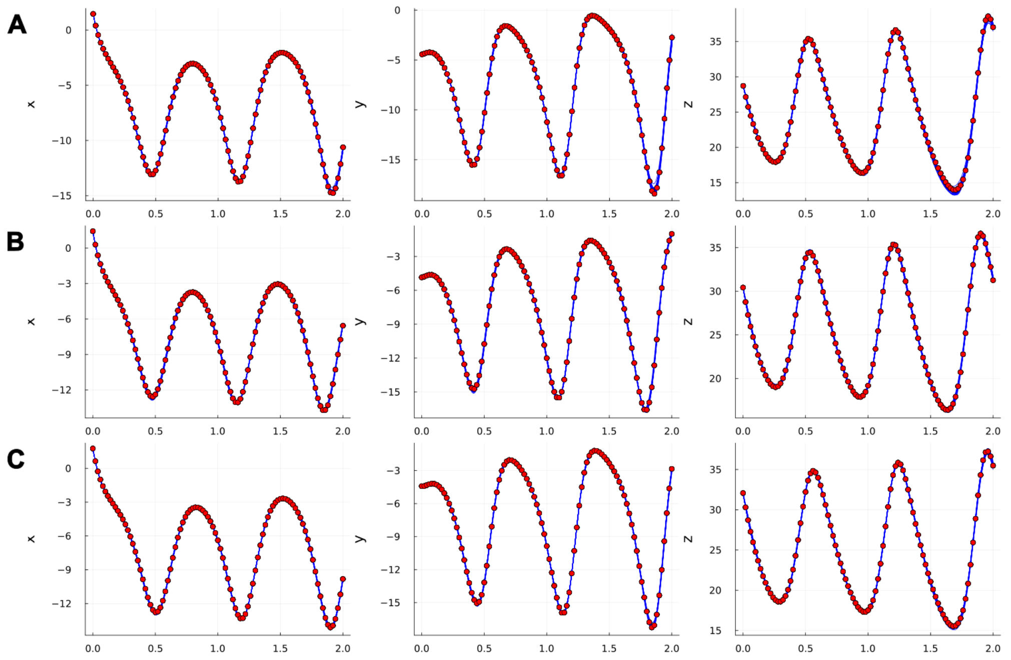


Figure S34: **Fit of trained data-driven models composing the standard ensembles on training trajectories for the Lorenz test case.** Panels A, B, and C show the fits of the trained models (blue lines) to the data points of the first, second, and third training trajectories (red points), respectively.

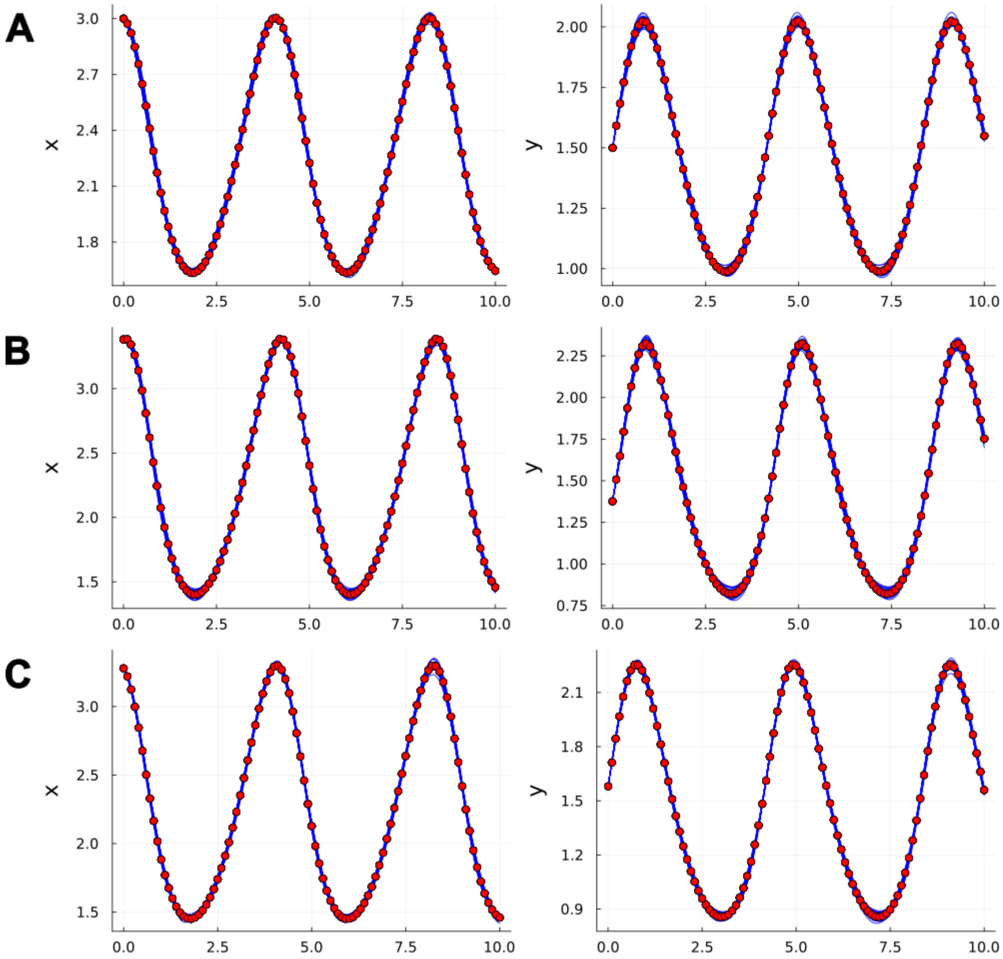


Figure S35: **Fit of trained data-driven models composing the MOD ensembles on training trajectories for the Lotka–Voterra test case.** Panels A, B, and C show the fits of the trained models (blue lines) to the data points of the first, second, and third training trajectories (red points), respectively.

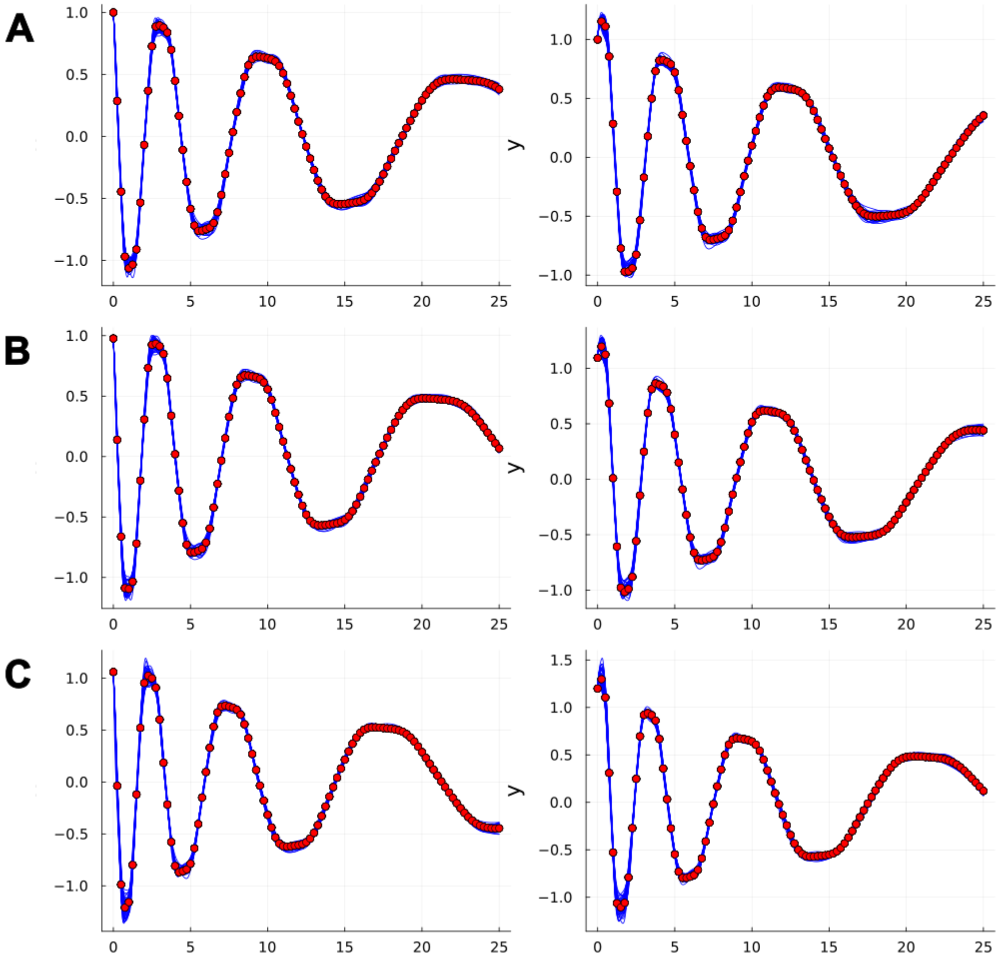


Figure S36: **Fit of trained data-driven models composing the MOD ensembles on training trajectories for the Damped Oscillator test case.** Panels A, B, and C show the fits of the trained models (blue lines) to the data points of the first, second, and third training trajectories (red points), respectively.

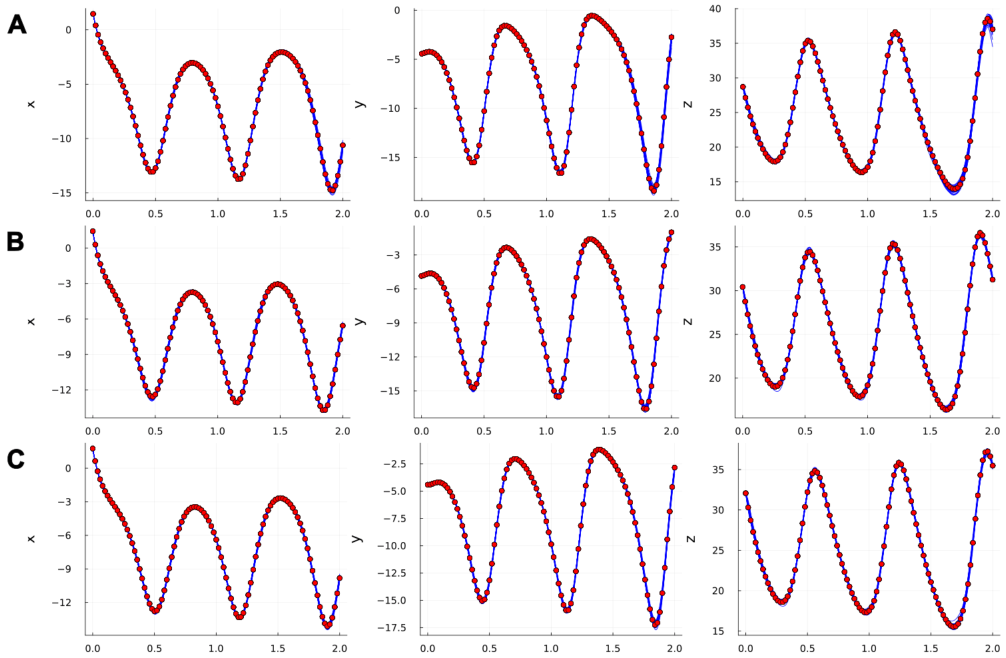


Figure S37: **Fit of trained data-driven models composing the MOD ensembles on training trajectories for the Lorenz test case.** Panels A, B, and C show the fits of the trained models (blue lines) to the data points of the first, second, and third training trajectories (red points), respectively.

### S12 Impact of mechanistic parameter identifiability on MOD ensemble training (partial reconstruction with unknown mechanistic parameters)

In this section, we examine how mechanistic parameter identifiability affects the performance of our algorithm in the partial reconstruction scenario. In the considered test cases, two distinct situations arise: the mechanistic parameters in the Lotka–Volterra UDE (both $\alpha$ and $\delta$) and in the damped oscillator UDE ($\alpha$) are non-identifiable, whereas the mechanistic parameters in the Lorenz UDE (both $\sigma$ and $r$) are identifiable.

This behavior can be informally attributed to the universal approximation properties of neural networks and can be empirically assessed either through the spread of parameter values obtained from standard ensemble training or by evaluating the Hessian of the cost function at the minima identified during training, as proposed in [GRF+24] (further details are provided at the end of this section).

By examining the trajectories of the mechanistic parameters during MOD ensemble training (Fig. S38), we observe that the algorithm explores diverse regions of the parameter space when parameters are non-identifiable (Lotka–Volterra and damped oscillator systems), whereas it remains constrained near the initial parameter values when they are identifiable (Lorenz system).

#### S12.1 Identifiability analysis of mechanistic parameters

The analysis is conducted using two distinct approaches: first, we examine the practical parameter identifiability, as described by [LDM22], by analyzing the dispersion of the parameter estimates in the trained models that compose the standard ensembles. Second, we assess the identifiability at a point [QM09] using the sensitivity-matrix-based method proposed by [GRF+24], applied to each model within the standard ensembles. These two approaches provide a comprehensive evaluation of parameter identifiability.

The distributions of the mechanistic parameters in the trained models composing the standard ensembles (models accepted with accuracy threshold $\epsilon_{\text{acc}} = 10^{-3}$) are shown in Fig. S39. In the Lotka–Volterra case, both $\alpha$ and $\delta$ are practically non-identifiable, with a coefficient of variation of 0.35 for each parameter. In the Damped Oscillator case, the mechanistic parameter $\alpha$ is practically non-identifiable, with a coefficient of variation of 0.32. In contrast, in the Lorenz system case, both parameters $r$ and $\sigma$ are practically identifiable, with coefficients of variation of 0.001 and 0.008, respectively.

The practical identifiability results obtained by analyzing the parameter distributions are further confirmed by assessing the at-a-point identifiability. This method involves analyzing the projections of the mechanistic parameters onto the null space of the Hessian matrix of the cost function (see [GRF+24] for details). Intuitively, these projections are bounded between 0 and 1, with high projections indicating non-identifiability of the parameters. The results of this analysis, performed using the same hyperparameters as in [GRF+24], are presented in Fig. S40. As with the previous analysis, they confirm the non-identifiability at a point of both mechanistic parameters in the Lotka–Volterra case, the mechanistic parameter in the Damped Oscillator case, and the at a point identifiability of both mechanistic parameters in the Lorenz system.

It is worth noting that the lack of identifiability observed for some mechanistic parameters in the Lotka–Volterra and Damped Oscillator cases arises from the insertion of the neural network into the dynamical system, rather than from intrinsic identifiability issues of the original mechanistic ODE models. To demonstrate this, we first analyzed the structural identifiability of the parameters assuming the same observable variables considered in the partial reconstruction scenario, using the method proposed in [DGHP23]. We then assessed the local (at-a-point) practical identifiability of the parameters at their literature values, reported in Supplementary Section S3, following the approach of [QM09]. The results, summarized in Table S9, show that all parameters in the considered systems are both structurally and locally practically identifiable.

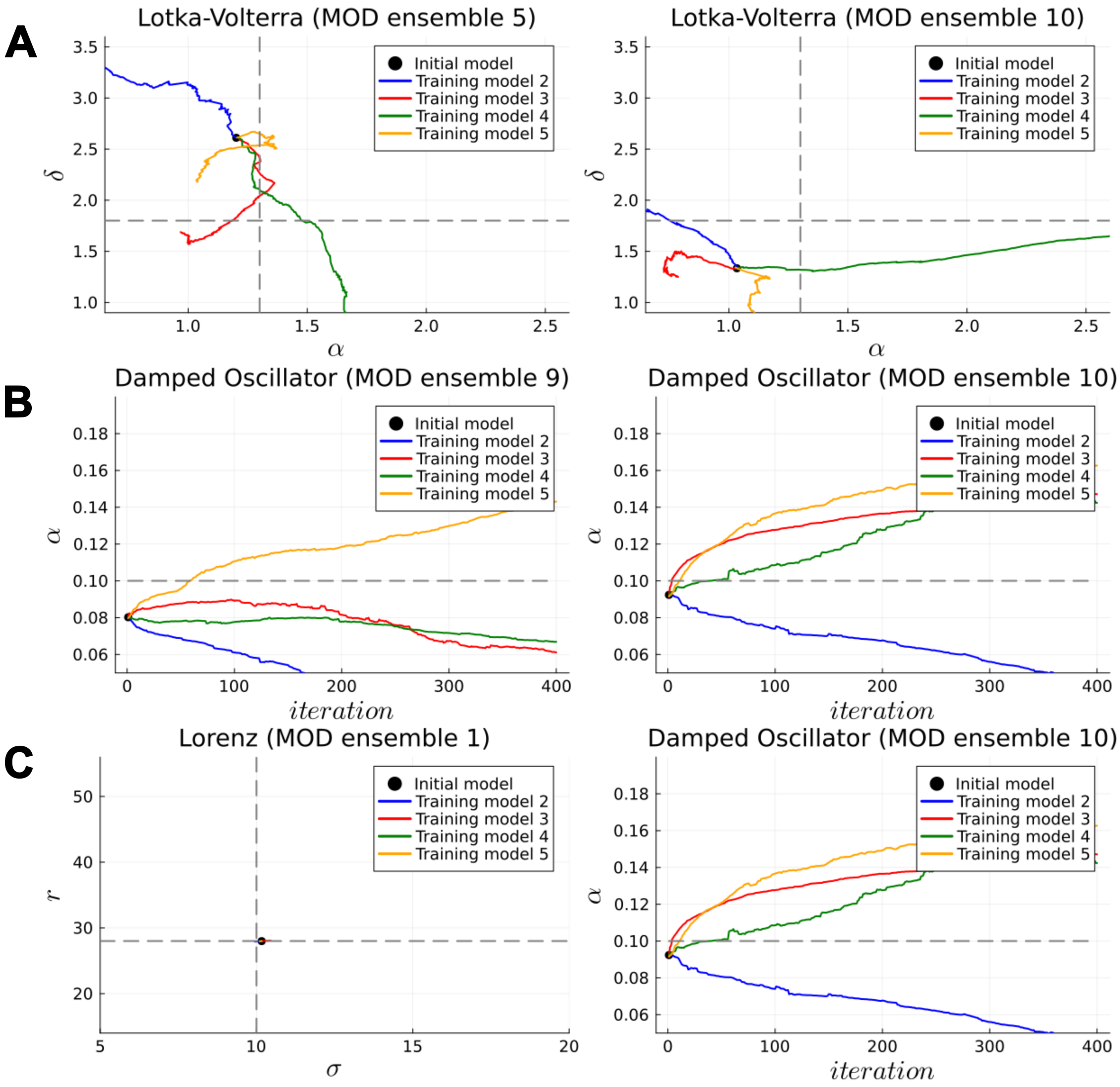


Figure S38: **Trajectories described by the mechanistic parameters during the training of models in MOD ensembles.** Panels A, B, and C show the trajectories described by the mechanistic parameters during the training of models in the MOD ensembles for the Lotka–Volterra, Damped Oscillator, and Lorenz systems, respectively. For each system, trajectories from two example ensembles are presented. The solid black point represents the mechanistic parameters of the initial model from which the algorithm begins, while the grey dashed lines indicate the ground truth values of the parameters. For the Lotka–Volterra and Lorenz systems, the trajectories are depicted in the two-dimensional space of the mechanistic parameters of the UDEs. For the Damped Oscillator, the trajectory described by the sole mechanistic parameter of the UDE is plotted against the number of iterations. The limits of the plots correspond to the boundaries assumed for the mechanistic parameter values.

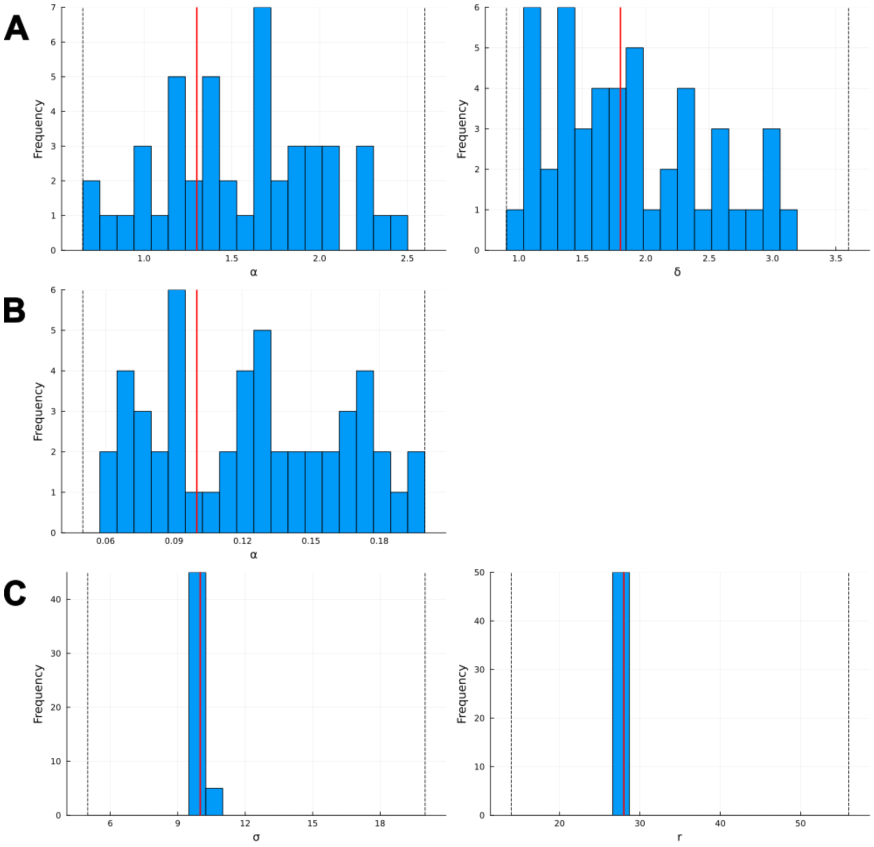


Figure S39: **Distribution of estimated mechanistic parameters across the trained UDE models.** Panels A, B, and C show the distributions of the estimated mechanistic parameters across the trained UDE models in the standard ensembles for the Lotka–Volterra system ($\alpha$ and $\delta$), the Damped Oscillator test case ($\alpha$), and the Lorenz system ($r$ and $\sigma$), respectively. The vertical red line indicates the ground-truth value, while the dashed black lines represent the boundaries of the parameter initialization.

| System | Parameter | Structurally identifiable | Locally identifiable |
|---|---|---|---|
| Lotka–Volterra | $\alpha$ | ✓ | ✓ |
| Lotka–Volterra | $\beta$ | ✓ | ✓ |
| Lotka–Volterra | $\gamma$ | ✓ | ✓ |
| Lotka–Volterra | $\delta$ | ✓ | ✓ |
| Damped oscillator | $\alpha$ | ✓ | ✓ |
| Damped oscillator | $\beta$ | ✓ | ✓ |
| Lorenz | $a$ | ✓ | ✓ |
| Lorenz | $b$ | ✓ | ✓ |
| Lorenz | $r$ | ✓ | ✓ |

Table S9: **Structural and local practical identifiability of the mechanistic parameters for the considered dynamical systems**.

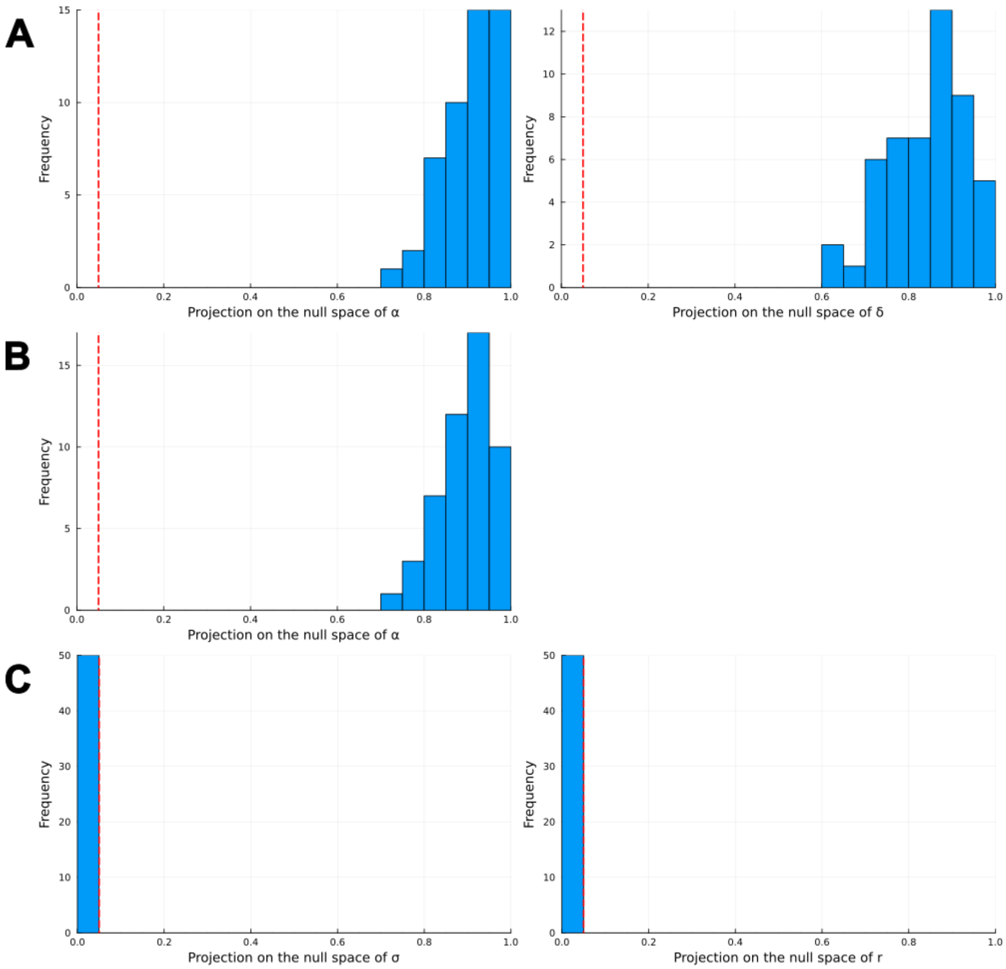


Figure S40: **Distribution of the projections of mechanistic parameters on the null space of the Hessian across trained UDE models.** Panels A, B, and C display the distributions of the projection of mechanistic parameters on the null space of the Hessian matrix across UDE models in standard ensembles for the Lotka–Volterra system (parameters $\alpha$ and $\delta$), the Damped Oscillator test case (parameter $\alpha$), and the Lorenz system (parameters $r$ and $\sigma$), respectively. The vertical red line marks the threshold used in [GRF$^+$24] to differentiate at-a-point identifiable parameters (to the left of the line) from at-a-point non-identifiable ones (to the right of the line).

### S13 Cell apoptosis test case: performance of the models on the training data

Fig. S41 reports the dynamics of $y_4$ under cell-survival training conditions simulated by the models composing the standard and MOD ensembles. All simulations achieve a training-set loss lower than 0.005.

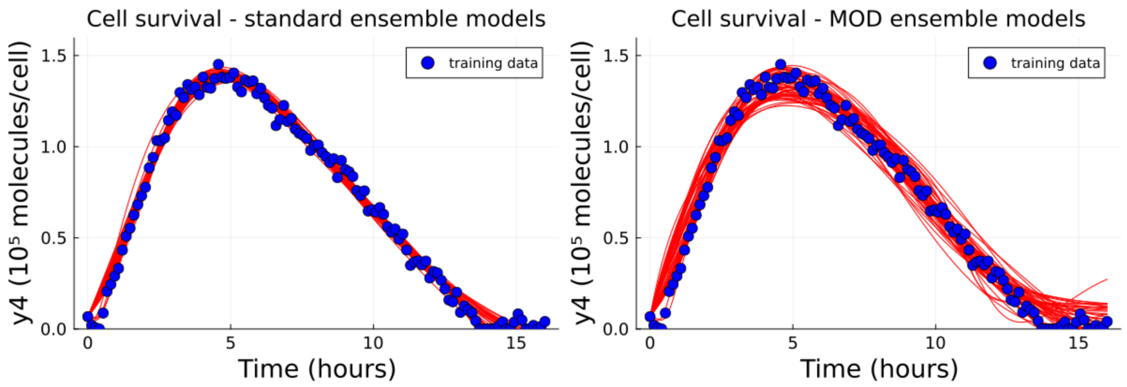


Figure S41: **Fit of models from the standard and MOD ensembles on the training data.** The plots show the dynamics of $y_4$ simulated by the models composing the standard ensembles (left) and the MOD ensembles (right) under cell-survival conditions. The training data are shown as blue points.

#### S14 Cell apoptosis test case: 0.95-prediction intervals for $y_4$ in the cell-death scenario

Fig. S42 shows the 0.95 prediction intervals for the dynamics of $y_4$ under cell-death conditions obtained with the standard and MOD ensembles for the 10 ensemble pairs.

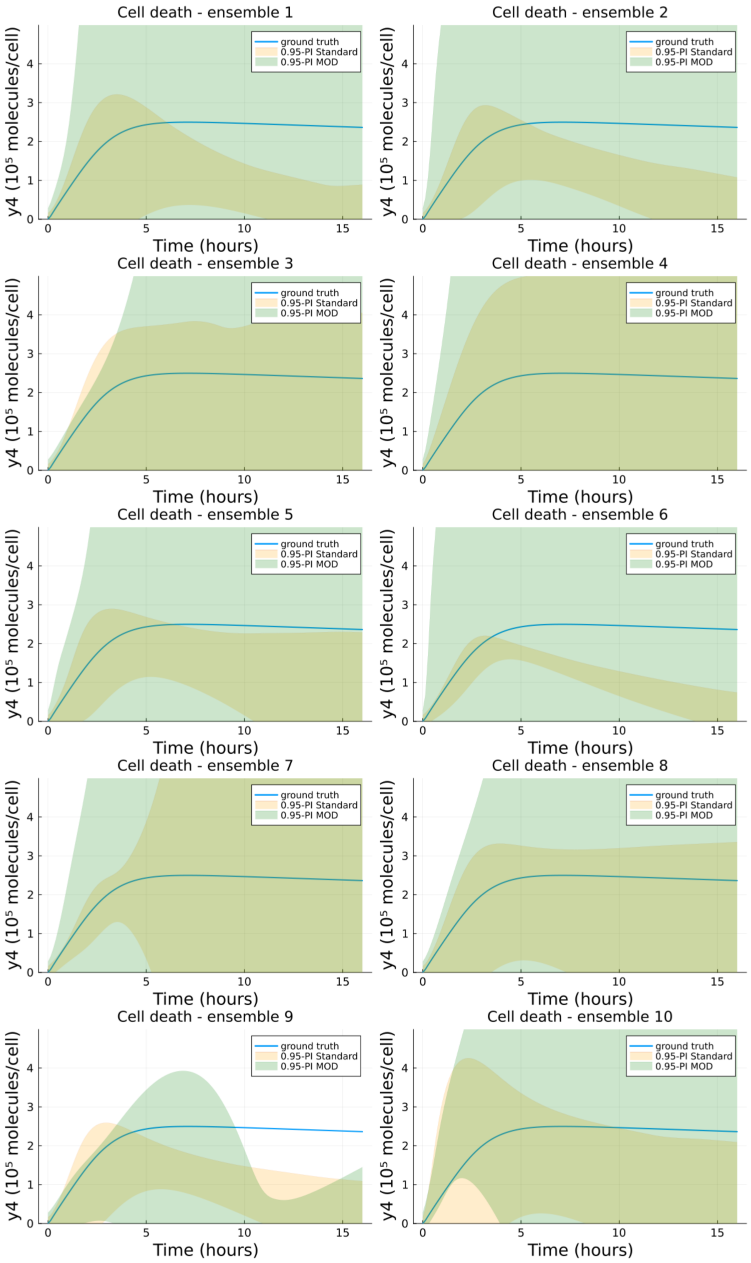


Figure S42: **0.95-prediction intervals for $y_4$ in the cell-death scenario.** The plots show the 0.95-prediction intervals obtained with standard ensembles (yellow) and MOD ensembles (green) for the 10 ensemble pairs. The blue line represents the dynamics of the ground-truth model for the variable $y_4$.

### S15 Comparison of MOD algorithm performance under different levels of prior system knowledge

In this section, we compare the training performance of MOD ensembles in terms of the distribution of coverage proportions (CP) across different levels of prior system knowledge. For each scenario considered—full reconstruction, partial reconstruction, and partial reconstruction with known mechanistic parameters—we examine the CP distributions obtained from the 10 different MOD ensembles presented in the main text. Increasing levels of prior knowledge are generally expected to reduce uncertainty in the system behavior; however, the extent to which this expectation is reflected in practice depends on the specific test case. The comparisons of CP values for vector field reconstruction and out-of-distribution trajectory reconstruction shown in Figs. S43 and S44 provide qualitative evidence of this trend in some, but not all, scenarios. The out-of-distribution datasets used to evaluate CP for both vector field and trajectory reconstruction are those described in the **Methods** section of the main text.

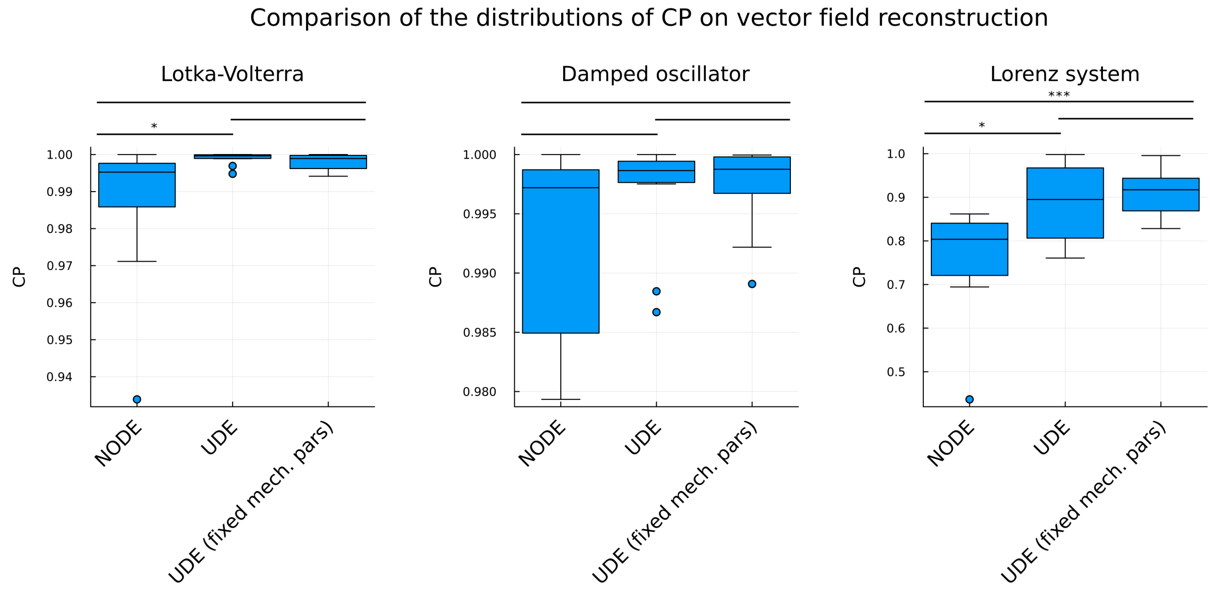


Figure S43: **Distribution of CP for vector field reconstruction using MOD ensembles.** The plots show, for each test case, a comparison of the CP distributions obtained from vector field reconstructions using MOD ensembles under different scenarios: full reconstruction (NODE), partial reconstruction (UDE), and partial reconstruction with known mechanistic parameters (UDE with fixed mechanistic parameters). The distributions are obtained by considering 10 different MOD ensembles for each scenario, as described in the **Results** section of the main text. The CP is evaluated over the same region of the state space considered in the main text and detailed in the **Methods** section. Statistical differences between distributions are assessed using the Mann–Whitney test; * indicates $p < 0.05$, ** indicates $p < 0.01$, and *** indicates $p < 0.001$.

For vector field reconstruction, an increase in the median of the CP distribution is observed when moving from full reconstruction to partial reconstruction of the system. This increase is statistically significant in the Lotka–Volterra and Lorenz test cases, while no statistically significant difference is observed in the Damped Oscillator test case. Furthermore, the comparison between partial reconstruction and partial reconstruction with known mechanistic parameters does not yield statistically significant differences in any of the test cases considered. One possible explanation for this observation is that the structural form of the model is known in both the partial reconstruction and the partial reconstruction with known mechanistic parameters scenarios and the vast majority of the model parameters is associated with the neural network component. In particular, in the Lotka–Volterra test case the neural network includes 1218 parameters, compared to 2 mechanistic parameters; in the Damped Oscillator test case, the neural network includes 1218 parameters and the mechanistic model includes 1 parameter; and in the Lorenz test case, the neural network includes

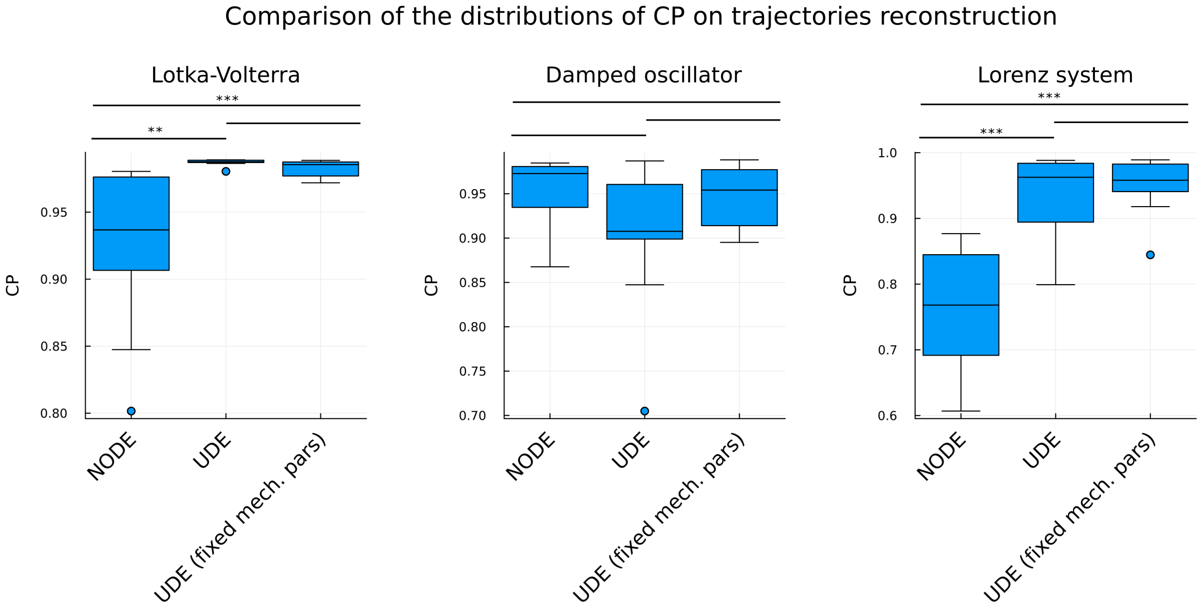


Figure S44: **Distribution of CP for trajectory reconstruction using MOD ensembles.** The plots show, for each test case, a comparison of the CP distributions obtained from trajectory reconstructions using MOD ensembles under different scenarios: full reconstruction (*Neural Ordinary Differential Equations, NODE*), partial reconstruction (*Universal Differential Equations, UDE*), and partial reconstruction with known mechanistic parameters (UDE with fixed mechanistic parameters). The distributions are obtained by considering 10 different MOD ensembles for each scenario, as described in the **Results** section of the main text. The CP is evaluated over the same trajectories considered in the main text and detailed in the **Methods** section. Statistical differences between distributions are assessed using the Mann–Whitney test; * indicates $p < 0.05$, ** indicates $p < 0.01$, and *** indicates $p < 0.001$.

1250 parameters, compared to 2 mechanistic parameters.

The analysis of CP distributions for trajectory reconstruction exhibits a broadly similar behavior in the Lotka–Volterra and Lorenz test cases. In the Damped Oscillator test case, no statistically significant differences are observed among the different scenarios, consistent with the results obtained for vector field reconstruction (it is worth noting that, in this test case, CP values remain close to the nominal threshold of 0.95 across all scenarios, which may further limit the detectability of differences between scenarios).

### S16 Comparing MOD with the Laplace Approximation (full reconstruction - numerical test cases)

In this section, we compare in the full reconstruction scenario of the numerical test cases the performance of MOD ensemble with another standard UQ method beyond the standard ensembles: the Laplace approximation [DKI+21, KHH20, IKB21]. This is a state-of-the-art method for UQ in machine learning, and it has been proposed also for NODEs [OTH23].

We briefly summarize the Laplace approximation for NODEs (we refer to [OTH23, DKI+21] for a complete description). Assuming a Gaussian prior $p(\theta) = \mathcal{N}(0, \sigma_0^2 I)$ on the neural network parameters $\theta$, given a training dataset $\mathcal{D}$, in the Bayesian framework the posterior distribution of the parameters is

$$p(\theta \mid \mathcal{D}) = \frac{p(\mathcal{D} \mid \theta)\, p(\theta)}{Z}, \qquad Z = \int p(\mathcal{D} \mid \theta)\, p(\theta)\, d\theta, \tag{8}$$

where $Z$ is the marginal likelihood. Since $Z$ is intractable for neural networks, the posterior is approximated locally around the trained parameters, corresponding in the Bayesian setting to the maximum a posteriori estimate $\theta_{\text{MAP}}$. A second-order Taylor expansion of the negative log-posterior at $\theta_{\text{MAP}}$ yields the Gaussian approximation

$$q(\theta) = \mathcal{N}(\theta_{\text{MAP}}, \Sigma), \qquad \Sigma = \left[-\nabla_\theta^2 \log p(\theta \mid \mathcal{D})\big|_{\theta_{\text{MAP}}}\right]^{-1} = \left[\nabla_\theta^2 \mathcal{L}(\theta)\big|_{\theta_{\text{MAP}}} + \sigma_0^{-2} I\right]^{-1}, \tag{9}$$

where $\mathcal{L}(\theta) = -\log p(\mathcal{D} \mid \theta)$. Uncertainty on the vector field and on novel trajectories can be obtained by sampling $n$ parameterizations from this distribution and proceeding as indicated in the **Methods** section of the main text.

To compare the UQ provided by MOD ensembles and the Laplace approximation, we considered the 10 trained MOD ensembles for each test-case. For each ensemble, we applied the Laplace approximation around the trained NODE corresponding to the initialization point of the MOD algorithm and drew 100 samples from the resulting Gaussian posterior over parameters. This allowed us to compare the coverage proportions obtained with MOD ensembles and with the Laplace approximation, both on the vector field and on test trajectories. The Hessian of the log-likelihood was approximated using the Gauss–Newton matrix [DKI+21]. Since no Tikhonov regularization (i.e., Gaussian prior) was used during training, the prior variance $\sigma_0^2$ was treated as a hyperparameter and selected by maximizing the approximate marginal likelihood [OTH23] (to ensure the validity of the local quadratic approximation underlying the Laplace method and the numerical stability of the ODE integration, the search for $\sigma_0^2$ was restricted to a range that keeps posterior mass within a neighborhood of the MAP estimate where the dynamics remain numerically stable).

The coverage proportions of the 0.95 prediction intervals on the vector field are reported in Table S10. In all test cases, the MOD ensembles achieve coverage proportions that are statistically significantly closer to the nominal level of 0.95 than those obtained with the Laplace approximation. The pointwise mean coverage analysis (Fig. S45) further shows that, under our experimental conditions, the Laplace-based prediction intervals tend to be overconfident, particularly in OOD regions far from the training data. This behavior is especially pronounced in the Damped Oscillator and Lorenz test cases, and is likely related to the mismatch between the true loss landscape and the local quadratic approximation underlying the Laplace method.

| | **Lotka–Volterra** | **Damped Oscillator** | **Lorenz** |
|---|---|---|---|
| Laplace approximation | $0.616 \pm 0.013$ | $0.203 \pm 0.009$ | $0.003 \pm 0.000$ |
| MOD | $\mathbf{0.986 \pm 0.006}$ (**) | $\mathbf{0.993 \pm 0.003}$ (**) | $\mathbf{0.760 \pm 0.040}$ (**) |

Table S10: **Comparison of the CP of 0.95-prediction intervals on the vector field (full reconstruction scenario).** Mean CP of 0.95-prediction intervals on the vector field within a selected region of the state space. The results are presented as the mean ± SEM across 10 ensembles. Statistical significance was assessed using the Wilcoxon signed-rank test on absolute deviations from 0.95 between the two distributions of CP values. Asterisks indicate the following significance levels: * $p \leq 0.05$, ** $p \leq 0.01$, and *** $p \leq 0.001$. The values shown in bold are those closest to the theoretical value.

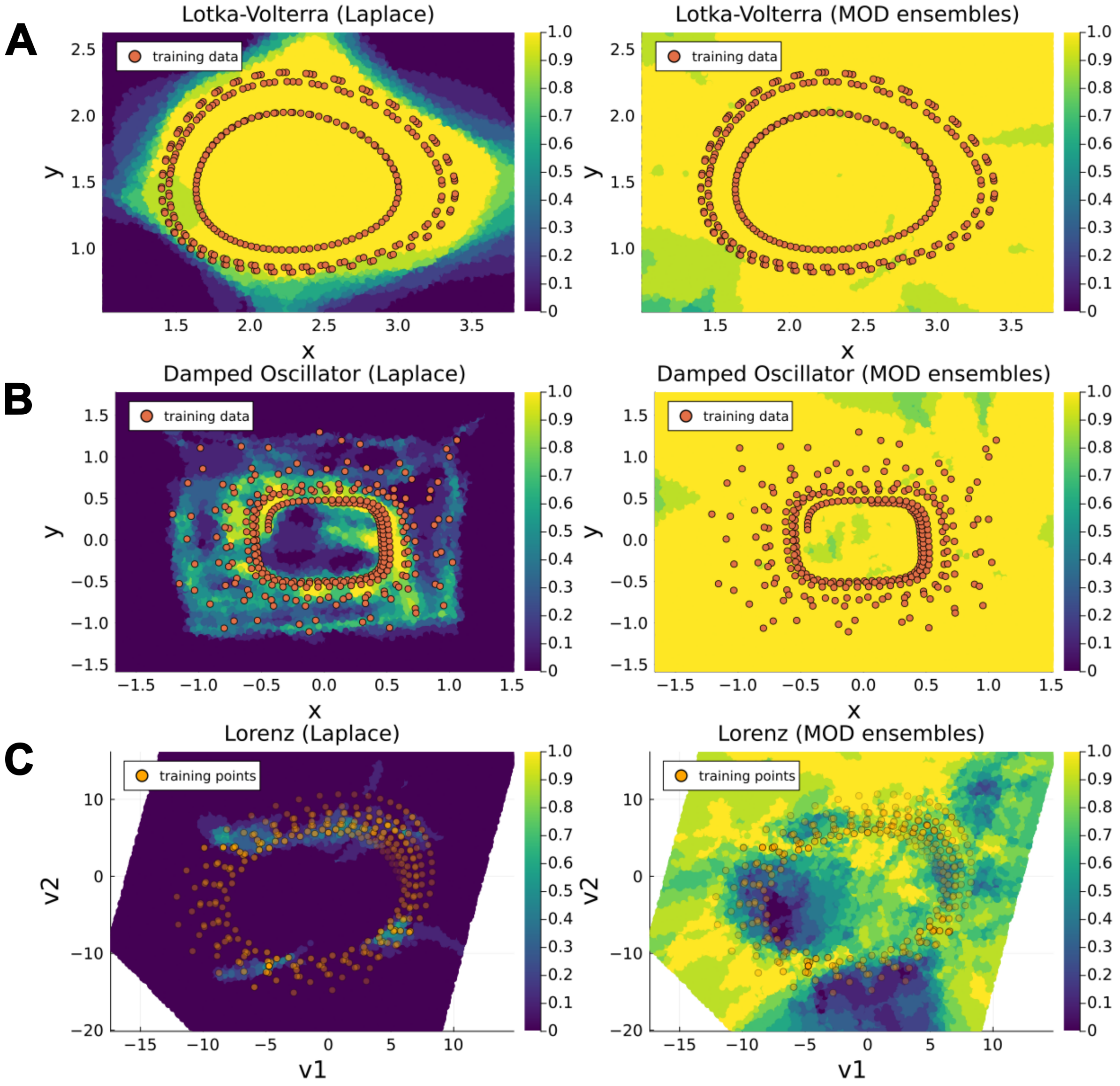


Figure S45: **Comparison of 0.95-prediction interval coverage on state space: Laplace approximation vs. MOD Ensembles (full reconstruction scenario).** Panels A, B, and C display heatmaps of the mean coverage (computed across 10 ensembles) of 0.95-prediction intervals on the state space for the Lotka–Volterra, Damped Oscillator, and Lorenz systems, respectively. For each system, the left subpanel shows results obtained with Laplace approximation, while the right subpanel shows results obtained with MOD ensembles. Orange overlays represent the points from the training trajectories, providing spatial context. For the Lorenz system, the mean coverage is visualized on a two-dimensional plane spanned by the coordinates $(v_1, v_2)$, obtained via linear regression on the Lorenz attractor (see Supplementary Section S8 for details). These coordinates provide a low-dimensional representation of the original three-dimensional state space $(x, y, z)$; here, the transparency of training trajectory points indicates their orthogonal distance from the training trajectory points to the plane.

The coverage proportions of the 0.95 prediction intervals on the test trajectories (reported in Table S11) are consistent with those observed for the vector field. In all test cases, the prediction intervals obtained with the MOD ensembles achieve coverage proportions that are statistically significantly closer to the nominal level of 0.95 than those obtained with the Laplace approximation. In this case as well, the difference is especially pronounced in the Lorenz test case.

| | **Lotka–Volterra** | **Damped Oscillator** | **Lorenz** |
|---|---|---|---|
| Laplace approximation | 0.483 ± 0.017 | 0.728 ± 0.019 | 0.019 ± 0.002 |
| MOD | **0.926 ± 0.019** (**) | **0.954 ± 0.012** (**) | **0.738 ± 0.030** (**) |

Table S11: **Comparison of the CP of 0.95-prediction intervals on the trajectories (full reconstruction scenario).** Mean CP of 0.95-prediction intervals on trajectories on the selected points of test trajectories. The results are presented as the mean ± SEM across 10 ensembles. Statistical significance was assessed using the Wilcoxon signed-rank test on absolute deviations from 0.95 between the two distributions of CP values. Asterisks indicate the following significance levels: * $p \leq 0.05$, ** $p \leq 0.01$, and *** $p \leq 0.001$. The values shown in bold are those closest to the theoretical value.

These results are particularly noteworthy when considering the loss achieved by the NODEs composing the MOD ensembles compared to the loss of the models sampled from the Laplace approximation. The NODEs drawn from the Laplace posterior are not constrained to satisfy the training accuracy threshold $\epsilon_{\text{acc}}$, as illustrated in the corresponding plot. In principle, this lack of constraint could increase the variability induced by the Laplace approximation, since it allows the inclusion of NODE parameterizations that perform worse on the training set (in contrast to the MOD ensembles). Nevertheless, as shown in the previous analysis, the resulting prediction intervals remain overconfident compared to those obtained with the MOD ensembles, underscoring the potential advantages of exploring the parameter space beyond a strictly local neighborhood of the MAP estimate.

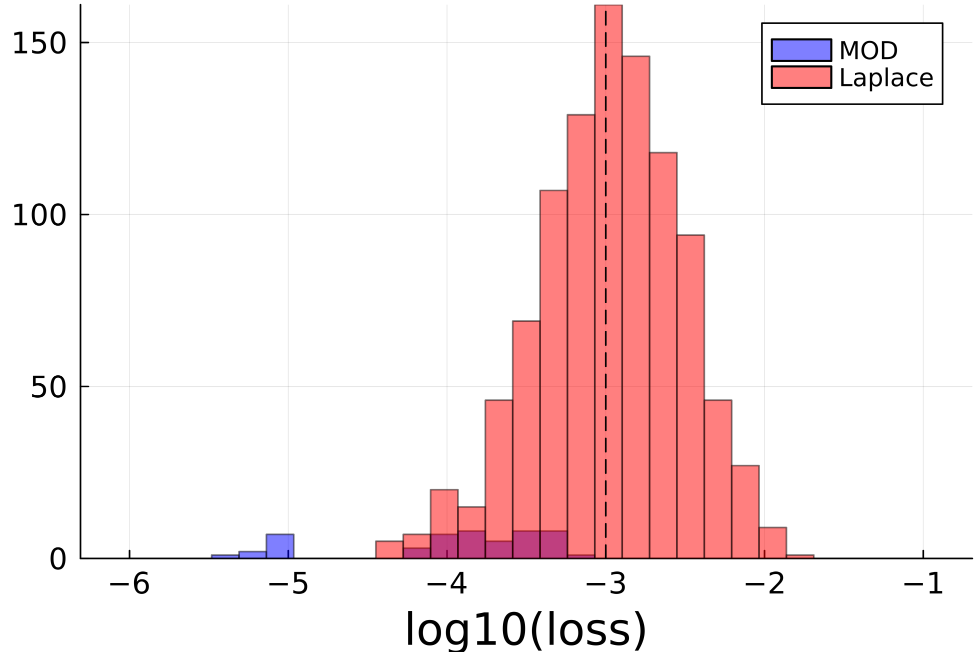


Figure S46: **Comparison among loss function values on training trajectories between models composing the ensembles sampled from the Laplace approximation and the MOD ensembles in the Lotka–Volterra test case.** The histogram shows the distribution of the training-set loss values achieved by the NODEs composing the ensembles sampled from the Laplace approximation (red, 1000 models) and the MOD ensembles (blue, 50 models). The dashed line indicates the training accuracy threshold $\epsilon_{\text{acc}}$ used during the MOD ensemble training.

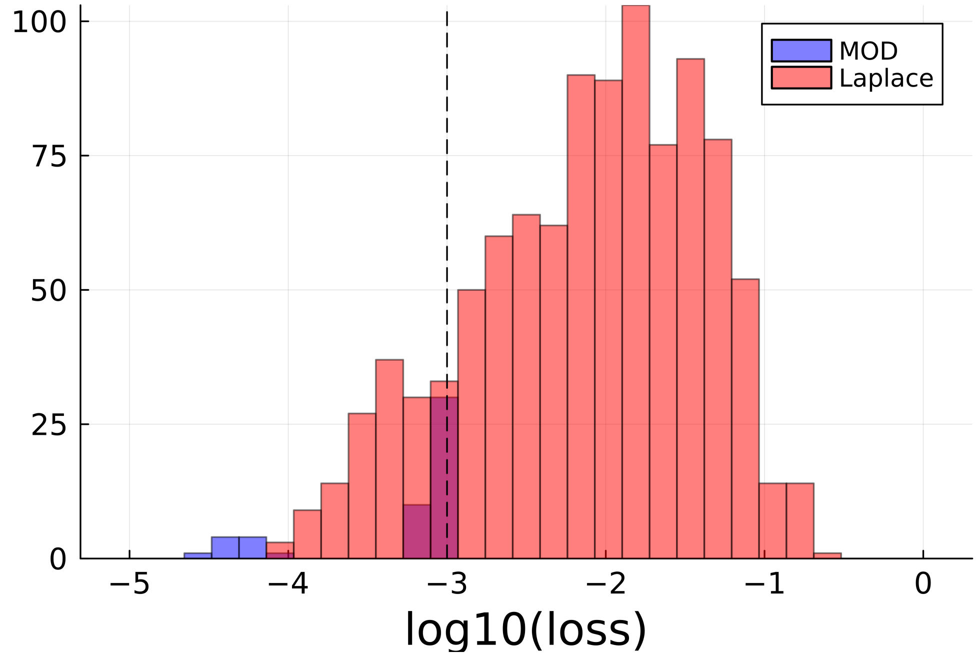


Figure S47: **Comparison among loss function values on training trajectories between models composing the ensembles sampled from the Laplace approximation and the MOD ensembles in the Damped Oscillator test case.** The histogram shows the distribution of the training-set loss values achieved by the NODEs composing the ensembles sampled from the Laplace approximation (red, 1000 models) and the MOD ensembles (blue, 50 models). The dashed line indicates the training accuracy threshold $\epsilon_{\mathrm{acc}}$ used during the MOD ensemble training.

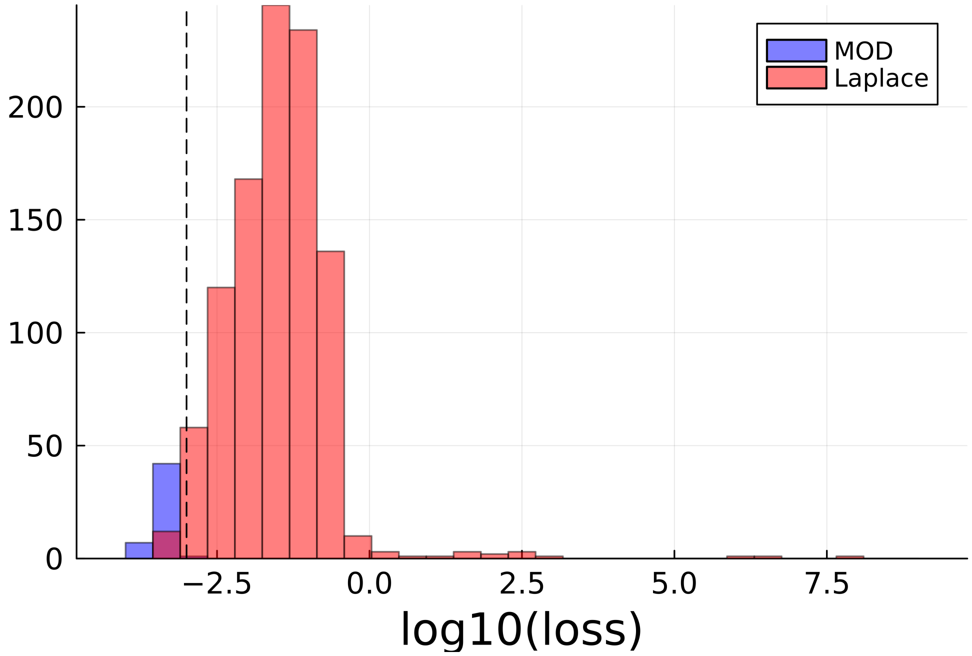


Figure S48: **Comparison among loss function values on training trajectories between models composing the ensembles sampled from the Laplace approximation and the MOD ensembles in the Lorenz test case.** The histogram shows the distribution of the training-set loss values achieved by the NODEs composing the ensembles sampled from the Laplace approximation (red, 1000 models) and the MOD ensembles (blue, 50 models). The dashed line indicates the training accuracy threshold $\epsilon_{\text{acc}}$ used during the MOD ensemble training.

### S17 Numerical test cases: full reconstruction with lower $\epsilon_{\mathbf{acc}}$

This section presents the results of the proposed algorithm for training MOD ensembles using a loss threshold of $\epsilon_{\text{acc}} = 10^{-4}$. The algorithm was applied only to the Lotka–Volterra and Damped Oscillator test cases, as the models in standard ensembles for the Lorenz test case never achieved a loss value below the specified threshold.

Following the methodology outlined in the main text, we first trained 10 standard ensembles using the defined training trajectories. From these, we retained only the models that achieved a loss below $10^{-4}$. For each ensemble, we then randomly selected one of the accepted models to initialize our MOD ensemble training algorithm. This process resulted in 10 MOD ensembles, each composed of models with loss values below the fixed threshold.

The fit results of the models composing the standard ensembles on the training trajectories are shown in Figs. S49 and S50, corresponding to the Lotka–Volterra and Damped Oscillator test cases, respectively. Similarly, the fit results of the models composing the MOD ensembles on the same training trajectories are presented in Figs. S51 and S52, again for the Lotka–Volterra and Damped Oscillator cases, respectively. In both test cases, the fit results of the standard and MOD ensembles are visually indistinguishable.

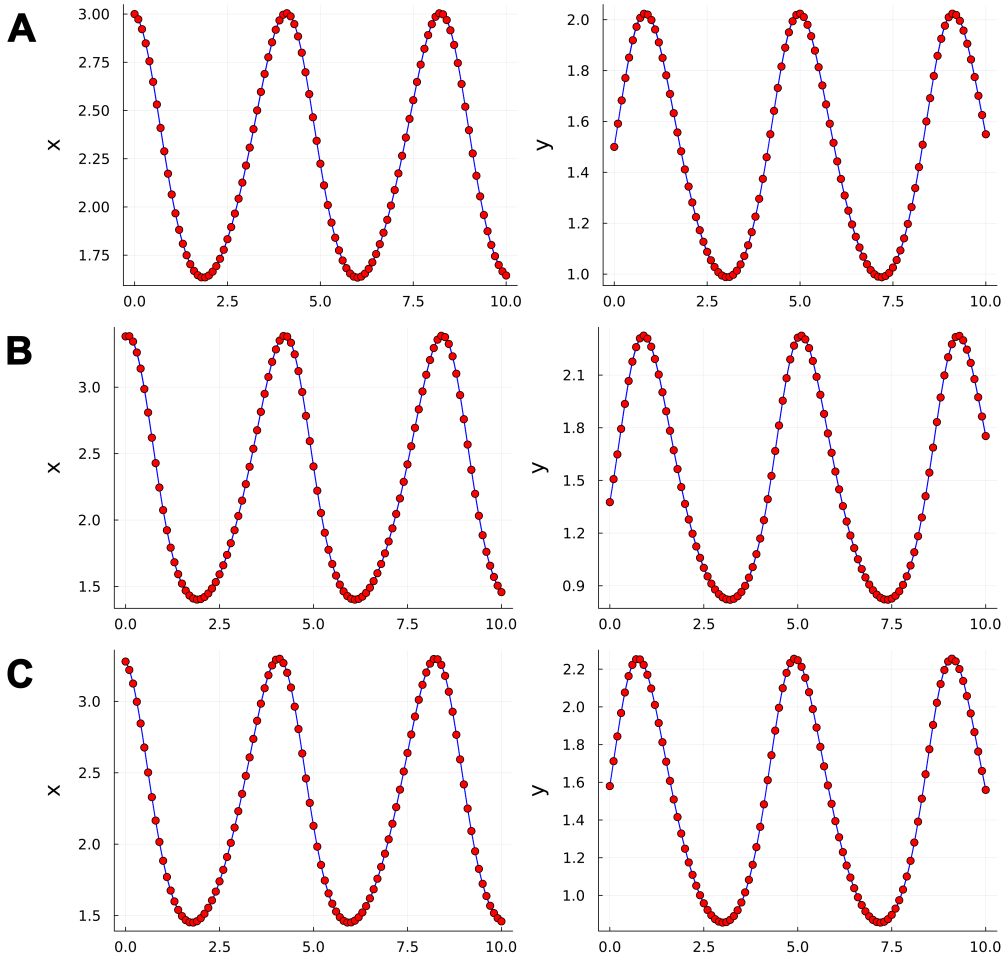


Figure S49: **Fit of trained data-driven models composing the standard ensembles on training trajectories for the Lotka–Voterra test case.** Panels A, B, and C show the fits of the trained models (blue lines) to the data points of the first, second, and third training trajectories (red points), respectively. A total of 50 models were trained (5 per ensemble, across 10 ensembles).

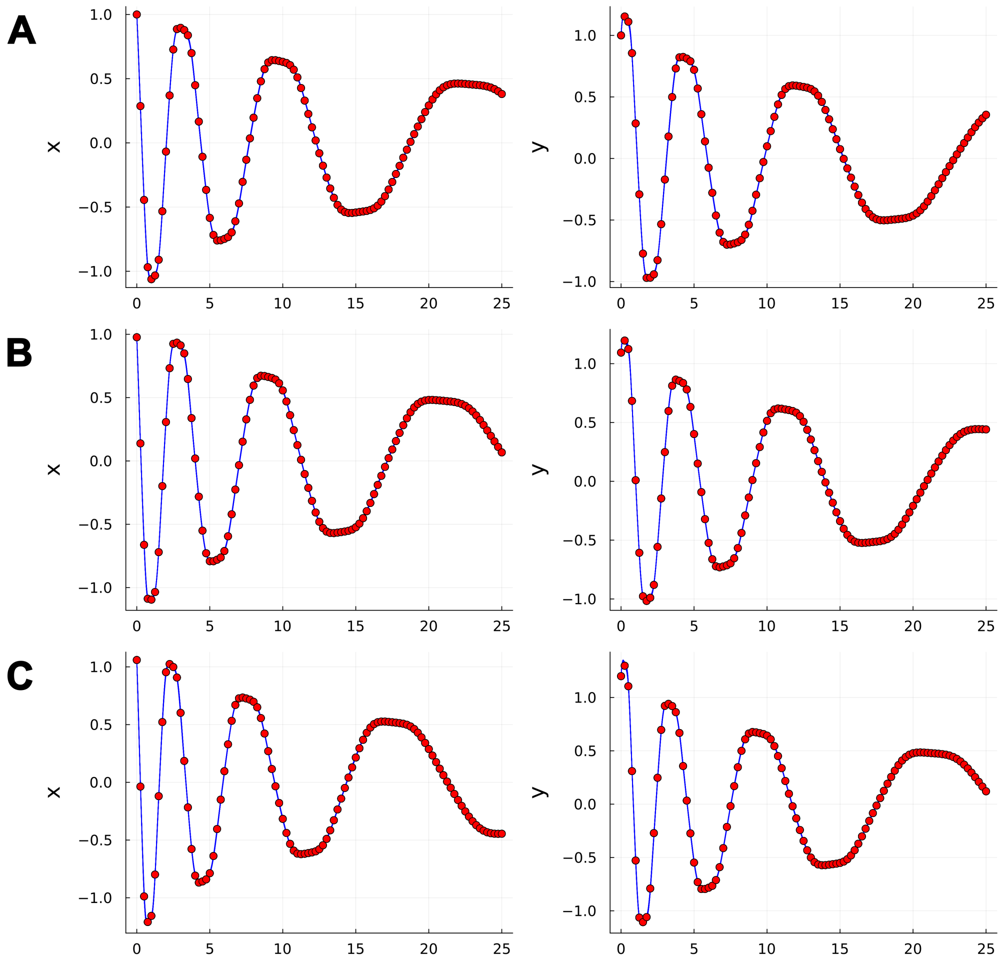


Figure S50: **Fit of trained data-driven models composing the standard ensembles on training trajectories for the Damped Oscillator test case.** Panels A, B, and C show the fits of the trained models (blue lines) to the data points of the first, second, and third training trajectories (red points), respectively. A total of 50 models were trained (5 per ensemble, across 10 ensembles).

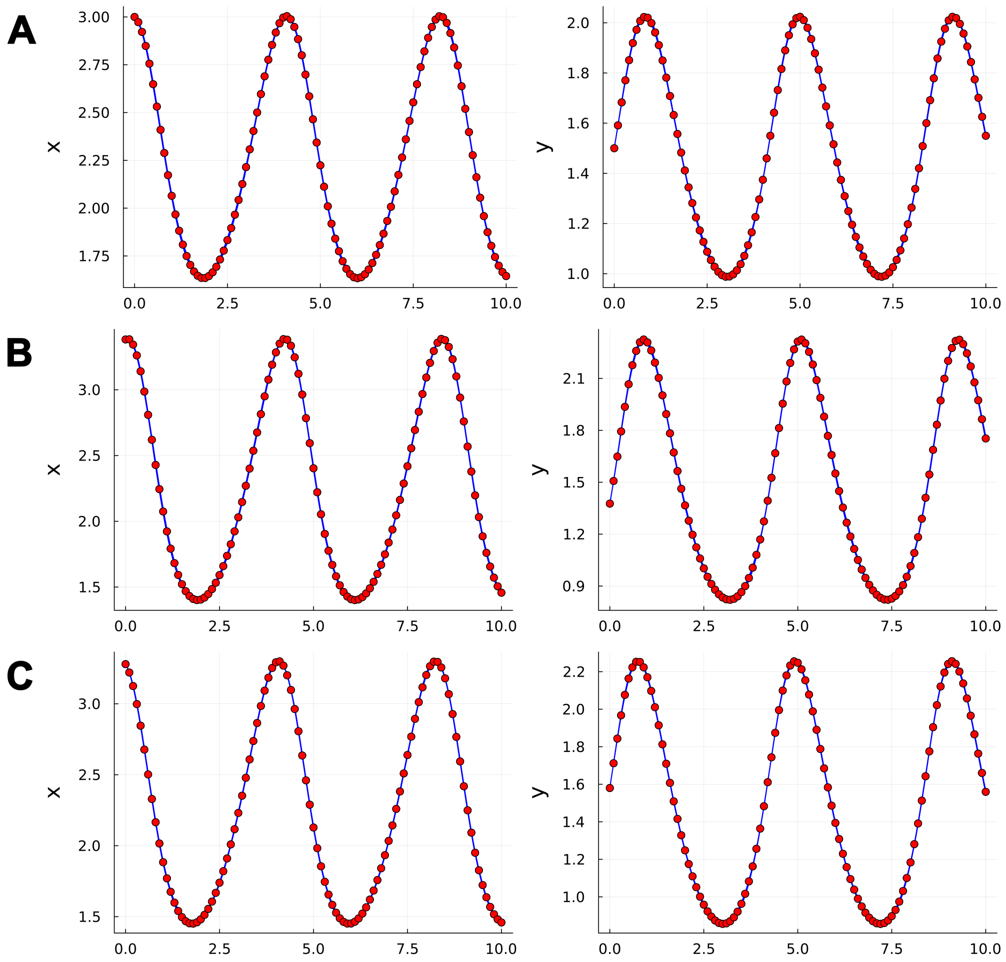


Figure S51: **Fit of trained data-driven models composing the MOD ensembles on training trajectories for the Lotka–Voterra test case.** Panels A, B, and C show the fits of the trained models (blue lines) to the data points of the first, second, and third training trajectories (red points), respectively. A total of 50 models were trained (5 per ensemble, across 10 ensembles).

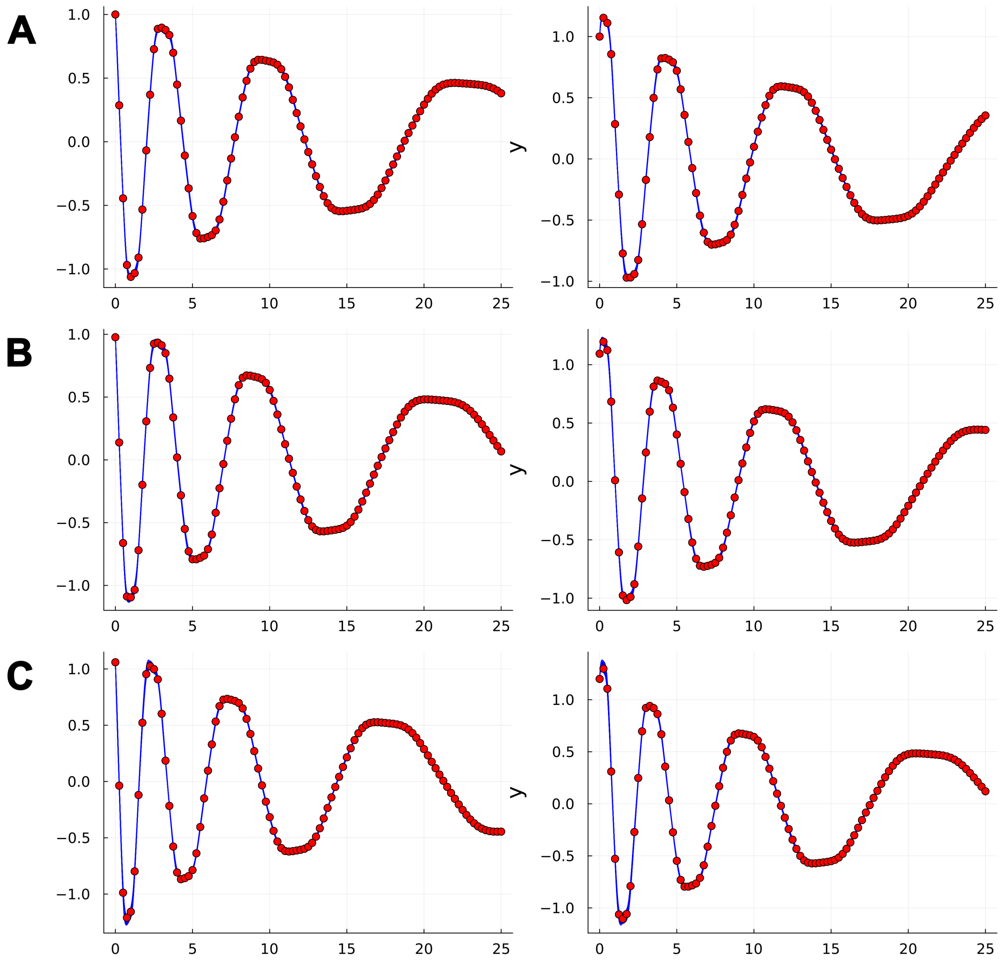


Figure S52: **Fit of trained data-driven models composing the MOD ensembles on training trajectories for the Damped Oscillator test case.** Panels A, B, and C show the fits of the trained models (blue lines) to the data points of the first, second, and third training trajectories (red points), respectively. A total of 50 models were trained (5 per ensemble, across 10 ensembles).

We begin by analyzing the coverage of 0.95-prediction intervals on the vector field obtained with the two approaches (standard and MOD ensembles). The CP results are reported in Table S12, whereas the mean coverage heatmaps in the considered regions of the state space are shown in Fig. S53. The results with the threshold $\epsilon_{\text{acc}} = 10^{-4}$ are compatible to the results obtained with $\epsilon_{\text{acc}} = 10^{-3}$ presented in the main text of the paper.

| | **Lotka–Volterra** | **Damped Oscillator** |
|---|---|---|
| Standard | 0.850 ± 0.024 | 0.664 ± 0.015 |
| MOD | **0.994 ± 0.002** (**) | **0.965 ± 0.007** (**) |

Table S12: **Comparison of the CP of 0.95-prediction intervals on the vector field.** Mean CP of 0.95-prediction intervals on the vector field within a selected region of the state space. The results are presented as the mean ± SEM across 10 ensembles. Statistical significance was assessed using the Wilcoxon signed-rank test on absolute deviations from 0.95 between the two distributions of CP values. Asterisks indicate the following significance levels: * p $\leq$ 0.05, ** p $\leq$ 0.01, and *** p $\leq$ 0.001. The values shown in bold are those closest to the theoretical value.

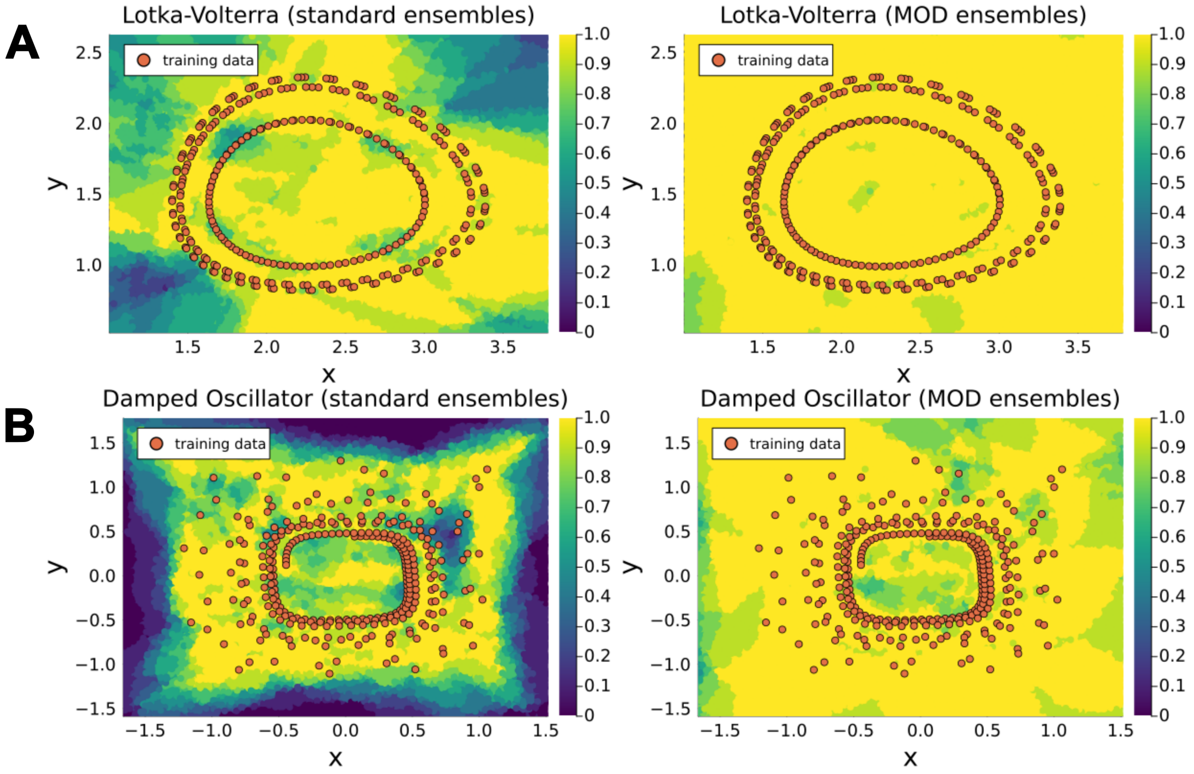


Figure S53: **Comparison of 0.95-prediction interval coverage on state space: Standard vs. MOD Ensembles.** Panels A, and B display heatmaps of the mean coverage (computed across 10 ensembles) of 0.95-prediction intervals on the state space for the Lotka–Volterra, and Damped Oscillator. For each system, the left subpanel shows results obtained with standard ensembles, while the right subpanel shows results obtained with MOD ensembles. Orange overlays represent the points from the training trajectories, providing spatial context.

The results in terms of coverage of the 0.95-prediction intervals on the trajectories are summarized in Table S13. Also in this case, the results obtained using the threshold $\epsilon_{\text{acc}} = 10^{-4}$ are consistent with those obtained using $\epsilon_{\text{acc}} = 10^{-3}$, as presented in the main text of the paper.

| | **Lotka–Volterra** | **Damped Oscillator** |
|---|---|---|
| Standard | 0.800 ± 0.042 | 0.599 ± 0.019 |
| MOD | **0.940 ± 0.016** (**) | **0.834 ± 0.033** (**) |

Table S13: **Comparison of the CP of** 0.95**-prediction intervals on the trajectories.** Mean CP of 0.95-prediction intervals on trajectories on the selected points of test trajectories. The results are presented as the mean ± SEM across 10 ensembles. Statistical significance was assessed using the Wilcoxon signed-rank test on absolute deviations from 0.95 between the two distributions of CP values. Asterisks indicate the following significance levels: * $p \leq 0.05$, ** $p \leq 0.01$, and *** $p \leq 0.001$. The values shown in bold are those closest to the theoretical value.

### S18 Supplementary Discussion: threshold for accepting an ensemble member.

The threshold $\epsilon_{\text{acc}}$ on the cost function used both to select the members to accept in the standard ensemble and to run the algorithm for training MOD ensembles is let as an arbitrary choice of the modeler. We have shown that the MOD ensemble training algorithm can function under different threshold values (Supplementary Section S17), but there is no formal justification for preferring one over another. Our cost function could be straightforwardly adapted to a likelihood-based formulation by assuming that the error model governing the phenomenon is an i.i.d. Gaussian process with a standard deviation proportional to the min-max oscillations of the variables. In mechanistic modeling theory, under this assumption, the threshold on the cost value can be derived from the likelihood ratio test [VRHB22]. We did not adopt this statistical criterion due to the non-identifiability of the models under investigation. It is well established that the asymptotic theory underpinning the likelihood ratio test does not apply in cases of non-identifiability [Lin95], which is the case for NODEs and UDEs, at least concerning the NN parameters. While extensions have been proposed to characterize the asymptotic distribution of the likelihood ratio test under specific forms of non-identifiability [LS03], these have not been considered in this work, and the assessment of a statistically meaningful threshold remains beyond the scope of the present study.

#### S19 Analysis of coverage on extended regions of state space (full reconstruction - numerical test cases)

In this section, we analyze the coverage proportion of 0.95 prediction intervals for the vector field obtained using both standard and MOD ensembles, evaluated over an extended region of the state space, in the full reconstruction scenario of the numerical test cases. The extended region is defined by expanding the original bounding box—used in the main text—by 100% along each dimension. The results for the three test cases are presented in Fig. S54.

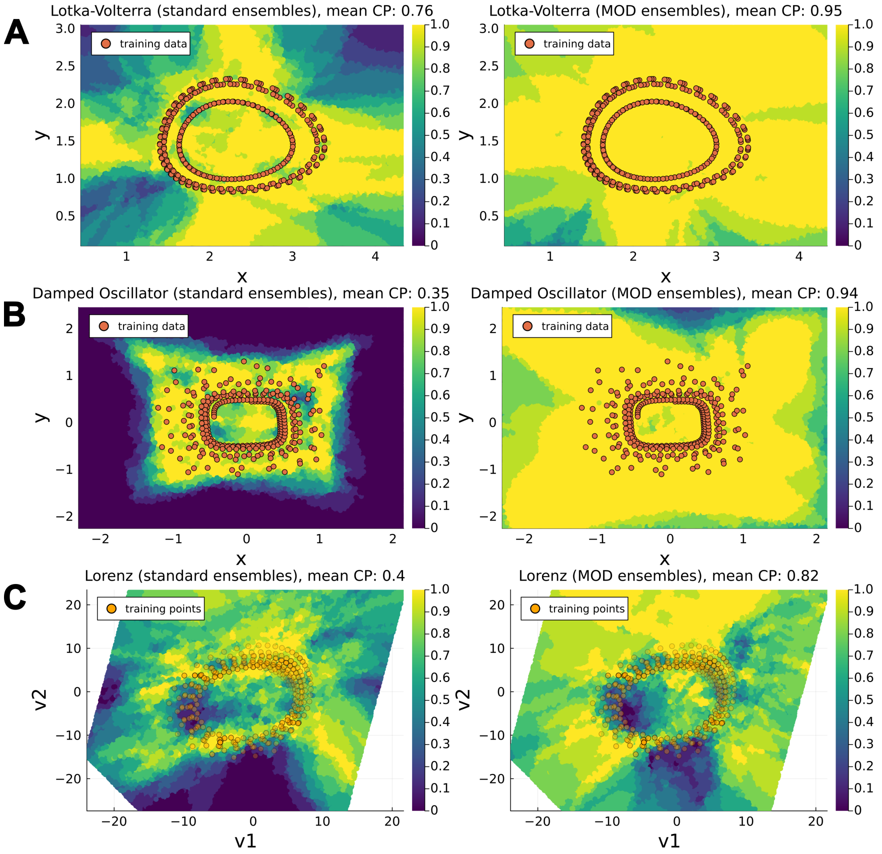


Figure S54: **Comparison of 0.95-prediction interval coverage on vector field in extended region of state space: Standard vs. MOD Ensembles.** Panels A, B, and C display heatmaps of the mean coverage (computed across 10 ensembles) of 0.95-prediction intervals on the state space for the Lotka–Volterra, Damped Oscillator, and Lorenz systems, respectively. For each system, the left subpanel shows results obtained with standard ensembles, while the right subpanel shows results obtained with MOD ensembles. Orange overlays represent the points from the training trajectories, providing spatial context. For the Lorenz system, the mean coverage is visualized on a two-dimensional plane spanned by the coordinates $(v_1, v_2)$, obtained via linear regression on the Lorenz attractor (see Supplementary Section S8 for details). These coordinates provide a low-dimensional representation of the original three-dimensional state space $(x, y, z)$ ; here, the transparency of training trajectory points indicates their orthogonal distance from the training trajectory points to the plane. The mean coverage proportion in the region is reported in the title for each plot.

### S20 Analysis of extended regions of state space (partial reconstruction with unknown mechanistic parameters - numerical test cases)

In this section, we analyze the coverage proportion of 0.95 prediction intervals for the vector field obtained using both standard and MOD ensembles, evaluated over an extended region of the state space, in the partial reconstruction scenario of the numerical test cases. The extended region is defined by expanding the original bounding box—used in the main text—by 100% along each dimension. The results for the three test cases are presented in Fig. S55.

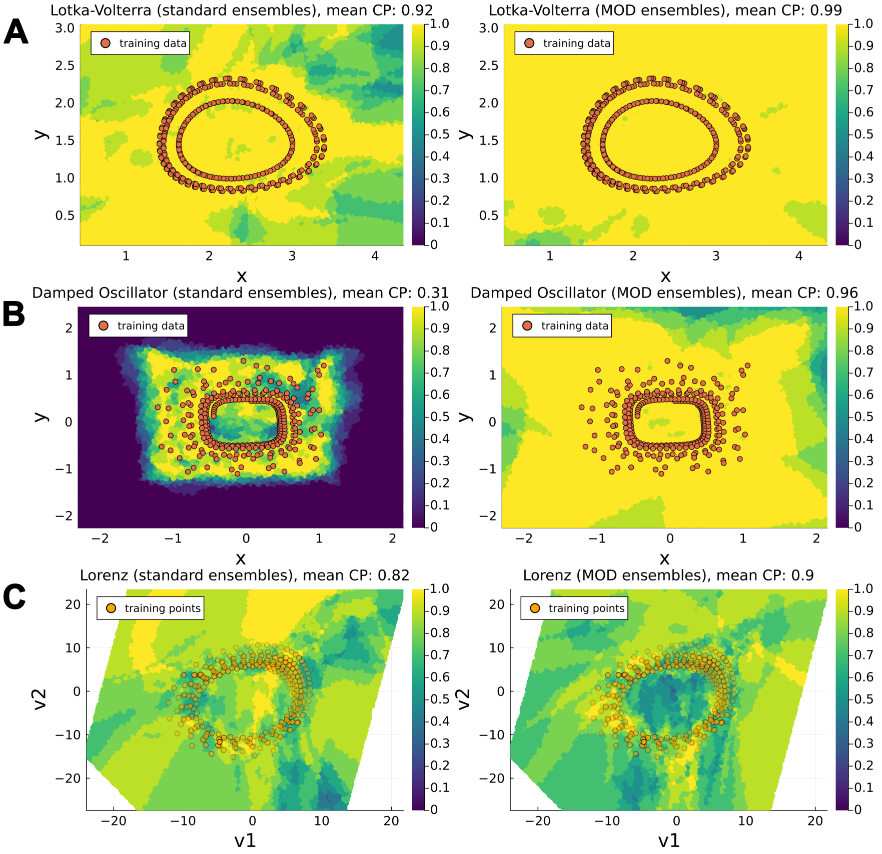


Figure S55: **Comparison of 0.95-prediction interval coverage on vector field in extended region of state space: Standard vs. MOD Ensembles.** Panels A, B, and C display heatmaps of the mean coverage (computed across 10 ensembles) of 0.95-prediction intervals on the state space for the Lotka–Volterra, Damped Oscillator, and Lorenz systems, respectively. For each system, the left subpanel shows results obtained with standard ensembles, while the right subpanel shows results obtained with MOD ensembles. Orange overlays represent the points from the training trajectories, providing spatial context. For the Lorenz system, the mean coverage is visualized on a two-dimensional plane spanned by the coordinates $(v_1, v_2)$, obtained via linear regression on the Lorenz attractor (see Supplementary Section S8 for details). These coordinates provide a low-dimensional representation of the original three-dimensional state space $(x, y, z)$ ; here, the transparency of training trajectory points indicates their orthogonal distance from the training trajectory points to the plane. The mean coverage proportion in the region is reported in the title for each plot.

### S21 Analysis of objective function behavior during training

In this section, we analyze the behavior of the objective function (the OOD disagreement) during the MOD ensemble training. This analysis motivates our decision to train the models for a fixed number of epochs, rather than imposing a threshold on the objective function value or using a convergence condition to stop the training. Since the objective function behaviors are similar across the various models trained in a MOD ensemble (4 models in the experiments described in the main text, beyond the starting parameterization) and across the different scenarios considered (purely data-driven, partially data-driven with fixed mechanistic parameters, and partially data-driven), we present the analysis only for the first model trained in the MOD ensemble under the partially data-driven scenario.

Fig. S21 shows the behavior of the objective function during the training of the first model in the MOD ensembles for the Lotka–Volterra, Damped Oscillator, and Lorenz systems. As expected, the objective function takes different values for each system, which led us to exclude the use of a fixed threshold to stop the optimization. Additionally, there is no evident convergence to a value within the 800 iterations considered. This is due to the flexibility of the neural network, which can assume very different values, especially far from the training domain, making the disagreement unlikely to converge to a limiting asymptotic value. This consideration led us to exclude the use of a convergence condition to stop the training.

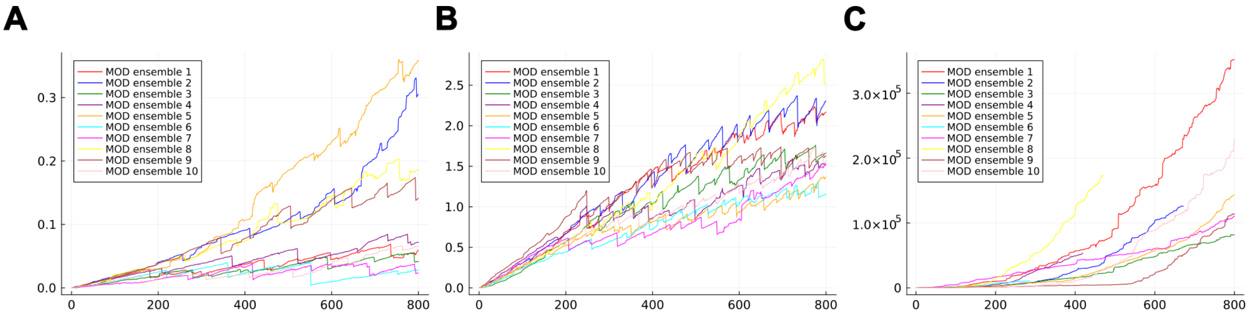


Figure S56: **Behavior of the objective function during the training of the MOD ensembles.** Panels A, B, and C display the behavior of the objective function during the training of the first model in the MOD ensembles (10 MOD ensembles were trained for each system) in the partially data-driven scenario for the Lotka–Volterra, Damped Oscillator, and Lorenz systems, respectively. The training was completed before 800 iterations, particularly for the Lorenz system, if it was not possible to reproject the parameters to a region where the loss function is less than $\epsilon_{\text{acc}}$.

Interestingly, the profile of the objective function displays a sawtooth pattern, particularly evident for the Lotka–Volterra and Damped Oscillator systems. This behavior is due to the reprojection mechanism, as explained in the **Methods** section of the main text: when the objective function exceeds a fixed threshold $\epsilon_{\text{acc}}$, the parameters are reprojected onto the minimum loss manifold via a short optimization. To demonstrate that this reprojection mechanism does not return the parameters to the starting point of the training algorithm, we analyzed the dynamics of the squared distance from the initial point during the training of the models in the MOD ensembles, as shown in Fig. S21. The monotonic increase of the distances indicates that the reprojection does not return the parameters toward the starting point of the training algorithm.

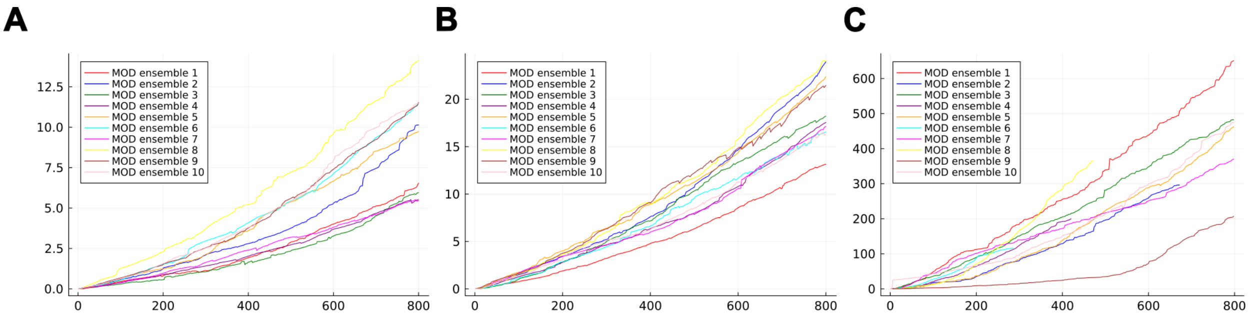


Figure S57: **Distance from the starting point during the training of the MOD ensembles.** Panels A, B, and C display the squared distance from the starting point of the parameters during the training of the first model in the MOD ensembles (10 MOD ensembles were trained for each system) in the partially data-driven scenario for the Lotka–Volterra, Damped Oscillator, and Lorenz systems, respectively. The training was completed before 800 iterations, particularly for the Lorenz system, if it was not possible to reproject the parameters to a region where the loss function was less than $\epsilon_{\text{acc}}$.

### S22 Investigation of undercoverage in the Lorenz test-case (full reconstruction setting)

In this section, we analyze the possible reasons for the undercoverage of the prediction intervals derived from the MOD ensembles in the Lorenz test case. We focus on the full reconstruction scenario, where the undercoverage is particularly pronounced.

In the results presented in the main text, the MOD ensemble training algorithm was run with a fixed maximum number of 800 epochs for each ensemble member. This choice was made to illustrate the behavior of the algorithm across different test cases and scenarios while keeping the same hyperparameters. However, since the Lorenz system is chaotic, in this section we allow for a more extensive exploration of the parameter space by increasing the maximum number of training epochs to 1500 per ensemble member. We compare the results obtained with a maximum of 800 and 1500 epochs in terms of coverage performance on both the vector field and previously unseen trajectories. All other hyperparameters are kept fixed to the values used in the full reconstruction scenario of the Lorenz test case presented in the main text.

The comparison of the mean coverage proportions of the 0.95 prediction intervals on the vector field (reported in Table S14) shows that increasing the maximum number of training epochs raises the mean coverage from 0.760 to 0.811. The Wilcoxon signed-rank test, applied to the absolute deviations from the nominal level of 0.95, yields a p-value of 0.051, which is slightly above the conventional significance threshold. This improvement is also qualitatively reflected in the mean coverage heatmap (Fig. S58). The results on previously unseen trajectories are consistent with those observed for the vector field: increasing the maximum number of epochs improves the mean coverage from 0.738 to 0.838. In this case, the distribution of coverage proportions is statistically significantly closer to the nominal value of 0.95.

| | **Vector field** | **Trajectories** |
|---|---|---|
| MOD (800 epochs) | $0.760 \pm 0.040$ | $0.738 \pm 0.030$ |
| MOD (1500 epochs) | $\mathbf{0.811 \pm 0.023}$ | $\mathbf{0.838 \pm 0.033}$ (**) |

Table S14: **Comparison of the CP of 0.95-prediction intervals on the vector field and trajectories (full reconstruction scenario).** Mean coverage proportions (CP) of the 0.95 prediction intervals on selected points of the test trajectories and on the vector field within a selected region of the state space. Results are reported as mean ± SEM across 10 ensembles. Statistical significance was assessed using the Wilcoxon signed-rank test on the absolute deviations from the nominal level of 0.95 between the two distributions of CP values. Asterisks denote significance levels as follows: * $p \leq 0.05$, ** $p \leq 0.01$, and *** $p \leq 0.001$. Bold values indicate those closest to the nominal level.

It is worth noting that, during the training of the MOD ensemble members, the maximum number of epochs is not necessarily reached. At each iteration, the displacement along the direction that maximizes disagreement while keeping the cost function locally constant is only an approximation; as a consequence, the cost function may increase. If it exceeds the prescribed accuracy threshold ($10^{-3}$ in our case), the algorithm attempts to re-project the current parameterization back into a low-cost region through a short optimization procedure (see the **Method** Section in the main text). If this reprojection step fails, the training of that ensemble member is terminated prematurely. Fig. S59 reports the number of epochs actually performed by the MOD algorithm for each trained model. In the Lorenz case, the vast majority of the members stop before reaching 1500 epochs, because the algorithm is unable to successfully reproject the solution into a low-cost region. The difficulty in re-optimization may stem from the fact that the MOD algorithm drives the parameterization into regions where numerical integration becomes more challenging. For this reason, training was not carried beyond 1500 epochs.

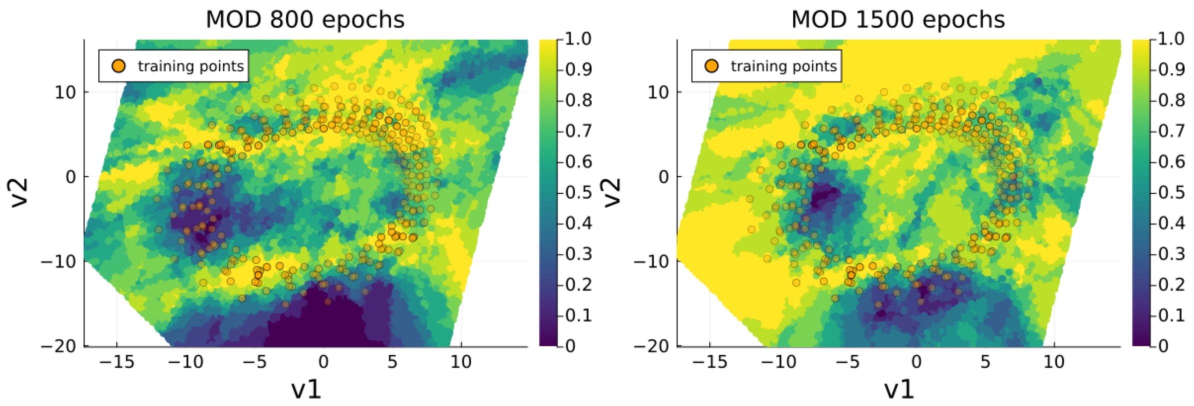


Figure S58: **Comparison of 0.95-prediction interval coverage on state space: MOD Ensembles with 800 epochs vs. 1500 epochs (full reconstruction scenario).** Heatmaps of the mean coverage (computed across 10 ensembles) of 0.95-prediction intervals on the state space for the Lorenz systems. The left subpanel shows results obtained with 800 epochs, while the right subpanel shows results obtained with 1500 epochs. Orange overlays represent the points from the training trajectories, providing spatial context. The mean coverage is visualized on a two-dimensional plane spanned by the coordinates $(v_1, v_2)$, obtained via linear regression on the Lorenz attractor (see Supplementary Section S8 for details). These coordinates provide a low-dimensional representation of the original three-dimensional state space $(x, y, z)$; here, the transparency of training trajectory points indicates their orthogonal distance from the training trajectory points to the plane.

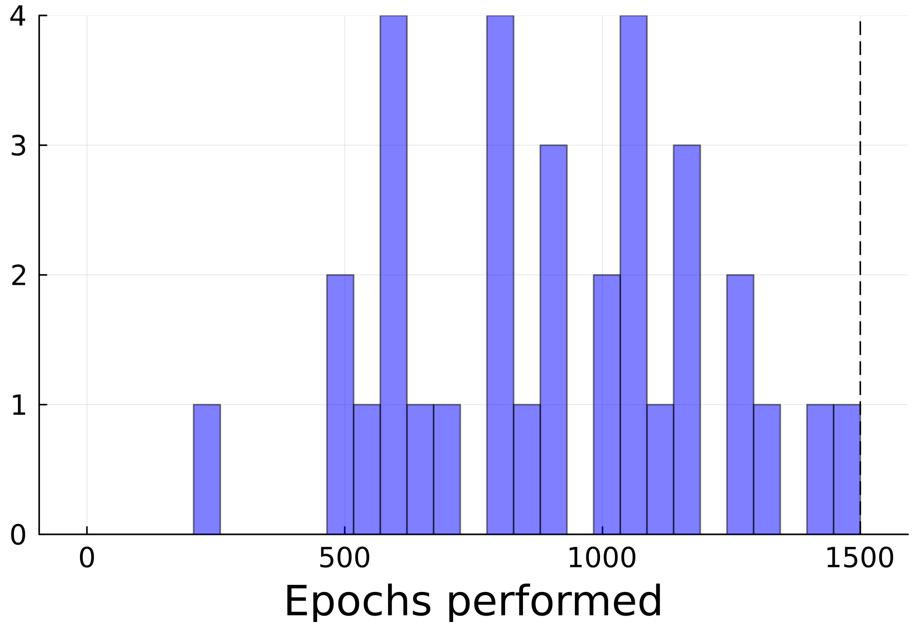


Figure S59: **Histogram of the number of training epochs performed by MOD ensemble members.** The histogram reports the frequency of the number of epochs performed by the MOD ensemble algorithm for each model (4 per ensemble, as the initial member is fixed, resulting in 40 models in total). The dashed line indicates the maximum number of epochs, set to 1500.

#### S23 Selection of the threshold $\epsilon_{eig}$

The hyperparameter $\epsilon_{eig}$ determines the threshold for identifying the null subspace of the Hessian matrix. Although its conceptual role is to select the "almost" zero eigenvalues, its value is not uniquely determined and must be chosen heuristically. In practice, the threshold mediates between two competing effects: values that are too large may include directions associated with increasing cost, requiring smaller adaptive step sizes to preserve stability, whereas values that are too small may prevent the exploration of nearly flat directions of the cost function, effectively limiting parameter perturbation. While in this work we adopt a conservative choice ($\epsilon_{eig} = 10^{-1}$) to favor the robust exploration of the parameter space, the performance of the method may still depend on this hyperparameter. Future work could investigate principled strategies for its automatic calibration.

### S24 Analysis of qualitative characteristics of the reconstructed vector fields

In this section, we assess whether the models comprising the MOD ensembles preserve key qualitative characteristics of the ground-truth vector fields. Specifically, we focus on the periodicity of the trajectories in the Lotka–Volterra system and the existence of a global attractor at (0.0) in the Damped Oscillator system. We show that these characteristics are preserved by the vector fields reconstructed by the MOD ensemble members only within a neighborhood of the training trajectories. However, when examining regions further out-of-distribution, these features are no longer maintained.

For analyzing the periodicity of the trajectories in the models reconstructing the vector field of the Lotka–Volterra system, we simulate the models comprising the MOD ensembles trained in each scenario —purely data-driven, partially data-driven with fixed mechanistic parameters, and partially data-driven— over the time interval $[0, 10]$, starting from initial points progressively farther from the training trajectories (note that the ground truth system always has a period of less than 10 for each initial point considered). The results, shown in Fig. S60, reveal that the periodicity (or at least a similar period to the ground truth system) is maintained for initial points closest to the training trajectories (blue trajectories in Fig. S60). However, when the initial points are farther from the training trajectories, the periodicity is not preserved, as seen with the pink and green trajectories in Fig. S60.

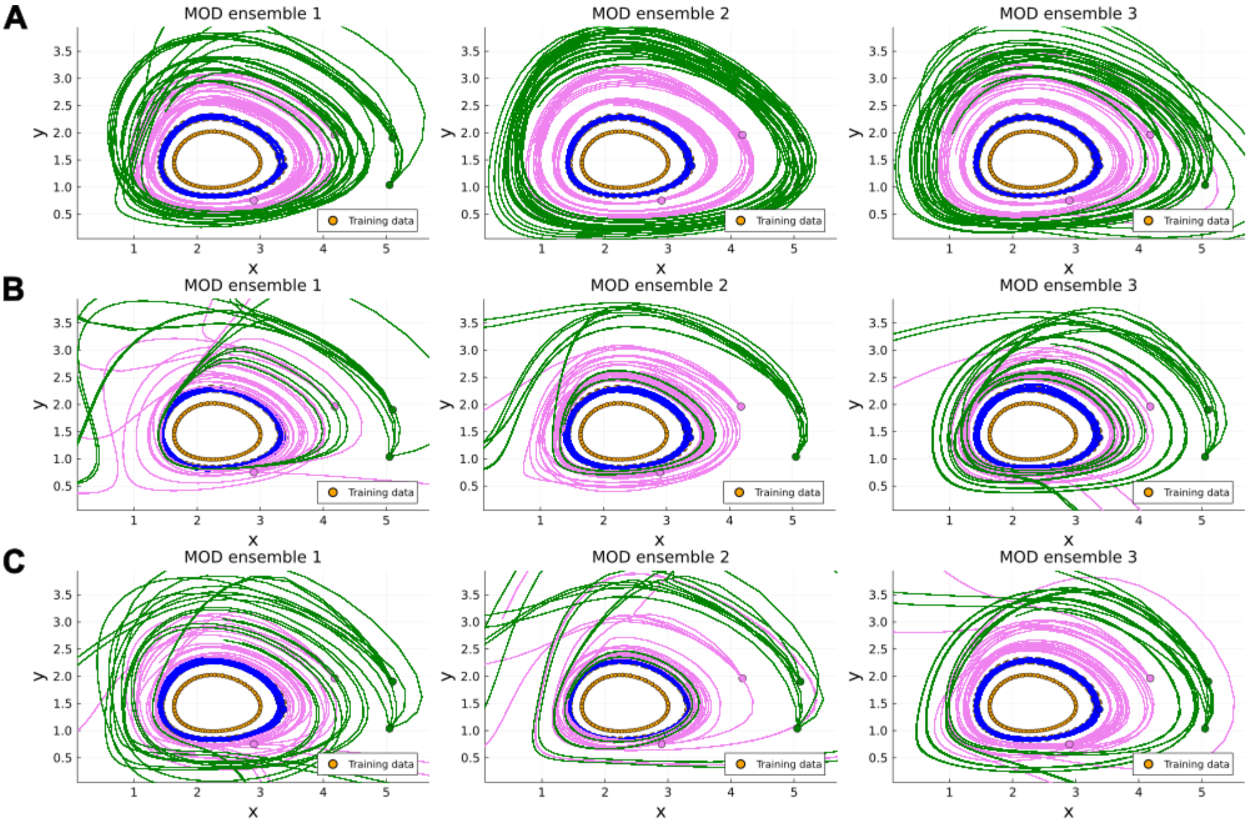


Figure S60: **Simulation of trajectories using models from the MOD ensembles starting from different initial points.** Panels A, B, and C display the trajectories obtained by simulating the models of the first three trained MOD ensembles in the purely data-driven scenario, the partially data-driven with fixed mechanistic parameters scenario, and the partially data-driven scenario, respectively, over the time interval $[0, 10]$. The dots represent the initial points considered. The color indicates the distance from the training sets: blue for trajectories starting near the training trajectories, pink for trajectories with an increased distance from the training trajectories, and green for trajectories starting in initial points with a further increased distance. Orange points (often covered by the blue lines) represent the training trajectories.

For analyzing the existence of a global attractor at $(0, 0)$ in the models composing the MOD ensembles for the Damped Oscillator system, we select initial points from different regions of the state space at varying distances from the training trajectories (see Fig. S24). We then simulate the models over the interval $[0, 100]$ and analyze the distribution of the norm of the last state of the system, comparing it with the norm of the last state of the ground-truth system. If the reconstructed vector fields preserve the existence of the attractor at $(0, 0)$ with a similar velocity to the ground truth system, the distribution should be concentrated around 0.0, as it is in the ground truth system. The results for the MOD ensembles trained in each scenario—purely data-driven, partially data-driven with fixed mechanistic parameters, and partially data-driven—indicate that, especially when the initial points are selected from regions farthest from the training trajectories, a portion of the trajectories does not converge to $(0, 0)$.

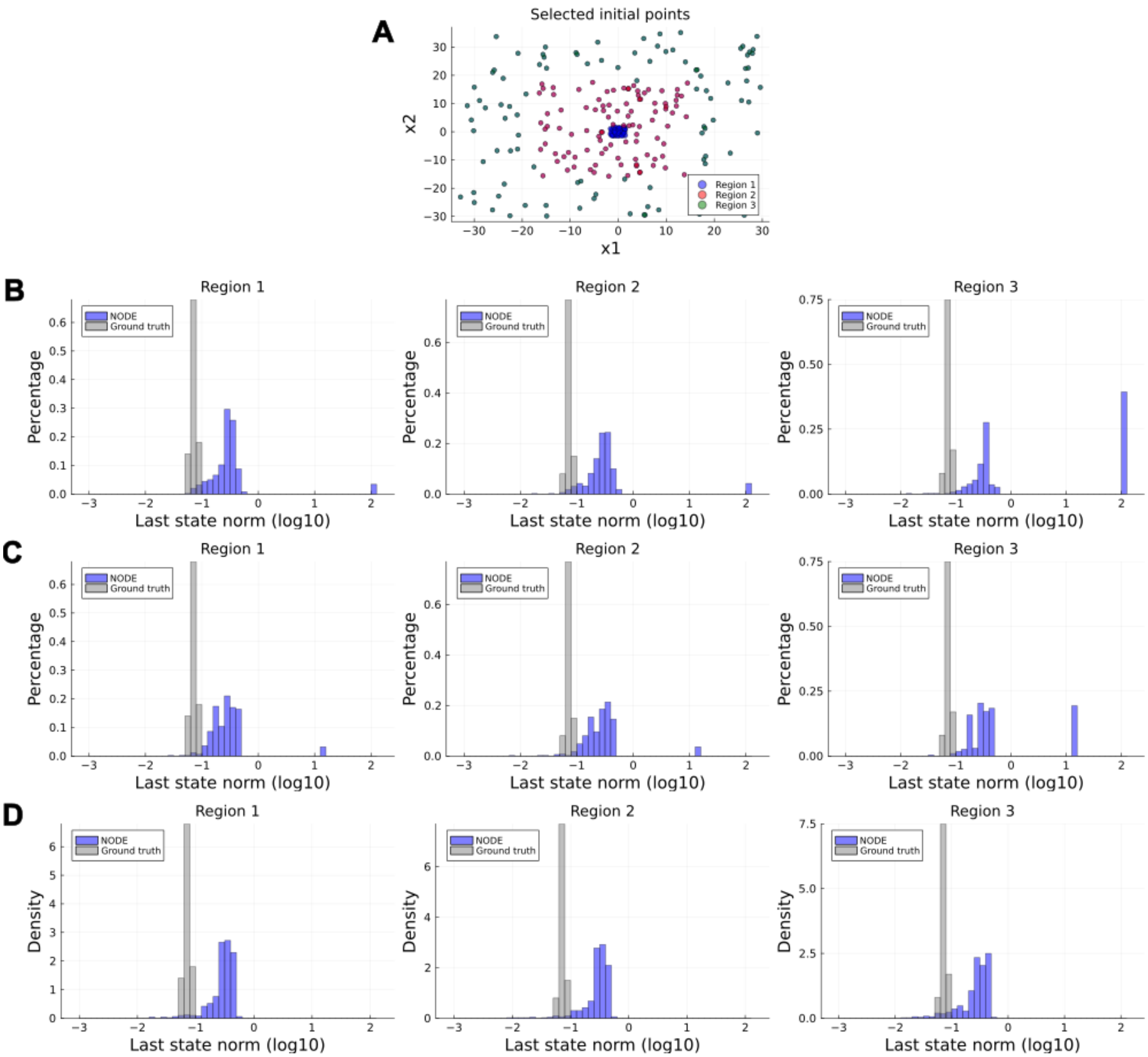


Figure S61: **Distribution of the norm of the last state of the simulations of the models composing the MOD ensembles, starting from different initial points.** Panel A shows the selected initial points, divided into three regions according to their distance from the training trajectories. Panels B, C, and D compare the last state of the simulations over the time interval $[0, 100]$ for the models in the MOD ensembles and the ground truth system, starting from points within the different regions. The comparison is made for the models trained in the purely data-driven scenario (Panel B), the partially data-driven with fixed mechanistic parameters scenario (Panel C), and the partially data-driven scenario (Panel D). For clarity, the norms have been clipped to $10^2$. In the UDE case, 53% and 60% of the trajectories were numerically unstable and diverge (norm exceeding $10^8$), and thus the related last state norm is not reported in the plot.

[GIP+18] Timur Garipov, Pavel Izmailov, Dmitrii Podoprikhin, Dmitry P Vetrov, and Andrew G Wilson. Loss surfaces, mode connectivity, and fast ensembling of dnns. *Advances in Neural Information Processing Systems*, 31, 2018.

[GRF+24] Stefano Giampiccolo, Federico Reali, Anna Fochesato, Giovanni Iacca, and Luca Marchetti. Robust parameter estimation and identifiability analysis with hybrid neural ordinary differential equations in computational biology. *NPJ Systems Biology and Applications*, 10(1):139, 2024.

[HOPY20] Hoon Hong, Alexey Ovchinnikov, Gleb Pogudin, and Chee Yap. Global identifiability of differential models. *Communications on Pure and Applied Mathematics*, 73(9):1831–1879, 2020.

[IKB21] Alexander Immer, Maciej Korzepa, and Matthias Bauer. Improving predictions of bayesian neural nets via local linearization. In *International conference on artificial intelligence and statistics*, pages 703–711. PMLR, 2021.

[KHH20] Agustinus Kristiadi, Matthias Hein, and Philipp Hennig. Being bayesian, even just a bit, fixes overconfidence in relu networks. In *International conference on machine learning*, pages 5436–5446. PMLR, 2020.

[LDM22] Nicholas N Lam, Paul D Docherty, and Rua Murray. Practical identifiability of parametrised models: A review of benefits and limitations of various approaches. *Mathematics and Computers in Simulation*, 199:202–216, 2022.

[Lin95] Bruce G. Lindsay. *Mixture Models: Theory, Geometry, and Applications*, volume 5 of *NSF-CBMS Regional Conference Series in Probability and Statistics.* Institute of Mathematical Statistics, Hayward, CA, 1995.

[LS03] Xin Liu and Yongzhao Shao. Asymptotics for likelihood ratio tests under loss of identifiability. *The Annals of Statistics*, 31(3):807–832, 2003.

[OTH23] Katharina Ott, Michael Tiemann, and Philipp Hennig. Uncertainty and structure in neural ordinary differential equations. *arXiv preprint arXiv:2305.13290*, 2023.

[QM09] Tom Quaiser and Martin Mönnigmann. Systematic identifiability testing for unambiguous mechanistic modeling–application to jak-stat, map kinase, and nf-$\kappa$ b signaling pathway models. *BMC systems biology*, 3(1):50, 2009.

[RMM+20] Christopher Rackauckas, Yingbo Ma, Julius Martensen, Collin Warner, Kirill Zubov, Rohit Supekar, Dominic Skinner, Ali Ramadhan, and Alan Edelman. Universal differential equations for scientific machine learning. *arXiv preprint arXiv:2001.04385*, 2020.

[SGJ+21] Berfin Simsek, François Ged, Arthur Jacot, Francesco Spadaro, Clément Hongler, Wulfram Gerstner, and Johanni Brea. Geometry of the loss landscape in overparameterized neural networks: Symmetries and invariances. In *International Conference on Machine Learning*, pages 9722–9732. PMLR, 2021.

[TJ21] Evren Mert Turan and Johannes Jäschke. Multiple shooting for training neural differential equations on time series. *IEEE Control Systems Letters*, 6:1897–1902, 2021.

[Vis04] Divakar Viswanath. The fractal property of the lorenz attractor. *Physica D: Nonlinear Phenomena*, 190(1-2):115–128, 2004.

[VRHB22] Alejandro F Villaverde, Elba Raimúndez, Jan Hasenauer, and Julio R Banga. Assessment of prediction uncertainty quantification methods in systems biology. *IEEE/ACM transactions on computational biology and bioinformatics*, 20(3):1725–1736, 2022.

[Wil79] Robert F Williams. The structure of lorenz attractors. *Publications Mathematiques de l'IHES*, 50:73–99, 1979.

[YLRK20] Alireza Yazdani, Lu Lu, Maziar Raissi, and George Em Karniadakis. Systems biology informed deep learning for inferring parameters and hidden dynamics. *PLoS computational biology*, 16(11):e1007575, 2020.
